# aureoRNA enables the interactive, genome-wide analysis of *Staphylococcus aureus* small RNA targetomes and uncovers new regulatory functions

**DOI:** 10.64898/2026.11.17.767300

**Authors:** Fayyaz Hussain, Maximilian P. Kohl, Shengwei Hou, Steffen Lott, Edith Ams, Karsten Voigt, David Lalaouna, Pascale Romby, Stefano Marzi, Isabelle Caldelari, Wolfgang R. Hess, Jens Georg

## Abstract

Small RNAs (sRNAs) control key steps of infection and stress acclimation in the opportunistic pathogen *Staphylococcus aureus*. Yet most of the more than 250 annotated candidates remain uncharacterized, and even the best-studied have incomplete target spectra. Here we present **aureoRNA**, an interactive tool that assembles for each sRNA comparative target predictions, available MAPS interactomes, sRNA–mRNA expression correlation across curated public transcriptomes and placement in groups of independently regulated genes (iModulons), and makes these data explorable in one place. GO-and KEGG-based enrichments summarize the datasets at the pathway level for every sRNA, and a weighted rank product combines available data into prioritized target candidate lists. We generated a new MAPS interactome for RsaD and experimentally validated predictions for three sRNAs: new RsaD targets in threonine and arginine catabolism, *citZ* as an RsaE target, and prophage genes repressed by RNAIII. aureoRNA is a hypothesis-generating resource for the systematic characterization of the sRNA repertoire of this pathogen.

## Introduction

Bacteria inhabit numerous ecological niches and experience rapidly varying environmental conditions. To understand the way bacteria sense, transmit and respond to external signals, and the mechanisms of dynamically regulating their metabolism and pathogenicity, genome-wide transcriptomic approaches have been applied, frequently detecting an unexpected complexity of the RNA landscape including multiple types of non-coding RNAs (Caldelari *et al*, 2013; Dar & Sorek, 2018; Westermann *et al*, 2012; Dove & Melamed, 2026). Non-coding RNAs include *trans*acting regulatory small non-coding RNAs (sRNAs), antisense RNAs (asRNAs) overlapping annotated genes, or cis-acting RNAs such as riboswitches, T-boxes or RNA thermometers, as well as structural RNAs, tRNAs and rRNAs.

*Staphylococcus aureus* is a pyogenic, toxigenic, mainly extracellular Gram-positive pathogen responsible for infections from minor skin abscesses to life-threatening diseases (Tong *et al*, 2015). Methicillin-resistant *S. aureus* (MRSA) is a leading cause of bacterial infections (Lee *et al*, 2018). In *S. aureus* and other studied bacteria, sRNAs have been found at the heart of regulatory pathways controlling responses to adverse conditions, population density, cell surface composition, metabolism during growth or differentiation, and virulence gene expression and immune escape (Bronesky *et al*, 2016; Rios-Delgado & Boyd, 2026; Menard *et al*, 2022). Therefore, it is of utmost significance to identify the regulatory targets of these sRNAs.

Most sRNAs target functionally related mRNAs. To facilitate genome-wide analyses of sRNA/mRNA interactions, specific approaches such as CLASH and RIL-seq have been developed which rely on the interaction of sRNAs/targets with proteins (Melamed *et al*, 2016; Waters *et al*, 2017). RIL-seq has expanded the ensemble of known sRNA targets in *E. coli* (Melamed et al, 2016), *Clostridioides difficile* (Fuchs et al, 2023), *Pseudomonas aeruginosa* (Gebhardt et al, 2023), *Salmonella enterica* (Matera et al, 2022), and *Vibrio cholerae* (Huber et al, 2022), and revealed the dynamics of regulatory networks under various stress conditions (Silverman & Melamed, 2025). These methods can identify many RNA-RNA interactions simultaneously. However, they cannot uncover RNA interactions without a cognate RNA-binding protein. Consequently, Direct Duplex Detection methods have been developed (for review, see (Schönberger *et al*, 2018)), which do not require any interacting protein. For example, LIGR-seq, originally developed for the global mapping of RNA-RNA interactions in humans (Sharma *et al*, 2016), has been applied to identify interacting RNAs in *Bacillus subtilis* (Durand *et al*, 2021), and in modified form in a cyanobacterium (Mandal *et al*, 2026). Although significant discoveries have been made using these methods, they lack sensitivity in detecting poorly expressed targets, distinguishing between direct and indirect targets, and struggle to identify sRNA targets with multiple pairings. Many of these issues were solved by tagging the sRNA of interest with an RNA aptamer suitable for affinity chromatography, such as in the MS2-affinity purification coupled with sequencing (MAPS), in which sRNAs and interacting RNAs are fished together and sequenced (Carrier *et al*, 2016; Lalaouna & Massé, 2015). MAPS can also find sRNA targets that form multiple pairings through distant mRNA regions (Lalaouna *et al*, 2017, 2018). MAPS was the key technique in *S. aureus* in identifying mRNAs encoding the transcription factor IcaR, the *glcU_2* glucose transporter, and the sRNA RsaG as targets of the sRNA RsaI (Bronesky *et al*, 2019), in revealing the *in vivo* interaction network of the iron starvation-relevant sRNA IsrR (Rios-Delgado *et al*, 2025), and many more. Nevertheless, despite the impressive progress in this field, the experimental identification of sRNA targets and functions has remained tedious. Therefore, the combination of large-scale experimental methods with alternative strategies based on computational predictions is attractive.

Computational approaches such as CopraRNA integrate phylogenetic data and structure information to predict sRNA regulatory domains and targets at genome-wide scale, perform functional enrichment, and reconstruct the regulatory networks (Wright *et al*, 2013, 2014). The predictive power of such approaches has been further improved by taking the globally available suite of putative sRNA homologs in different bacteria into account. Independent studies revealed that CopraRNA currently provides the best strategy to predict sRNA targets and to classify the sRNAs according to their functions in different bacteria (Pain *et al*, 2015; Naskulwar & Peña-Castillo, 2022). However, prediction of sRNA targets at the genome-wide level was originally less efficient in low GC Gram-positive bacteria, such as *S. aureus*, most likely due to subtle differences in base pairing formation compared to the better studied Enterobacteriaceae. Moreover, the more restricted conservation of *S. aureus* sRNAs and their targets could have been problematic. Finally, no RNA chaperone proteins have so far been identified as participating in sRNA regulation in *S. aureus,* limiting the use of methods such as RIL-seq. Considering the situation in *S. aureus*, we concluded that a combination of experimental and computational approaches provides the maximum of reliable information for the targetome of a bacterial sRNA and improved the algorithmic approach with the current version of CopraRNA (Georg *et al*, 2020).

Previously, hundreds of putative sRNAs were reported for *S. aureus* (Mäder *et al*, 2016; Romilly *et al*, 2012; Tomasini *et al*, 2014; Guillet *et al*, 2013). On the other hand, when 5’ and 3’ UTR-derived sRNAs were excluded, only a subset of about 50 were proposed to represent true sRNAs (Liu *et al*, 2018). Indeed, besides sRNAs, bacterial genomes also contain open reading frames in unusual places, in-frame as well as out-of-frame within longer coding sections, in cis-asRNAs or in intergenic regions previously considered too short for gene calling (Orr *et al*, 2020). These frequently short ORFs (sORFs) might encode smaller proteins, often with a regulatory function, but systematic evidence from ribosome profiling has only recently been obtained for *S. aureus* (Kohl *et al*, 2026). Consequently, there is an increasing number of sRNAs, which are recognized to also encode peptides, or to represent bona-fide mRNAs. In *S. aureus*, the quorum-sensing induced RNAIII, which forms base pairings with multiple mRNAs encoding transcriptional regulators and virulence factors, also encodes a phenol-soluble modulin (PSM) peptide with hemolytic activity (Hld) (Bronesky *et al*, 2016; Novick & Geisinger, 2008).

It can be very challenging to obtain proper sRNA-related information given their heterogeneity, the fact that there are potentially hundreds of different sRNAs in any given bacterium and that for the genus *Staphylococcus* alone, as of today (May 03, 2026), 143,415 completed and draft status genome sequences are available (https://www.ncbi.nlm.nih.gov/genome/browse#!/prokaryotes/Staphylococcus). Therefore, dedicated databases can be of substantial value. As the first sRNA database centered on a single genus, previously the ‘Staphylococcal Regulatory RNA Database’ (SRD) was developed for *S. aureus* (http://srd.genouest.org/) (Sassi *et al*, 2015). The SRD focuses on sRNAs in *S. aureus* strain N315 as a reference providing a list of 159 independently transcribed likely instances of sRNAs, complemented by comparisons against homologs within altogether 32 *Staphylococcus* genomes. For each of these sRNAs, the SRD connects to the respective literature allowing the user to assess the respective validity and significance. In contrast, here we present a database, integrating experimental data and computational methods for curating novel *S. aureus* sRNA target predictions for the first time. Since we have performed extensive MAPS-and omics analyses, including high-resolution ribosome profiling for translation start site mapping in *S. aureus* HG001, we focused our computational approach on this strain with abundant complementary experimental data. HG001 is a derivative of strain NCT8325, containing a repaired *rbsU* gene encoding an activator of SigB (Caldelari *et al*, 2017; Herbert et al., 2010). Following a previously drafted strategy (Wright & Georg, 2018), we implemented a comprehensive bioinformatic workflow for sRNA characterization based on the full set of sRNA candidates identified in the course of the genome (Caldelari *et al*, 2017) and several transcriptome analyses (Geiger *et al*, 2012; Mäder *et al*, 2016).

In the present study, we searched systematically for all known and putative sRNAs and their likely homologs in the Bacillota (Firmicutes) phylum using GLASSgo (Lott et al, 2018). We identified those sRNAs that form conserved families and present their shared features with regard to sequence, predicted secondary structure and synteny. Next, we analyzed the predicted targetome with CopraRNA (Wright et al, 2013) and combined it with experimental MAPS data for IsrR (Rios-Delgado et al, 2025), RNAIII (Hallier et al, 2024), RsaA (Tomasini et al, 2017), RsaC (Lalaouna et al, 2019), RsaD (this study), RsaI/RsaOG (Bronesky et al, 2019), RsaG (Desgranges et al, 2022), and ZinS/RsaX20 (Charbonnier *et al*, 2026). We also examined gene expression correlations across a wide range of publicly available *S. aureus* transcriptome datasets. The results are available as an interactive online resource (https://aurorna.biologie.uni-freiburg.de/) to facilitate the characterization of sRNAs in *S. aureus*.

## Results and Discussion

### The aureoRNA workflow

The genome of *S. aureus* HG001 contains 61 annotated tRNAs, 15 rRNAs, 1 transfer-messenger RNA (tmRNA), and 59 other annotated non-coding RNAs (Caldelari *et al*, 2017). The SRD lists 418 non-coding RNAs including 235 putative sRNAs which were experimentally validated (http://srd.genouest.org/browsevalidated) in *S. aureus*, while the RFAM database lists 88 confirmed and characterized sRNA entries for *S. aureus* sp. *aureus* NCTC 8325, which possesses the closest genome sequence to the *S. aureus* strain HG001. To consider the widest possible spectrum, here we started with the analysis of the entire set of listed sRNAs. In several instances, we could join overlapping fragments of separately annotated sRNAs, for example the locus tags HG001_00389, HG001_00390, HG001_03236 and HG001_00391 were joined as these belong to the 4.5S RNA. We also grouped sets of identical sRNAs into families, which contain up to four identical members. As a result, the initial complexity was reduced by about half to 252 putative sRNAs.

The data generation workflow (**Supplementary Figure S1**) started with a homology search using GLASSgo (Lott et al, 2018) against the NCBI database, restricted to the phylum Bacillota. The found hits were complemented by members of the respective Rfam families. Using Taxonkit (Shen & Ren, 2021), we analyzed the taxonomic distribution of sRNA homologs exhibiting ≥ 57% sequence identity to the *S. aureus* HG001 reference. These distributions are visualized in **Figure 1** for the genus *Staphylococcus*. The whole distribution data can be viewed on the aureoRNA website (**Supplementary Figure S2**). The identification of these homologs enabled a comparative genomics workflow to construct consensus secondary structures, analyze synteny and promoter conservation, and predict *trans*-acting sRNA targets using CopraRNA (Wright et al, 2013). A primary advantage of aureoRNA is the integration of these predictions with public GEO transcriptome datasets, MAPS interactome data, and functional enrichments (GO, KEGG, UniProt, and metabolites) into a unified “targetome” proposal. These lists can be mapped to KEGG metabolic pathways (Kanehisa et al, 2025) and genome-scale networks (Renz & Dräger, 2021) for further investigation. The resulting platform (https://aurorna.biologie.uni-freiburg.de/) serves as a dynamic environment for filtering and re-analyzing data to generate testable hypotheses for downstream experiments.

**Figure 1.**
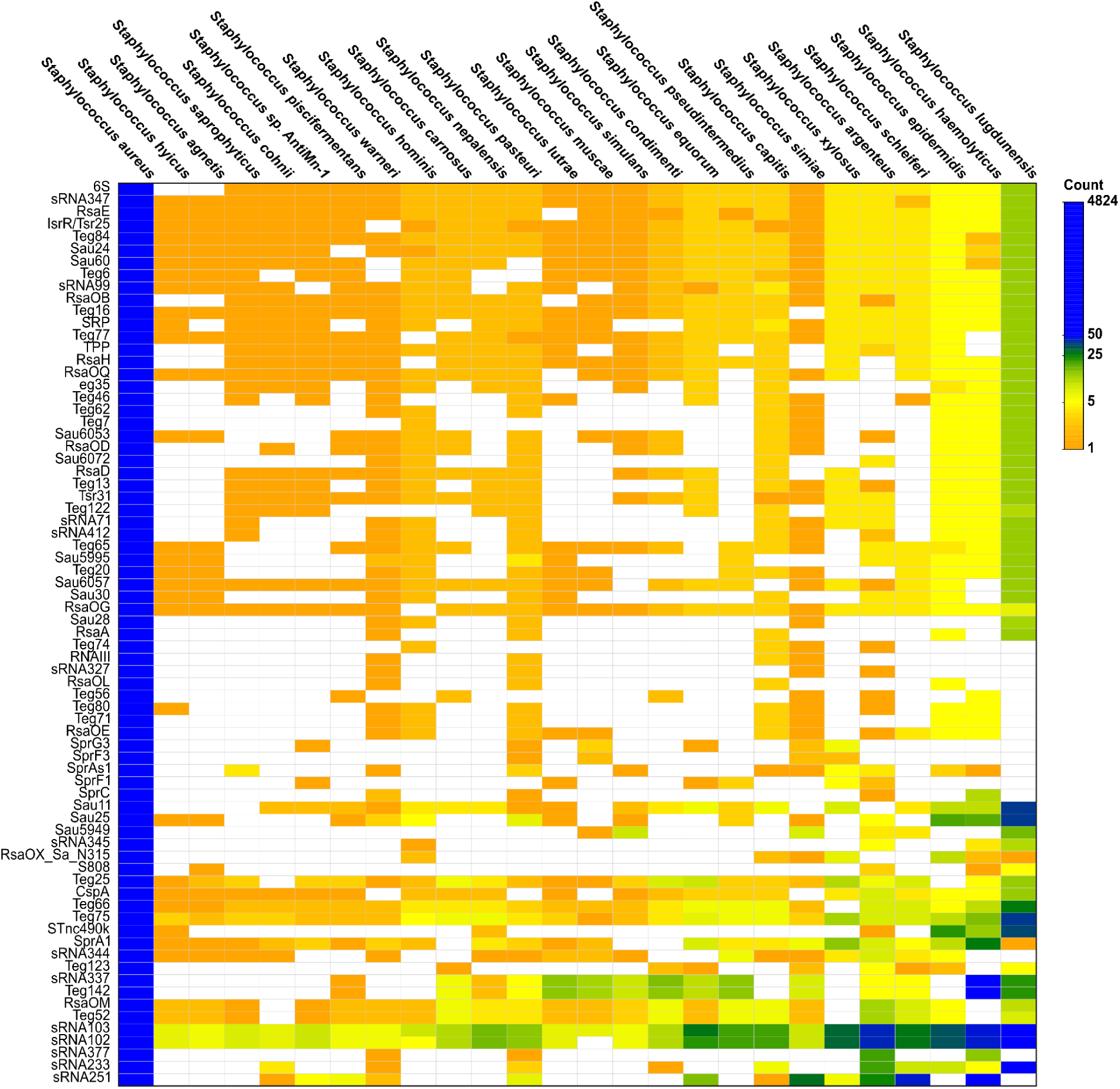
Taxonomic representation of all sRNAs present in at least 5 different species of the genus *Staphylococcus*. The color code indicates the number of homologs detected in the respective species and the corresponding lower taxonomic levels. Only homologs with an overall sequence identity ≥ 57% to the *S. aureus* HG001 query sequence were considered. The respective aureoRNA database allows for a full overview as well as diverse filter and selection options. See **Supplementary Figure S2** for detailed explanations.

### AureoRNA usage: sRNA discovery by search

sRNAs of interest can be searched through diverse strategies (**Supplementary Figure S3**). A direct search by the sRNA name, diverse aliases or the locus-tag is complemented by explorative analyses. In addition, a target-centric search can be used to identify putative sRNA regulators for a specific mRNA of interest. Here, we present examples for three mRNAs that raised our attention based on recent ribosome profiling (Ribo-Seq) data (Kohl *et al*, 2026).

A target-centric search was performed to identify sRNAs containing the query mRNA within the top 100 predictions for *citZ* mRNA, which encodes citrate synthase, and in some *S. aureus* strains carries a non-canonical AUA start codon (**Figure 2**; (Kohl *et al*, 2026)). Among the predicted regulators, RsaE emerged as the second ranked candidate by p-value in the inverse CopraRNA search, while *citZ* appeared within the top predictions for RsaE (**Supplementary Figure S4A**). Regulation of *citZ* by RoxS (homolog of RsaE in *B. subtilis*) has been previously proposed (Allouche *et al*, 2023), but the actual interaction in *S. aureus* has not been experimentally verified thus far. Inspection of the predicted interaction revealed base pairing overlapping the Shine-Dalgarno region and start codon of *citZ* (**Figure 2A**), suggesting a mechanism of translational repression. Toeprinting assays showed that RsaE prevented ribosome recruitment to the *citZ* mRNA (**Figure 2D**). Notably, the inhibition of ribosome binding was more pronounced for constructs carrying the native AUA start codon compared to an AUG variant despite the fact that the base pairing scheme did not change. This observation indicates that RsaE-mediated regulation is particularly effective at weak translation initiation sites. These results demonstrate that aureoRNA-guided predictions can uncover regulatory interactions with direct mechanistic and physiological relevance.

**Figure 2.**
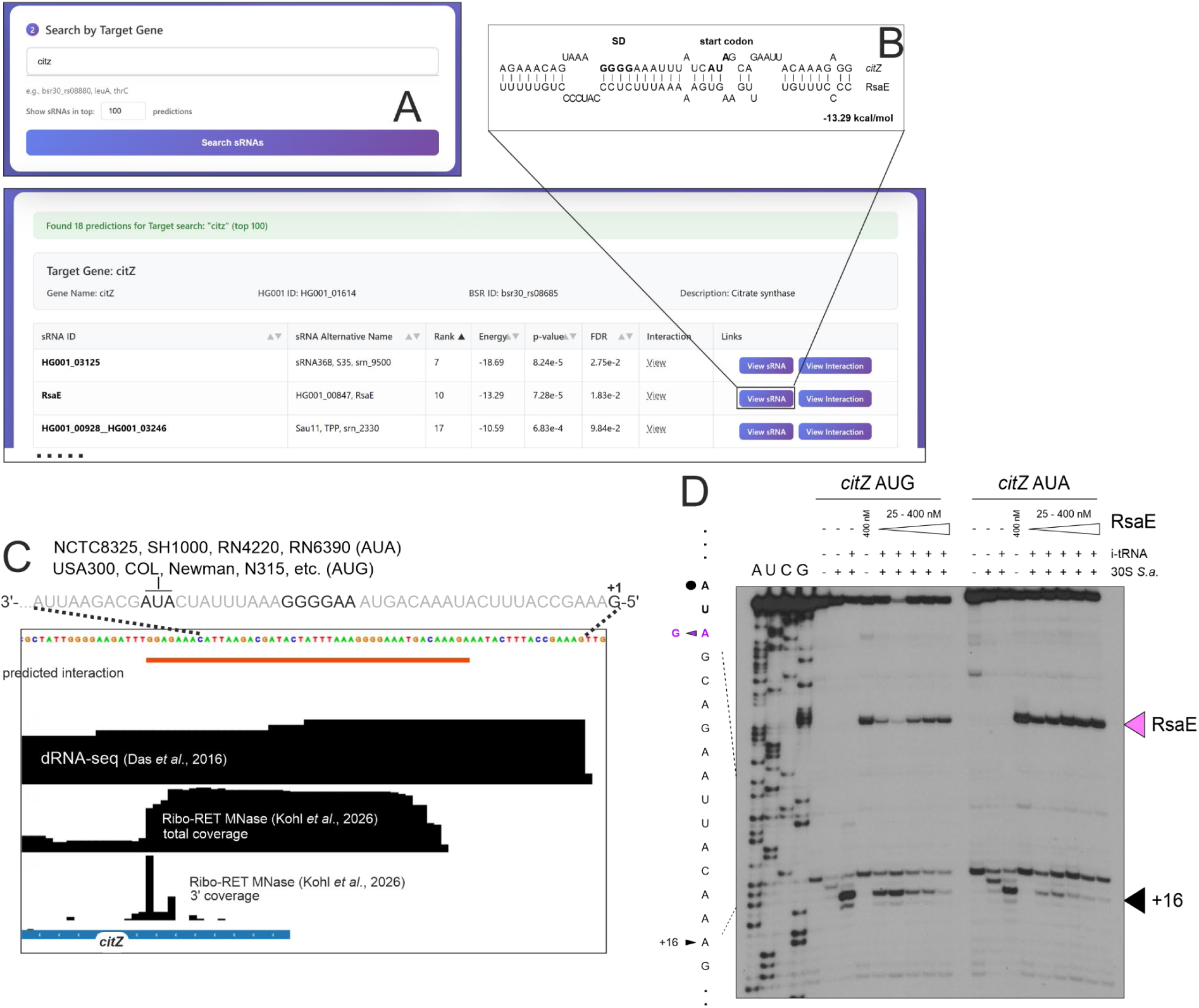
Interaction of *citZ* mRNA with RsaE as detected by the aureoRNA reverse search. **(A)** Screenshot showing the search slot and the first 3/19 search results with *citZ* within the top 100 CopraRNA predictions. More information is found in **Supplementary Figure S4**. **(B)** Predicted interaction region of *citZ* and RsaE. **(C)** Modified screenshot from the aureoRNA website showing the genomic region of *citZ* in an IGV browser panel which is a part of each sRNA’s landing page. The transcriptome coverage from dRNA-seq data (Das et al, 2016) and the Ribo-Ret data (full coverage and 3’ coverage) are shown for reference. **(D)** Toeprinting assays with RsaE and *citZ* mRNA variants. A simplified initiation complex was formed with *citZ* mRNA, *S. aureus* 30S and the initiator tRNA_f_^Met^ (i-tRNA) in the absence or presence of increasing concentrations of RsaE (25, 50, 100, 200, 400 nM). Control experiments were performed on the mRNA alone, and in the presence of either 30S or RsaE (400 nM). Lanes A, U, C, G are sequencing ladders. The toeprint at position +16 and the stop induced by RsaE binding are shown on the right side of the autoradiography.

We next applied the same target-centric strategy to additional candidates identified from recent Ribo-seq data (Kohl *et al*, 2026), focusing on two highly similar sORFs located upstream of the prophage holin (*HG001_01353*) and endolysin (*HG001_01849*) genes (**Figure 3A, B**). The reverse search identified 18 candidate sRNA regulators within the top 100 predictions (**Supplementary Figure S4B**). From this ranked list, RNAIII (rank 7) was selected for further analysis, as the predicted interaction site within its H14 domain targets the Shine–Dalgarno sequences of both sORFs, which are identical. RNAIII is the main sRNA effector of the *S. aureus* quorum sensing system, displaying increased expression levels with higher cell density in the exponential growth phase. A direct interaction of RNAIII linked to the regulation of prophage genes is of interest, because it could provide the phage with a cue for surrounding cell density. Such a cue could allow the phage to adjust its development of productive phage particles, according to the availability of potential hosts following lysis, as is the case for other quorum-sensing phages (León-Félix & Villicaña, 2021; Sprenger *et al*, 2024). Toeprinting assays showed that RNAIII binding prevented the formation of the ribosomal initiation complex, supporting the hypothesis that RNAIII acts as a translational repressor of the two small prophage proteins (**Figure 3C, D**). Moreover, deletion analysis of RNAIII confirmed that both binding and regulatory activity depend on the H14 domain, consistent with the predicted interaction site (**Figure 3E, F**).

**Figure 3.**
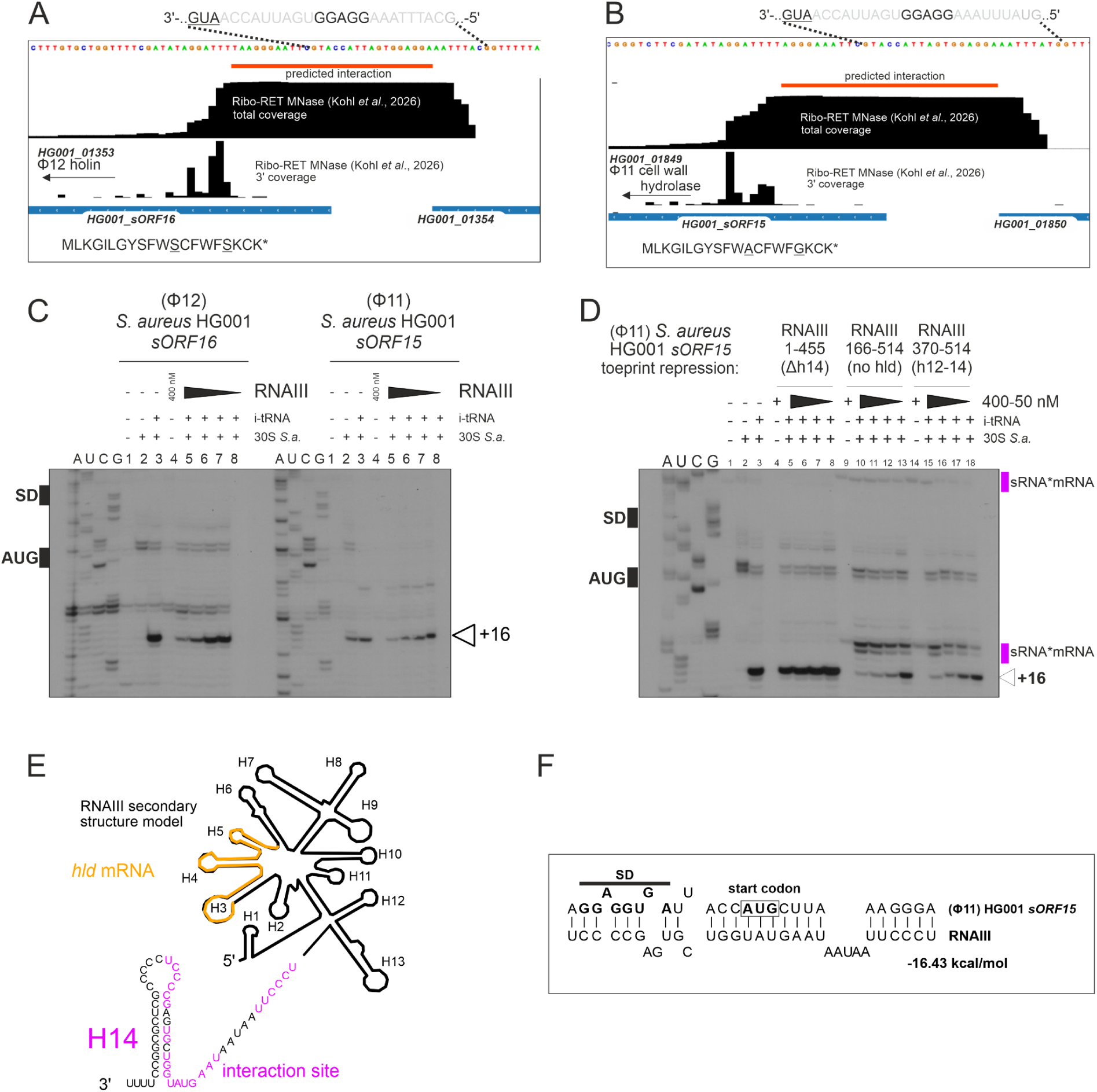
Interaction of phage sORFs with RNAIII as detected by the aureoRNA reverse target search. (A, **B)** Modified screenshot from the aureoRNA website showing the genomic region of *HG001_sORF16* upstream of *HG001_01353* (encoding a phage holin) and *HG001_sORF15* upstream of *HG001_01849* (encoding a phage cell wall hydrolase) in IGV browser panels, respectively. The coverage from Ribo-Ret data (full coverage and 3’ coverage) are shown for reference. **(C)** Toeprinting analysis of RNAIII-mediated inhibition of initiation complex formation on the *S. aureus* HG001 sORF16 (ϕ12) and sORF15 (ϕ11) mRNAs. Simplified initiation complexes were assembled with mRNA, *S. aureus* 30S ribosomal subunits and initiator tRNAᶠᴹᵉᵗ (i-tRNA), in the absence or presence of RNAIII (50, 100, 200 or 400 nM). Controls contained mRNA alone or mRNA supplemented with either 30S subunits or RNAIII (400 nM). **(D)** Toeprinting analysis showing that RNAIII hairpin h14 is required for inhibition of initiation complex formation on *HG001_sORF15* mRNA. RNAIII truncation variants comprised nucleotides 1-455, lacking h14 (Δh14); nucleotides 166-514, lacking the *hld* coding region (no *hld*); or nucleotides 370-514, corresponding to the 3′ domain containing hairpins h12-h14. Each variant was tested at decreasing concentrations from left to right (400-50 nM). Pink bars mark reverse-transcription stops induced by RNAIII binding (sRNA*mRNA). In both panels, open arrowheads indicate toeprints at position +16 relative to the first nucleotide of the AUG start codon. Shine-Dalgarno (SD) sequences and start codons are annotated beside the gels. A, U, C and G denote sequencing ladders. **(E)** Scheme of RNAIII secondary structure model with relevant hairpins labelled. The nucleotides involved in the interaction site are colored in pink. The *hld* ORF is shown in yellow. **(F)** The predicted interaction of RNAIII and *HG001_sORF15,* covering the Shine-Dalgarno (SD) motif and start codon.

A third search option allows to investigate if a specific KEGG pathway (Kanehisa et al, 2025), GO term (The Gene Ontology Consortium, 2026), UniProt keyword (The UniProt Consortium, 2025), or a gene set associated with a specific metabolite (based on an *S. aureus* NCTC 8325 genome-scale metabolic model) (Renz & Dräger, 2021) is potentially regulated by an sRNA (**Supplementary Figure S5**). **Figure 4** shows examples for sRNAs predicted to regulate the citrate cycle (TCA), while **Supplementary Figure S6** shows predictions for lipid biosynthesis associated targets or targets that are involved in reactions with the metabolite acetyl-CoA.

**Figure 4.**
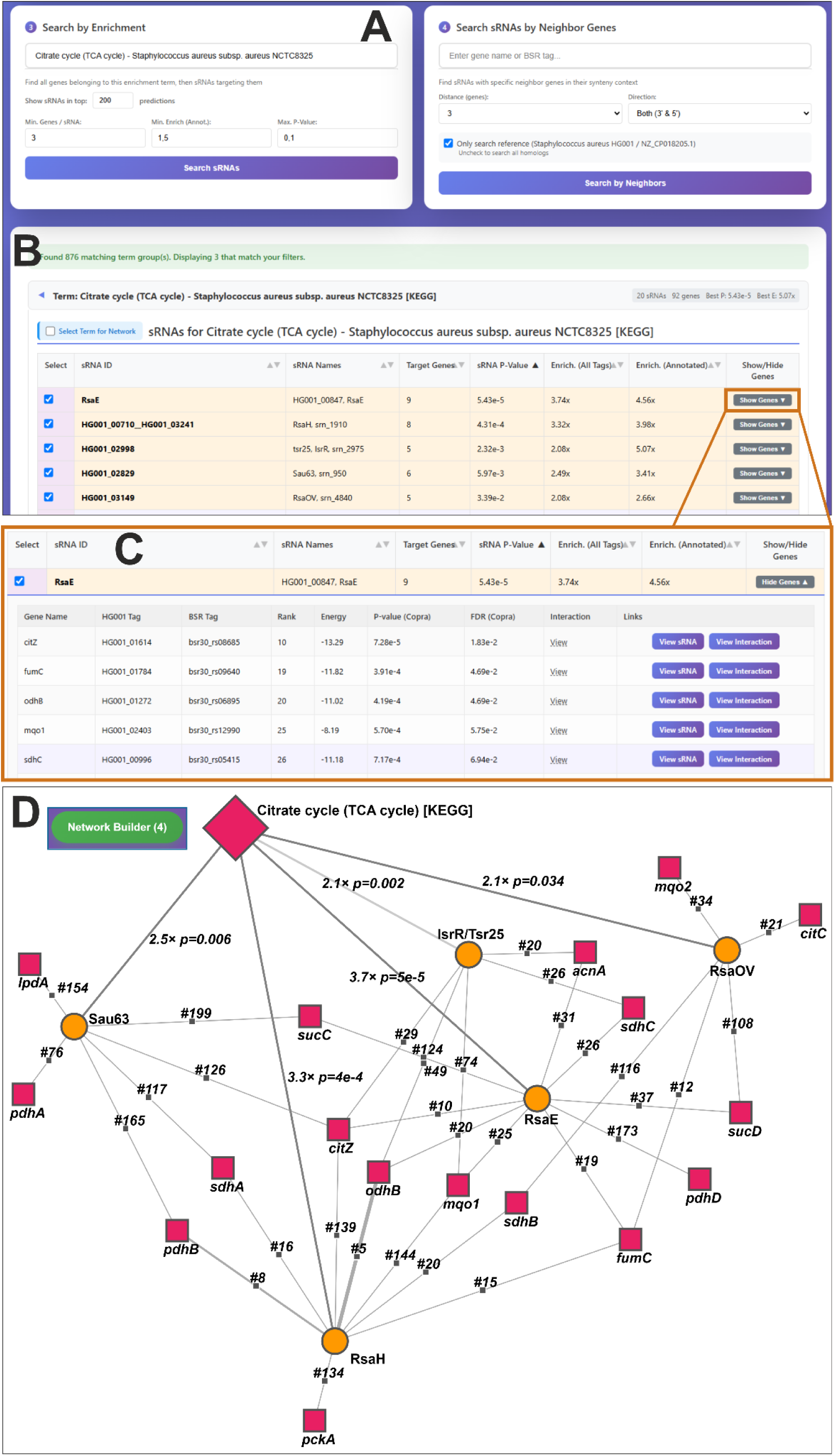
Example for an enrichment-centric explorative sRNA search. **(A)** The term “Citrate cycle (TCA cycle)” is searched with a CopraRNA prediction rank threshold of 200. AureoRNA performs a functional enrichment within all top200 predictions of all included sRNAs for all annotation terms that contain the string “Citrate cycle (TCA cycle)”. The CopraRNA rank threshold, the minimal fold-enrichment, the maximal p-value and the minimal member count of a term are adjustable parameters. Annotations that are enriched include GO terms, UniProt key words and KEGG pathways for *S. aureus* NCTC 8325 as well as metabolites from the KEGG pathways and a genome wide metabolic model. **(B)** The term “Citrate cycle (TCA cycle)” is enriched in 20 sRNAs; the top five (enrichment p-value sorted) are displayed here. **(C)** Clicking on the “show genes” button expands the list of enriched genes for the selected sRNA with links to the sRNA landing page and the interaction conservation page. 5/9 predicted RsaE TCA-cycle targets are shown. **(D)** Found sRNAs can be selected by checking the select box to be transferred to the aureoRNA “network builder”. Multiple sRNAs from different enrichment searches can be added. The example shows the top five sRNAs predicted to regulate TCA-cycle genes, as determined by CopraRNA predictions and UniProt keyword searches. Prediction ranks, term enrichment fold-changes and p-values are written at the respective edges. The aureoRNA network viewing tool (**Supplementary Figure S5**) is based on a Cytoscape plugin (Franz et al, 2023).

### AureoRNA usage: sRNA discovery by expression profiles

AureoRNA taps the wealth of *S. aureus* transcriptome datasets (RNA-seq & microarray) to improve the sRNA target prediction by assessing the expression correlation of sRNAs and potential targets (Modi *et al*, 2011; Derinkok *et al*, 2025; de Carvalho *et al*, 2026). The sRNA expression patterns can be investigated over a broad range of conditions and mutant strains. To maximize coverage, we incorporated data from various *S. aureus subsp. aureus* strains (NCTC8325, N315, USA300, UAMS-1, Newman, 15981, and environmental samples) and their sub-strains. All data were mapped to the HG001 genome, a derivative of strain NCTC8325, to ensure consistency across the dataset. The main gene expression analysis resource is a clustered fold-change heatmap which can be filtered and subsetted dynamically (**Figure 5A**). Given the technical heterogeneity of the samples, encompassing different strains, diverse methods such as microarray and RNA-seq, varying library preparations, and different sequencing platforms, we utilized fold-changes relative to study-specific controls rather than globally normalized expression values to facilitate cross-study comparison. It should be noted that especially smaller sRNAs may be underrepresented in some datasets due to size limitations in the experimental workflows. In order to place the sRNAs into groups of similar regulated genes or regulons, the RNA-seq data are further used to define independently modulated groups of genes (iModulons) (Sastry et al, 2019; Poudel et al, 2020). This yielded 117 robust iModulons of which 109 are retained after filtering accounting for 66.6% of the data variance placing 176 sRNAs into 98 modules. From these modules, 75 of them (58.2% variance) contain biologically interpretable content. The database can be used to address the following questions:

1. Which genes are co-expressed with the query sRNA or gene, based on correlation or on the cophenetic distance within the clustered expression data? The most correlated genes can also be analyzed by functional enrichment to identify similarly regulated GO-terms or pathways.
2. To which iModulon, i.e. potentially regulon, does an sRNA or gene belong to?
3. Under which conditions does the expression of the sRNA change? In other words, when does the sRNA, or the corresponding iModulon, become relevant?
4. Which sRNAs are up-or down-regulated in a given condition?

**Figure 5.**
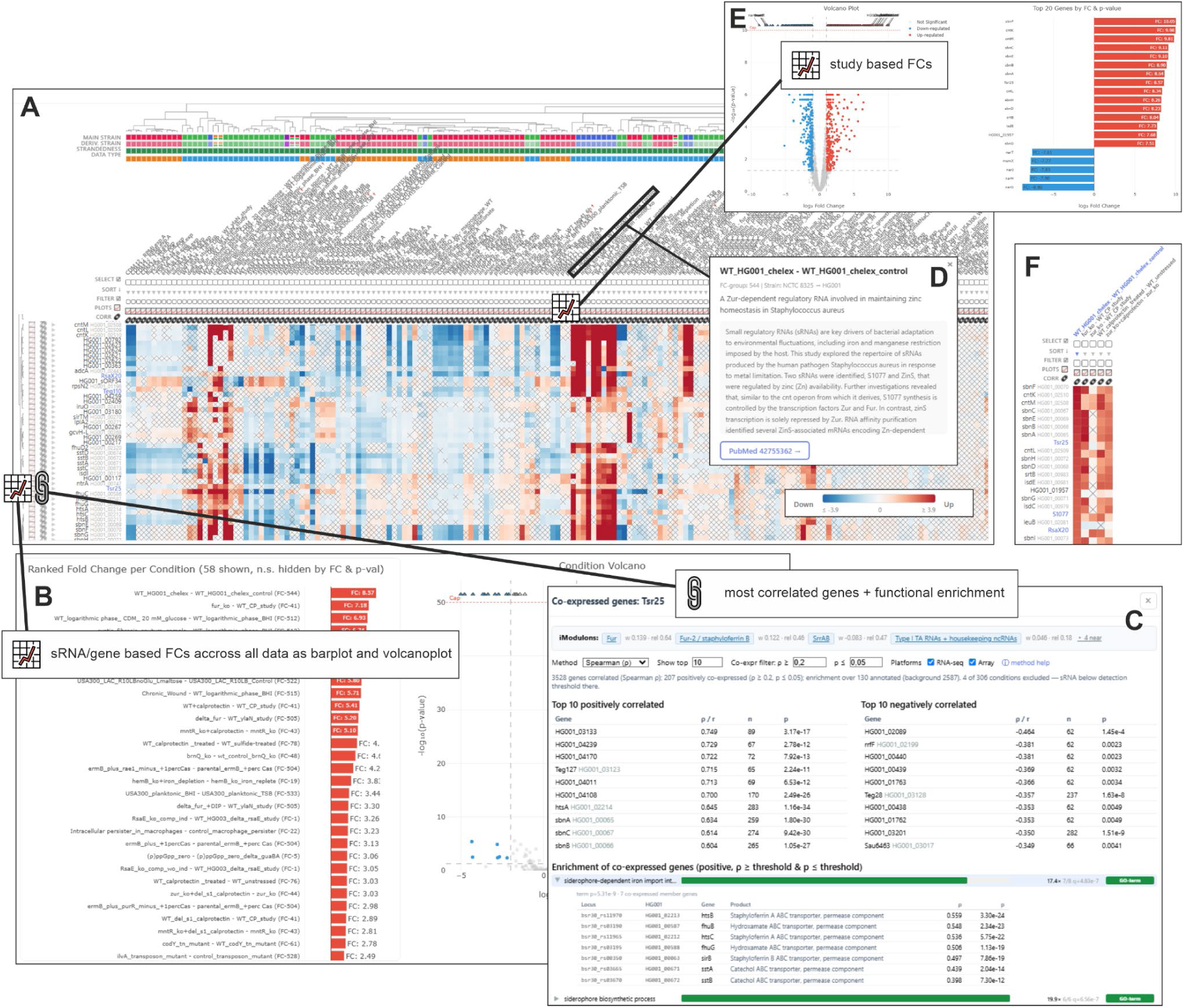
Expression pattern page with examples focused on IsrR/Tsr25. Modified aureoRNA screenshot, for a full explanation, please use the “guided tour” on the website. **(A)** Heatmap of expression changes across >300 fold-change comparisons from, currently 62 RNA-seq and microarray transcriptome studies in different *S. aureus* substrains as indicated in the annotation block above the heatmap. The heatmap shows either only the included 252 sRNAs or all *S. aureus* HG001 genes. Rows and columns are organized using hierarchical clustering using Ward’s true minimum variance method on standardized fold changes in R (hclust, ward.D2). The current view is centered around IsrR and its 20 closest context rows based on cophenetic distance. Non-significant FCs (default: p-Values > 0.05, adjustable) are indicated by a black cross. **(B)** Clicking on the chart symbol beneath each row name provides a popup with a bar chart-or volcano plot for the respective gene across all FCs. The heatmap could also be ordered by a specific FC comparison (column) and non-significant rows can be filtered. **(C)** A click on the link-clamp at the start of a row will open a window showing the most correlated and anti-correlated genes to the selected gene as well as the iModulons a sRNA/gene is assigned to. In addition, there is a functional enrichment for the most correlated genes. **(D)** Clicking on the FC title (column name) opens a pop-up containing the respective study title, abstract and PubMed link. **(E)** Clicking on the comparison (column) chart symbol opens the bar chart or volcano plots for the selected FC column. **(F)** Examples for an explorative search for interesting sRNAs based on their expression profiles showing the most up-regulated genes within Chelex metal depletion data (Charbonnier *et al*, 2026) and other metal stress data (Reyes Ruiz *et al*, 2024). The columns used for sorting are indicated with a blue triangle.

Below, we describe an example for an sRNA-centric workflow. Studying the metal depletion response in *S. aureus* HG001 revealed IsrR, ZinS and S1077 as the most strongly induced sRNAs (Charbonnier *et al*, 2026). In *S. aureus*, Fe, Mn, and Zn homeostasis are all controlled by a tandem of a transcription factor and an sRNA: FurIsrR (Rios-Delgado et al, 2025; Rios-Delgado & Boyd, 2026; Coronel-Tellez et al, 2022), MntR-RsaC (Lalaouna et al, 2019) and Zur-ZinS (Charbonnier et al, 2026). The highest-weighted iModulon memberships for IsrR, ZinS and S1077 reproduce this regulator assignment, “Zur” for ZinS and S1077, and “Fur” for IsrR and S1077, without any prior knowledge of the experiment.

Filtering the expression data for conditions in which the “Zur” iModulon is differentially activated (**Figure 6**) reveals, besides metal depletion studies, infection relevant conditions such as chronic wounds, infected macrophages and mouse liver infection, consistent with the sRNAs identified in human chronic wounds and cystic fibrosis sputum by a recent pangenome-based framework (Briaud *et al*, 2026). Fur, Zur and MntR control primarily metal transport systems, while the sRNAs contribute to metal-sparing by suppressing the synthesis of metal-utilizing proteins (Charbonnier et al, 2026; Rios-Delgado & Boyd, 2026; Lalaouna et al., 2019). To limit bacterial proliferation, host cells restrict the availability of metals, a defense mechanism known as nutritional immunity. It is therefore tempting to propose that the sRNA-mediated metal-sparing response is part of a major defense mechanism, particularly during chronic infection (Briaud et al, 2026; McFarlane et al., 2025).

**Figure 6.**
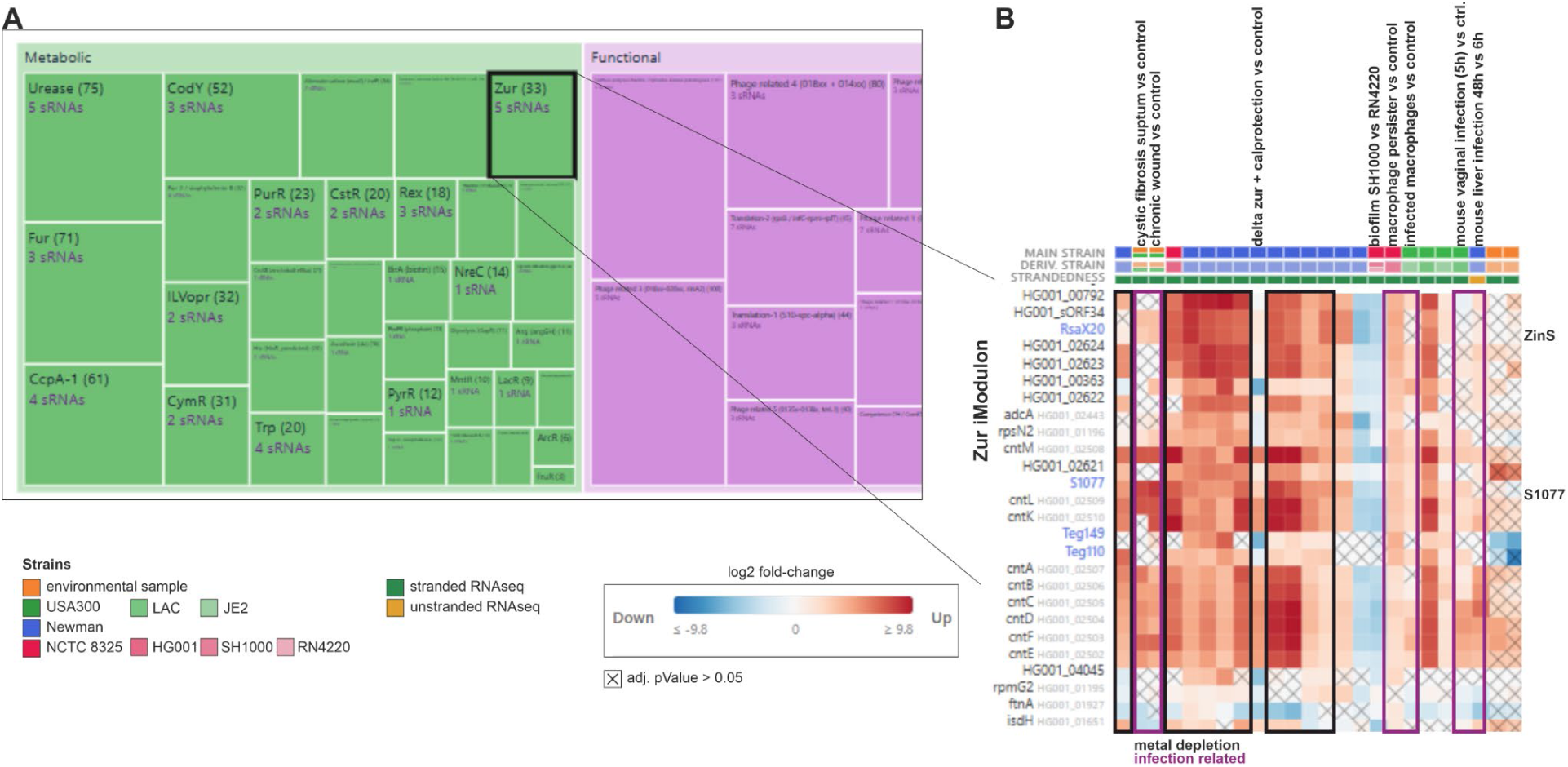
iModulon based expression analysis (modified screenshots). **(A)** Schematic view of the iModulon treemap. Each rectangle is one iModulon, labelled with its name, gene count and number of sRNA members, the area is proportional to gene count and color denotes the functional category (here Metabolic and Functional). **(B)** The Zur module is selected and the genes belonging to the module are shown, ordered by membership weight from high to low. Only fold-change columns in which the Zur iModulon activity changed by |Δ| ≥ 6 are displayed without a statistical significance threshold (RNA-seq only). Cell color gives the log_2_ fold change relative to the study-specific control (red, up; blue, down); crossed cells did not pass the significance threshold (i.e. adj. p-value > 0.05). The tracks above the heatmap show the main strain and the derivative strain in lighter shades of the main-strain color. For example, the main strain is NCTC 8325 and the derivative strain is HG001. Furthermore, it is indicated for each contrast column whether the respective libraries were stranded (green box) or unstranded (yellow box). Metal depletion and infection-related columns are framed in black and purple, respectively.

As an alternative way of identifying the relevance of an sRNA, the strongest expression FCs could be displayed as volcano or bar plot. In the case of IsrR, metal depletion conditions dominate the volcano plot (**Figure 5B**), while genes that are most strongly correlated with its expression are enriched for terms such as “siderophore-dependent iron import into cell” (**Figure 5C**). An unbiased, data-driven entry point, starting from a dataset rather than from a candidate sRNA is also possible. Sorting the Chelex metal depletion data (Charbonnier *et al*, 2026) by fold change puts IsrR/Tsr25, S1077 and ZinS/RsaX20 at rank 7, 17 and 19 of the most strongly induced genes (**Figure 5D-F**). In conclusion, the expression patterns of sRNAs can be investigated over a broad range of conditions and mutant strains. Using the IGV viewer plugin on the sRNA landing pages, it is moreover possible to view the transcriptome coverage for currently 367 different conditions genome-wide (**Supplementary Figure S7**).

Once a candidate sRNA is identified by one of the above-described methods, the dedicated landing page serves as a centralized repository for all associated resources and data (**Supplementary Figure S7**). This allows for an in-depth characterization of the sRNA including synteny (**Supplementary Figure S8**), (consensus) secondary structure, phylogenetic distribution (**Supplementary Figures S9-10**), as well as promoter and sequence conservation (**Supplementary Figure S10**). In the following, the whole process for exploratory analysis with aureoRNA is exemplified for investigating the targetome of a comparatively less well-characterized sRNA, RsaD. Furthermore, we detail the actual usage of all components via online “guided tours” on the aureoRNA website.

### The RsaD example: Synteny, promoter and signaling pathway

We first addressed the conservation and expression profiles of RsaD and detected in total 266 putative homologs for RsaD which were condensed into 31 representative sequences. A phylogenetic analysis of these 31 representative sRNAs delineates three major groups (**Figure 7A**). Such a tree is available for all sRNAs on their synteny pages (**Supplementary Figure S9**) or promoter/sequence consensus pages (**Supplementary Figure S10**). The database provides expression patterns as bar charts (**Figure 7C, Supplementary Figure S12**) or volcano plots for the individual transcripts. By inspecting this information, the user can analyze at which conditions a regulatory sRNA, here RsaD, might be relevant. In combination with the provided sequence logos of the sRNA promoter consensus (**Figure 7B**), potentially, relevant transcription factors can be identified.

**Figure 7.**
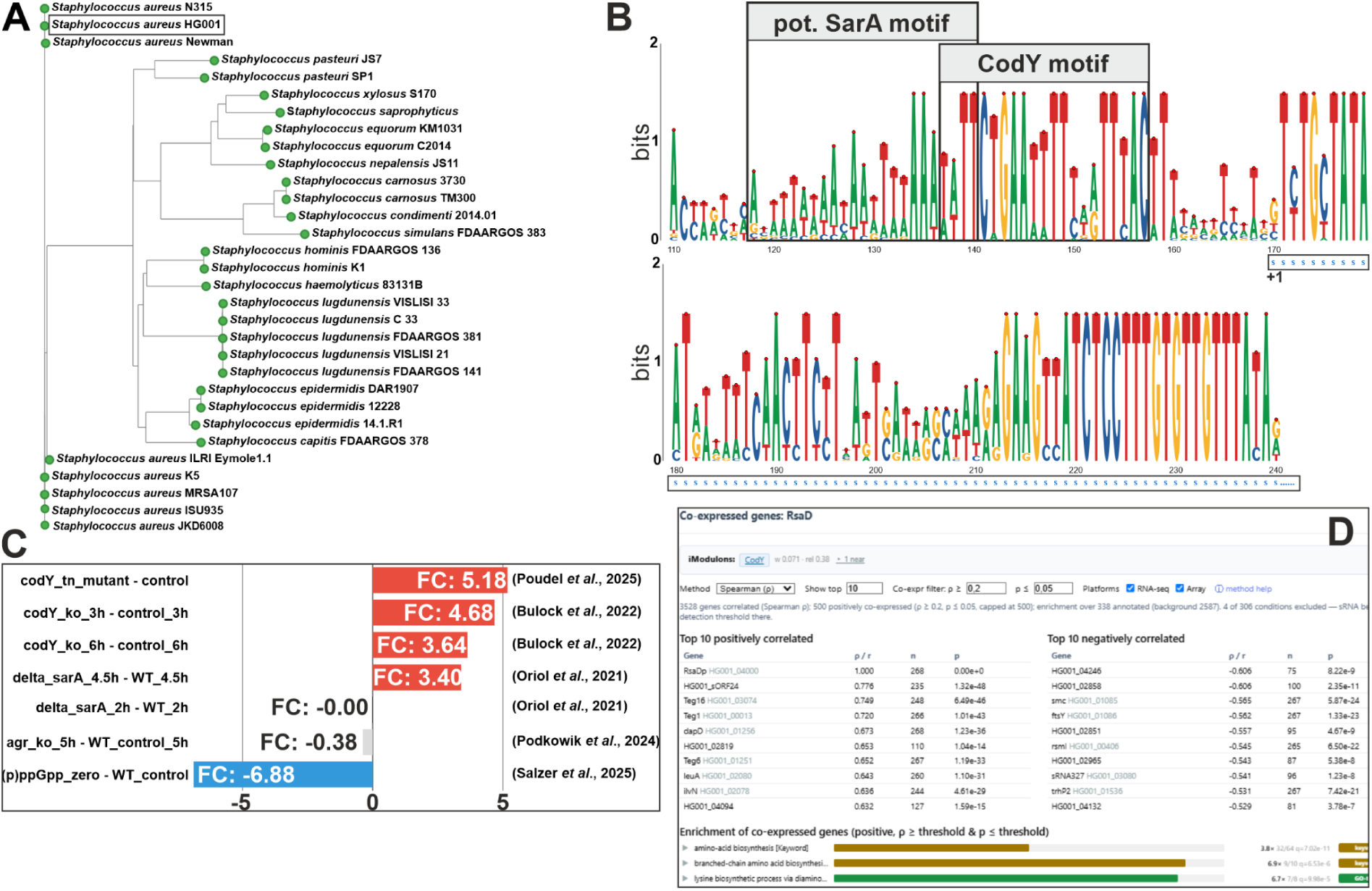
Analysis of RsaD (modified aureoRNA screenshots or exports). **(A)** Unrooted ML-Tree of 31/266 representative RsaD homologs covering the full phylogenetic distribution. The representatives are selected using a greedy filtering algorithm based on 16S rDNA and sRNA sequence distances. The procedure ensures to get the full phylogenetic distribution even if the sRNAs are very similar, while not missing multiple homologs within a single organism. **(B)** Partial sequence logo based on alignments of 31 representative RsaD homologs and their promoters. The binding site for CodY (Majerczyk et al, 2010) is indicated, as well as a predicted binding site for SarA. The start of the RsaD transcript is indicated by “+1”. The 5’ part of RsaD is shown for context and marked by a ribbon of blue “s” below the logo. **(C)** RsaD expression changes from selected conditions. An extended barplot is given in **Supplementary Figure S12**. **(D)** Correlation analysis for RsaD showing the top 10 correlated and anti-correlated (spearman) genes regarding RsaD and the top3 enriched terms based on correlated genes (ρ≥0.2, p≤0.05).

RsaD is a stress-responsive sRNA controlled by multiple signals and transcriptional regulators (Augagneur *et al*, 2020). CodY, which senses the metabolic state via cellular GTP and isoleucine concentrations (Brinsmade, 2017) is the main transcriptional repressor of RsaD (**Figure 7C**) (Gao et al, 2023; Bulock et al, 2022; Augagneur et al, 2020; Poudel et al, 2025). Our unbiased iModulon analysis based on the transcriptome data also puts RsaD correctly in a CodY-regulated module led by amino acid-and peptide transport genes (**Supplementary Figure S11),** together with three other sRNAs: Teg6 (*lysC*-UTR), RsaOB (5’ of *metI*), and Teg52 (5’ of *metN2*). Regulation by CodY is further supported by the global gene expression profiles, which show strong correlation with certain amino acid biosynthesis genes (**Figure 7D**), and by the conserved CodY motif in the RsaD promoter (**Figure 7B**).

A strong repression of RsaD was observed in mutant strains that were unable to produce the alarmone (p)ppGpp (**Figure 7C**), presumably due to increased concentrations of the CodY effector GTP (Salzer *et al*, 2025). Furthermore, the expression of RsaD has been described to be under the control of several other transcriptional regulators. It is repressed by SarA (Oriol et al, 2021), which is consistent with the presence of a conserved AT-rich motif in the RsaD promoter (**Figure 7B**), and AgrA (Marroquin et al, 2019; Augagneur et al, 2020). However, the agr axis regulation seems to be condition dependent as there was no significant effect in another *agr* knockout study (Podkowik et al, 2024). Expression is enhanced by SrrAB, which responds to nitric oxide exposure or anaerobiosis (Bronesky et al, 2019).

Another layer of signal integration occurs through the interaction of RsaD with the sRNA RsaI, which is repressed by CcpA once glucose is consumed (Bronesky et al, 2019). Interestingly, all these regulators are known to affect the *agr* response, contributing to the coordination of metabolic pathways, stress response pathways and virulence gene expression (Somerville & Proctor, 2009).

### The RsaD example: sRNA targetome refinement

The aureoRNA platform facilitates the combination of heterogeneous data (CopraRNA predictions, MAPS interactome data (Lalaouna *et al*, 2017), global gene expression correlations, and functional enrichment analyses) (**Figure 8, Supplementary Figure S13**, **Supplementary Table S1**). Each individual method is valuable but has its strengths and weaknesses, which are discussed for CopraRNA and MAPS elsewhere (Georg *et al*, 2020). The expression correlation, while informative, is an indirect measure, and also sRNA-mediated translational regulation is not necessarily accompanied by degradation/stabilization of the target mRNAs (Barrientos *et al*, 2021; Georg *et al*, 2025). Functional enrichment can detect high level patterns, but strongly relies on the quality of the input data and the gene annotation. Thus candidate targets were ranked by combining the available lines of evidence as a weighted rank product (Breitling *et al*, 2004), based on the assumption that the combination of largely independent and individually informative but noisy sources is more reliable than any one of them alone (Marbach *et al*, 2012).

**Figure 8.**
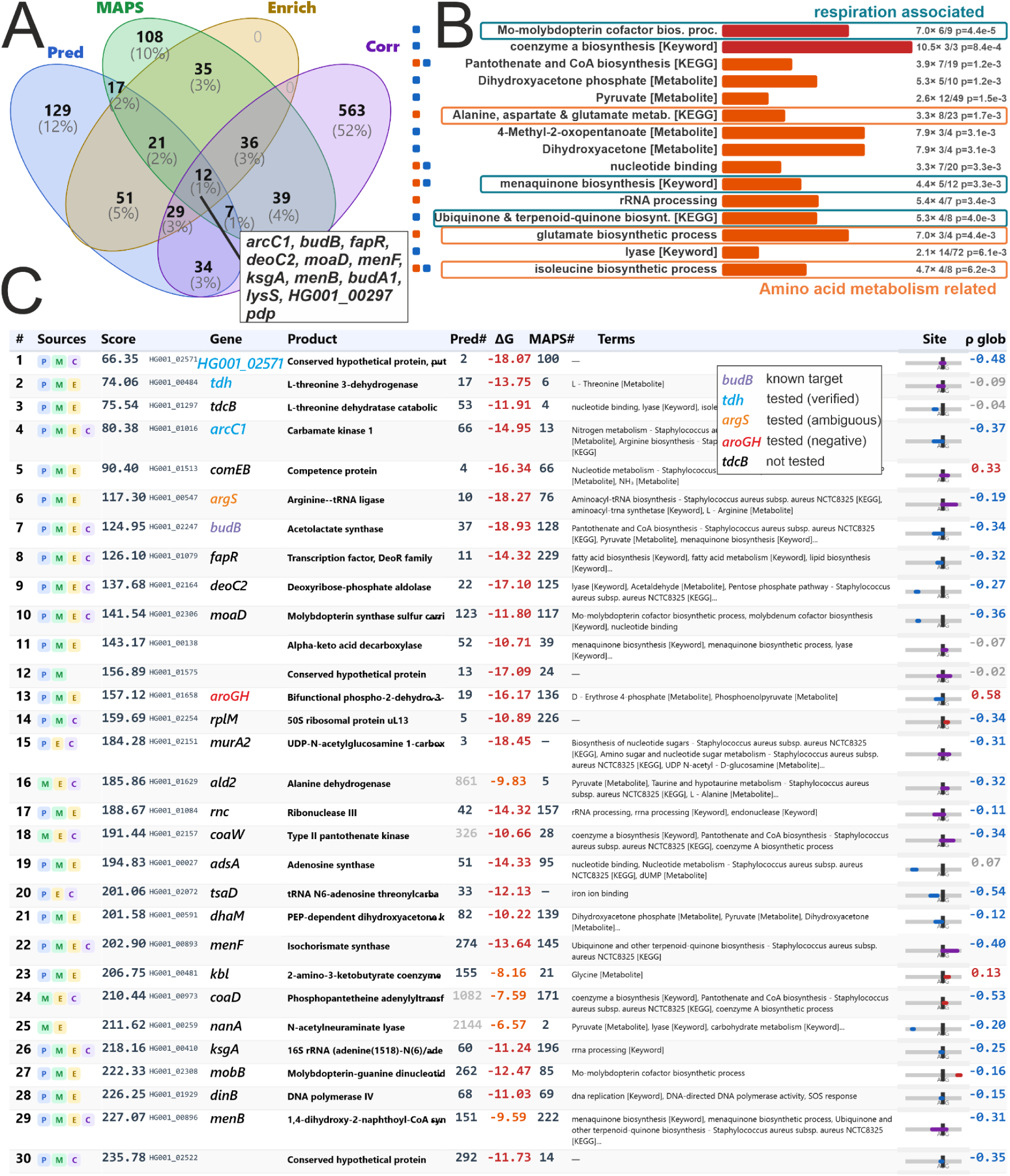
Multi-data integration for RsaD. Modified screenshots from the aueroRNA “Multi Data Integration Analysis” page (Parameters. Scoring: Rank Product | Weights: Pred=2, MAPS=2, Enrich=1, Corr=1, FC=1 | Pred Top N: 300 | MAPS Top N: 300 | Correlation (RNA-seq only): mode=global, method=standard error weighted disattenuated pearson correlation, |ρ|≥0.2, p≤0.05, only anti-correlation considered for scoring | Enrich: sources=Prediction, MAPS, N=250, enrichment foldchange ≥2, p-Value ≤0.15, enriched genes ≥3, genes in term≥3). **(A)** Venn diagram showing the overlap between the different data sources. The 12 genes that are within the thresholds for all data are shown in a box. **(B)** The top-15 enriched terms after removal of redundant terms. The left square indicates the source data for the enriched term (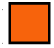 CopraRNA prediction, 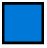 MAPS). The length of the colored bar is relative to the enrichment fold change and the color relative to the p-value. At the right enrichment fold change, enriched genes/all genes in term and the enrichment p-value are shown. All terms are shown in **Supplementary Figure S16**. **(C)** Top-30 results ranked based on the weighted rank product (smaller = better). For each candidate we provide several data: (#) combined rank based on the rank product score (smaller = better), (Sources) symbols for data sources supporting the target), (s) score (here weighted rank sum), (Gene) locus tag and gene name, (Product) annotated gene function, (Pred#) CopraRNA prediction rank, (ΔG) predicted interaction energy of the optimal interaction, (MAPS#) MAPS rank, (Terms) enriched terms for the respective gene, (Site) pictogram showing the position of the predicted interaction relative to the annotated start codon, (p glob) global correlation rho. The full refined list is available as **Supplementary Table S1**. For the full RsaD functional enrichment, see **Supplementary Figure S16**.

Throughout this analysis, the consolidated, rank-product-based rank for each proposed target is indicated (#rank). Notably, 12 candidates (*arcC1* #4, *alsS/budB* #7, *fapR* #8, *deoC2* #9, *moaD* #10, *menF* #22, *ksgA* #26, *menB* #29, *budA1* #31, *lysS* #32, *HG001_00297* [acetyl-CoA acetyltransferase] #36, and *pdp* #50) are supported by all four data sources while MAPS and CopraRNA prediction agree on 57 targets (**Figure 8A**). These 12 candidates contain the known RsaD target *budB*/*alsS* (#7, SE-weighted ρ =-0.34). The translation of the acetolactate synthase *budB/alsS* mRNA was found to be repressed by RsaD at the stationary phase of growth (Augagneur *et al*, 2020). By blocking AlsS/BudB synthesis, RsaD redirects the glycolytic flux via pyruvate away from acetoin production, a proton-consuming pathway that counteracts acidification, toward acetate production by the pyruvate:menaquinone oxidoreductase CidC (Augagneur et al, 2020). We also find *budA1* (#31, ρ =-0.35) encoding alpha-acetolactate decarboxylase as potential target, consistent with its metabolic function downstream of AlsS/BudB (**Supplementary Figure S17A**). Remarkably, RsaD transiently promotes a cell-death phenotype that is not linked to acetate production, indicating that RsaD exerts broader regulatory effects than currently appreciated (Augagneur et al., 2020).

To facilitate the planning of follow-up experiments, we used several tools that are compiled in aureoRNA. Individual targets can be investigated via the sRNA landing page (**Figure 9, Supplementary Figure S7**) and a dedicated interaction conservation page (**Supplementary Figure S14**). We used these tools to identify targets with predicted base-pairing interactions within transcribed parts of the mRNA and within ribosome binding sites. Furthermore, the predicted interaction region for selected targets is mapped to the consensus secondary structure of the sRNA (**Figure 9A**), and the genomic context of sRNA and target together with the transcriptome coverages can be visualized in two IGV viewer (Robinson *et al*, 2023) panels. A dedicated panel on the sRNA landing page shows the expression correlation between the sRNA and the selected target across the included transcriptome studies, displayed as a scatterplot and a heatmap (**Figure 9E-F**). For instance, an anticorrelation (SE-weighted ρ =-0.4) between RsaD and *HG001_02571* was observed, suggesting negative regulation at the RNA level.

**Figure 9.**
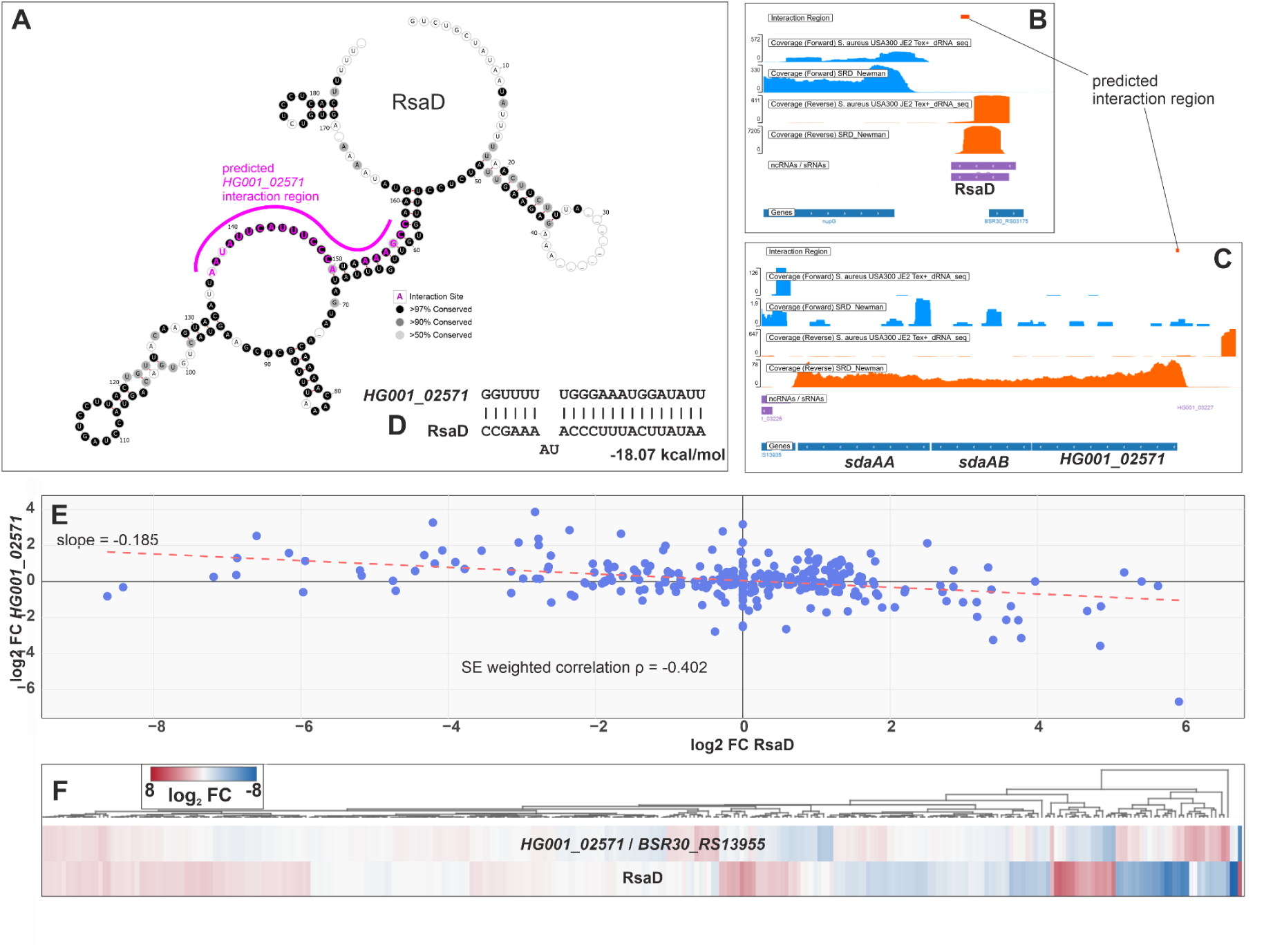
Target specific information on the sRNA landing page (Supplementary Figure S7). **(A)** Consensus secondary structure, the predicted interaction region with the selected target is highlighted in pink. Two IGV viewer (Robinson *et al*, 2023) panels show the genomic context of the sRNA. **(B)** and the selected target **(C)** together with the transcriptome coverage of the selected transcriptome datasets (dRNA-seq & SRD database transcriptome) (Das *et al*, 2016; Sassi *et al*, 2015). For reference, the predicted interaction regions are indicated by an orange bar. **(D)** Predicted IntaRNA base pairing of the selected target with the sRNA. Correlation of sRNA and selected target gene expression fold-changes across the studies included in the database in the form of a scatterplot **(E)** or a heatmap **(F)**. In the scatterplot, the linear regression is shown as a broken red line. The slope of the regression and the standard error weighted, disattenuated Pearson correlation are also shown.

However, sRNA-mediated translational regulation in *S. aureus* is not necessarily accompanied by degradation/stabilization of the target mRNAs (reviewed in Barrientos et al., 2021; Menard et al., 2022). Therefore, we selected six proposed targets for further experimental investigation (*HG001_02571* #1, *tdh* #2, *arcC1* #4, *argS* #6, *aroGH* #13, *arcD* #142) based on the combined data, prioritizing mRNAs that were enriched by MAPS and for which CopraRNA predicted base-pairing interactions within the ribosome binding sites, additionally enriched in amino acid metabolism related terms. We used two types of experiments: enzymatic probing assays to map the RsaD binding sites, and toeprinting assays to analyze the ability of RsaD to hinder formation of the ribosomal initiation complex. (**Figure 10, Supplementary Figures S18-S22**). For *tdh*, *arcC1, arcD,* and *HG001_02571*, binding of RsaD induces specific cleavages by the double-strand-specific RNase V1. These cleavages, which provide a positive signature of RsaD-mRNA duplex formation, occur within the predicted interaction sites. Furthermore, all the duplexes efficiently prevent the formation of the ribosomal initiation complexes. These data strongly suggest that RsaD acts as a translational repressor of these mRNAs. For *argS*, binding of RsaD changed the V1 cleavage pattern in a complex in a concentration dependent way but no cleavage increase was detected at the predicted interaction site (**Supplementary Figure S22)**. This may indicate an sRNA-dependent structural change. However, further experiments are required to confirm whether *argS* is a direct target of RsaD. For *aroGH*, we did not observe a significant effect of RsaD on the formation of the ribosomal initiation complex.

**Figure 10.**
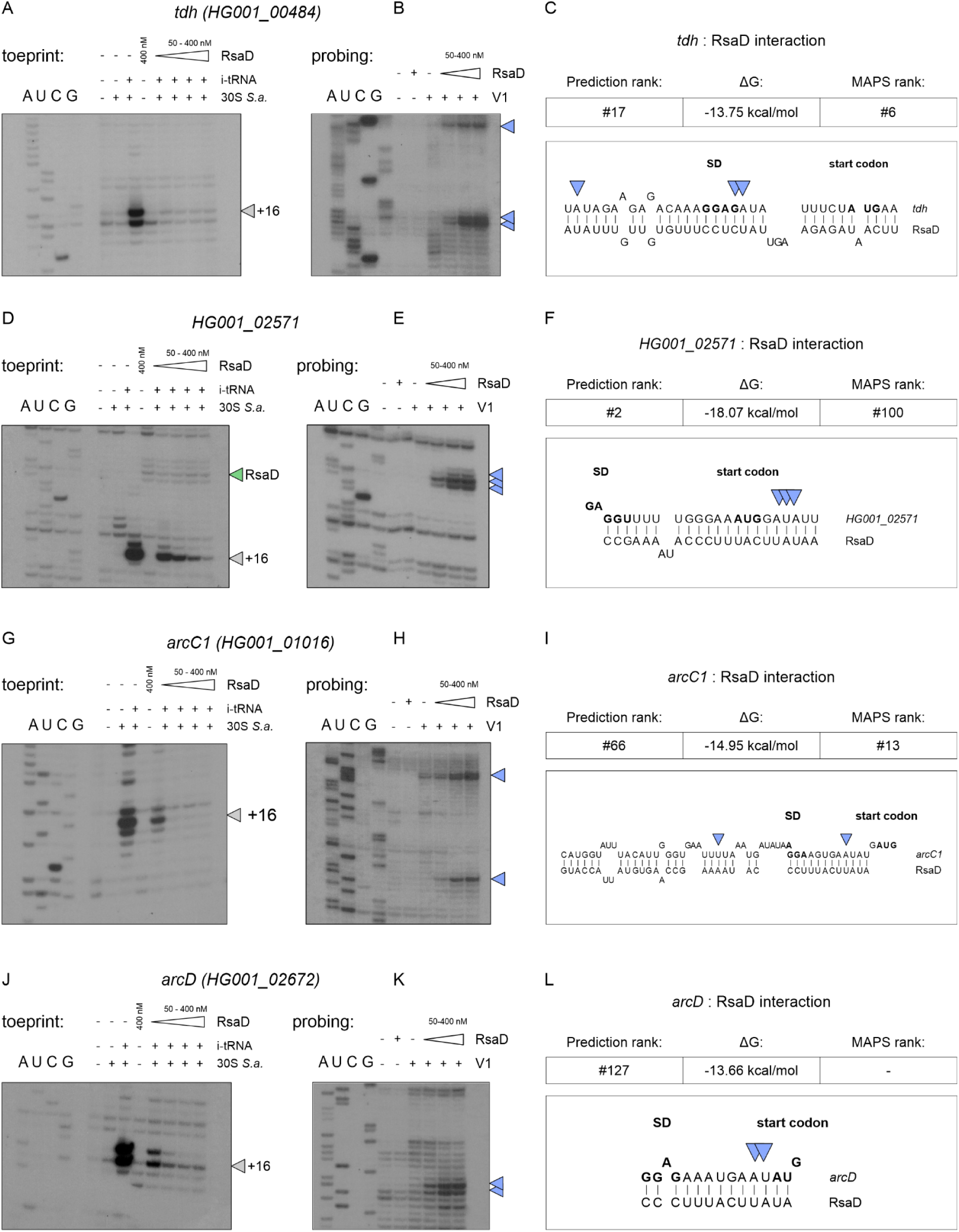
Experimental verification of predicted RsaD targets. (**A, D, G, J**) Toeprinting analysis of *tdh* (HG001_00484), HG001_02571, *arcC1* (HG001_01016) and *arcD* (HG001_02672) mRNAs, respectively, with *S. aureus* 30S ribosomal subunits and initiator tRNA (i-tRNA), in the absence or presence of RsaD. Experimental lanes, numbered from left to right after the sequencing ladders, contain mRNA alone (lane 1), mRNA with 30S subunits (lane 2), mRNA with 30S subunits and i-tRNA (lane 3), mRNA with RsaD alone (400 nM; lane 4), or mRNA with 30S subunits, i-tRNA and increasing RsaD concentrations (50-400 nM; lanes 5-8). Open arrowheads indicate toeprints at position +16 relative to the first nucleotide of the start codon. The green arrowhead in D marks an RsaD-associated reverse-transcription stop. (**B, E, H, K**) RNase V1 probing of the corresponding mRNAs in the absence or presence of increasing RsaD concentrations (50-400 nM). The first two experimental lanes contain controls without RNase V1, comprising mRNA alone or mRNA with RsaD, respectively. The remaining lanes contain RNase V1 without RsaD or with increasing RsaD concentrations. (**C, F, I, L**) Predicted base-pairing interactions between RsaD and the corresponding mRNA translation initiation regions. Shine-Dalgarno (SD) sequences and start codons are shown in bold. Computational prediction ranks, predicted interaction free energies (ΔG) and ranks from MS2 affinity purification coupled with RNA sequencing (MAPS) are indicated above each model. Blue arrowheads mark selected RNase V1 cleavage sites on the gels and pairing models. A, U, C and G denote sequencing ladders. Full experimental data are provided in **Supplementary Figs. S18-S22**.

### Integration of all data proposed that RsaD could be an altruistic bet-hedging regulator

Based on the newly confirmed and predicted targets, we propose an expansion of the *S. aureus* RsaD regulon. The two highest scoring annotated predictions are involved in threonine deamination (*tdh* #2, ρ =-0.09; *tdcB* #3, ρ =-0.04, not tested). Repressing oxidative threonine catabolism (*tdh*, *kbl* #23, ρ = 0.13) and catabolic deamination (*tdcB*) would prevent amino acid loss when branched-chain amino acid (BCAA) levels are low, redirecting metabolic flux toward isoleucine synthesis, a key CodY corepressor (**Figure 11A**). Counterintuitively, anabolic enzymes, including IlvA (#143), which initiates the anabolic threonine deamination leading to isoleucine, as well as IlvE # 156 and IlvD #170, are also predicted targets despite lacking MAPS enrichment (**Figure 11A**). Functional enrichment also suggested that several pathways related to amino acid metabolism related pathways are potentially regulated by RsaD (**Figure 8B**, **Supplementary Figure S6**). Many of those targets, such as *hom* #112 (SE-weighted *ρ*=0.70), *gltB* #62 (*ρ*=0.61), *ilvA* #143 (*ρ*=0.61), *ilvE* #156 (*ρ*=0.62)*, ilvD* #170 (*ρ*=0.70) are positively correlated with RsaD, likely reflecting the shared repression by CodY (Kaiser *et al*, 2018). Although it remains to be demonstrated, we propose that CodY and RsaD form an incoherent type 2 feed-forward loop (FFL) (Mangan & Alon, 2003). Such an architecture could accelerate regulatory responses, generate an initial expression pulse, and integrate additional signals into amino acid biosynthesis regulation via RsaD.

**Figure 11.**
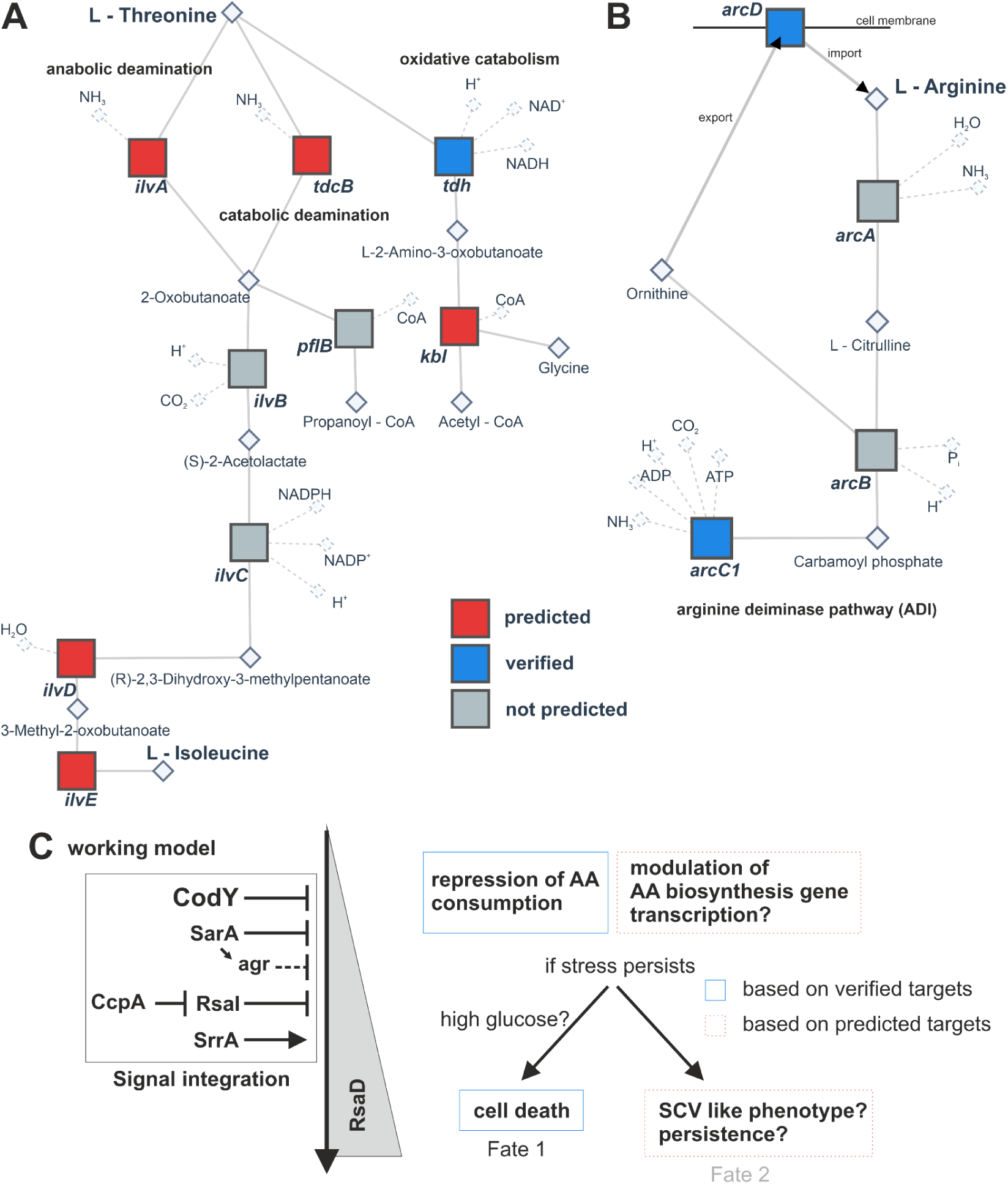
Proposed targets after the aureoRNA data integration can be mapped to their enriched terms or metabolic pathways (Modified screenshots from the aureoRNA network view page). The predicted targets within the pathways are indicated by filled red boxes, verified targets by blue boxes and other genes by grey boxes. **(A)** L-threonine dependent pathways. **(B)** Arginine deiminase (ADI) pathway. **(C)** Proposed high level working model of the RsaD response.

RsaD also targets the anaerobic, ATP-generating arginine deiminase (ADI) pathway (**Figure 11B**). Two representative validated targets are the mRNAs for the carbamate kinase *arcC1* #4 (*ρ*=-0.37) and the arginine/ornithine antiporter *arcD* #142 (*ρ*=-0.26). The *arc* operon is upregulated during mice infection and contributes to acid tolerance (Thänert et al, 2017; Lindgren et al, 2014). While RsaD represses the translation of *budB*/*alsS* which protects *S. aureus* against acid stress by channeling pyruvate into acetoin rather than acetate (Augagneur *et al*, 2020), NH_3_ production by amino acid-catabolizing enzymes would provide a complementary layer of protection directly counteracting the intracellular protons imposed by acetate accumulation (Zhou *et al*, 2019). Furthermore, the validated RsaD target *HG001_02571* (#1, *ρ*=-0.48), encodes a putative phosphotransferase system EIIC component (**Figure 9**), representing the first gene of the *sdaAB*/*AA* operon (**Figure 9C**). This suggests a link between *HG001_02571* and the catabolic deamination of serine to pyruvate and NH_3_.

Statistically significant enrichments without experimental validation point toward a repression of respiration related genes (**Figure 8B**). The involved targets, with genes for molybdenum cofactor (*moaD* #10 *ρ*=-0.36, *mobB* #27 *ρ*=-0.16, *mobA* #53 *ρ*=-0.36, *moaA* #57 *ρ*=-0.35, *moeA* #63 *ρ*=-0.39, *moaE* #72 *ρ*=-0.21), menaquinone (*menF* #22 *ρ*=-0.40, *menB* #29 *ρ*=-0.31, *menH* #71 *ρ*=-0.42, *menD* #83 *ρ*=-0.41), and heme biosynthesis (*ctaB* #33, *ρ*=-0.17) show a clear negative expression correlation to RsaD **Supplementary Figure S17C**. Because both the MoCo and menadione/menaquinone biosynthesis genes are organized in operons containing few high-ranking predicted target sites, individual strong interactions might have co-purified the polycistronic transcripts during MAPS.

Taken together, the data suggest that RsaD acts as a metabolic branch point with a layered response (**Figure 11C**). When GTP and essential branched-chain amino acids deplete and CodY repression is gradually reduced (Brinsmade, 2017; Waters et al, 2016), RsaD initially reduces the catabolic consumption of threonine, arginine, and potentially serine, while shaping anabolic fluxes via an incoherent FFL. Auxiliary inputs can be additionally integrated via SrrAB, CcpA-RsaI, and the SarA-Agr axis. Under prolonged stress or high glucose, RsaD-mediated acetate overflow (Augagneur *et al*, 2020) and impaired NH_3_ production may favor the altruistic cell death observed for *S*. *aureus* (Alqarzaee et al, 2021). Alternatively, repressing respiration genes like *menD*, for which mutations are shown to result in the slow-growing small-colony variant (SCV) phenotype (Proctor et al, 2006), may allow RsaD to promote SCV-like persistence under nutrient limitation, a hypothesis that awaits experimental validation.

## Conclusion

The aureoRNA platform facilitates the combination of sRNA target predictions, experimental interactome data, global gene expression profiles, and functional enrichment analyses for the unified characterization of the entire set of regulatory sRNAs. Starting from the annotated sRNAs and multiple available transcriptome datasets for *S. aureus* HG001 and closely related strains, we searched for sRNA homologs in the phylum Bacillota and presented their sequence and secondary structure features. In parallel, we addressed the synteny and gene expression correlations across a plethora of publicly available *S. aureus* transcriptome datasets. These expression profiles are helpful to investigate all annotated genes, far beyond the sRNA transcriptome. Our analysis allowed the genome-wide prediction of sRNA targets, which was supplemented with available experimental data for the sRNAs IsrR, RNAIII, RsaA, RsaC, RsaI/RsaOG, RsaG, and ZinS/RsaX20. Extending the approach to the sRNA RsaD, we experimentally validated four out of six newly predicted targets through toeprinting and enzymatic probing assays. The results suggest that RsaD regulates a metabolic branch point with a layered response.

General limitations of our analysis include the possibly wrong annotation of some unrecognized mRNAs encoding short proteins as candidate sRNAs. We are also aware that computational target predictions contain some false positives and false negatives, especially for longer sRNAs. Therefore, these predictions should be taken as lead, but would always require experimental scrutiny. The results are available as an interactive online resource (https://aurorna.biologie.uni-freiburg.de/) to facilitate the characterization of sRNAs in *S. aureus*. The aureoRNA website enables a holistic initial analysis of sRNAs of interest by integrating predictions with available interactome and gene expression data. Users can also leverage these datasets through various search options to identify new functions of *S. aureus* sRNAs. The goal of aureoRNA is to make all relevant sRNA data easily accessible without bioinformatics hurdles, thereby facilitating the generation of hypotheses and in-depth downstream experimental analyses.

## Materials and Methods

### Experimental Methods

#### Preparation of RNAs for *in vitro* experiments

RNAs (RNAIII, RsaE, RsaD, *citZ*, Φ11 sORF, Φ12 sORF HG001_02571, *tdh*, *arC1*, *arcD, aroGH*, HG001_00585, *argS*) were synthesized by T7 *in vitro* transcription essentially as described previously (Kohl *et al*, 2026) from PCR products containing a T7 promoter sequence directly introduced with the primer (**Supplementary Table S2**). Template DNA was subsequently digested by DNase I (Sigma), and RNA purified by preparative 6% PAGE 8 M urea gel and eluted with 0.5 M ammonium acetate pH 6.1, 1 mM EDTA, and 0.1% SDS and precipitated in cold absolute ethanol.

#### Toeprinting analysis

Toeprinting assays were performed essentially as described previously (Hartz *et al*, 1988; Fechter *et al*, 2009; Kohl *et al*, 2026) to assess inhibition of *S. aureus* 30S translation initiation complex formation by RsaD, RsaE and RNAIII on their respective predicted mRNA targets. Ternary ribosomal complexes contained *in vitro* transcribed mRNA, purified *S. aureus* 30S subunits and deacylated *E. coli* initiator tRNAᶠᴹᵉᵗ. The regulatory RNA was added together with the 30S ribosomal subunits, before the addition of the initiator tRNA. Primer extension using reverse transcriptase was used to reveal the position of the toe-print.

#### Enzymatic probing of sRNA:mRNA interactions

sRNA:mRNA interactions were analyzed by RNase V1 probing. RNA substrates were denatured and refolded individually using 1× enzymatic hydrolysis buffer [50 mM Tris-HCl (pH 7.5), 10 mM Mg(OAc)₂, 150 mM KOAc and 5 mM β-mercaptoethanol]. Refolded mRNA and sRNA were combined in the same buffer at final concentrations of 40 nM mRNA and 50 to 400 nM sRNA in a total reaction volume of 10 µL. Complexes were allowed to form for 5 min at room temperature before addition of RNase V1 at a final concentration of 0.01 U/µL in the presence of 0.1 mg/mL total tRNA. Enzymatic hydrolysis was performed for 5 min at 37 °C. Following hydrolysis, 40 µL of 0.3 M NaOAc was added, and RNA was recovered by acidic phenol/chloroform/isoamyl alcohol extraction (25:24:1) and ethanol precipitation. Pellets were washed with 80% ethanol, air-dried and resuspended in nuclease-free water. The recovered RNA was analyzed by primer extension using AMV reverse transcriptase (Promega) and ³²P-labelled single-stranded DNA oligos. Following primer extension, RNA templates were hydrolyzed, and cDNA products recovered as described for the toeprinting assays. Similarly, primer-extension products were separated by denaturing PAGE and visualized by autoradiography.

#### MS2-affinity purification

The MS2 sequence was fused to the 5’ end of the RsaD gene in the pCN51::P3-MS2-RsaD (see **Supplementary Table S2** for primer sequences). Crude extracts from MS2-RsaD and WT (control) strains were collected at an OD_600nm_ = 3. MS2-affinity purifications were conducted in duplicate, following the procedure described by Mercier et al. (2021). Bacterial cells were resuspended in Lysis Buffer (20 mM Tris-HCl pH 8, 150 mM KCl, 1 mM MgCl_2_, 1 mM DTT) and broken using the FastPrep apparatus (MP Biomedicals). Crude extracts were loaded on a MS2-MBP coated column and purified by affinity chromatography. Eluted RNAs were extracted with phenol:chloroform:isoamyl alcohol and ethanol precipitated. DNase I (0.1 U/mL, Roche) treatment was performed 30 min at 37°C. RNA quantity and quality were assessed using the 2100 Bioanalyzer (Agilent). Ribosomal RNA was depleted using the QIAseq FastSelect rRNA removal kit (QIAGEN), and cDNA libraries were prepared with the Small RNA Library Prep kit (NEB). Sequencing of cDNA libraries was performed on the NextSeq 2000 system (Illumina). RNA-seq analysis was carried out as outlined previously (Mercier *et al*, 2021).

### Bioinformatic Methods

#### sRNA homologs detection

*S. aureus* HG001 sRNA sequences were compiled from previous studies (Geiger et al, 2012; Romilly et al, 2014; Mäder et al, 2016; Caldelari et al, 2017). We analyzed 252 *S. aureus* sRNA candidates for their predicted targets and other details. The *S. aureus* HG001 sRNA sequences were used as queries against the NCBI nt database (downloaded on Oct 04, 2020) using GLASSgo v1.5.1 (Lott *et al*, 2018) and the allowed accession list “Firmicutes.acc” (10.5281/zenodo.1320179) with a minimum percentage identity of 52% to the input and londen mode turned off.

#### sRNA target prediction

Potential sRNA targets were predicted with CopraRNA (Wright et al, 2014, 2013) on a refined subset of all sRNA homolog sets using *S. aureus* HG001 as organism of interest (ooi) and its homolog sequences as reference. Sequence selection relied on the percent identity (PID) to the reference. First, sequences with a sequence identity of < 57% or 100% relative to the reference are discarded. In general, homologs were PID sorted and sampled along an ascending PID gradient using a target step interval (ΔPID) initialized at 2.9% ≤ ΔPID ≥ 3.0%. If the sequence gap exceeded 3.0%, the algorithm fell back to the nearest preceding candidate. If fewer than four homologs satisfied the step criteria across the gradient, the threshold base was adaptively decremented in steps of 0.1% until at least four homologs were captured.

### RNA-seq transcriptome data

#### Data retrieval and pre-processing

Selected public RNA-seq data (**Supplementary Table S3**) were downloaded with prefetch (SRA Toolkit) and converted to a gzip-compressed FASTQ with fastq-dump --split-files --gzip. Reads were quality trimmed with fastp (Chen *et al*, 2018) (--cut_tail-M 25), performing adapter removal, sliding-window 3′ quality trimming at a mean quality threshold of 25. Interleaved single-file paired libraries were split during this step (--interleaved_in). Reads were aligned to the *S. aureus* HG001 reference genome (NZ_CP018205.1) with bwa mem-M (Li & Durbin, 2009), and sorted by samtools (Danecek *et al*, 2021) (samtools view-b, samtools sort). Sorted BAM files were indexed with samtools index. Reads were assigned to annotated features with featureCounts (Liao *et al*, 2014) (-O-M, --maxMOp=50) against a custom *S. aureus* HG001 annotation. Paired-end and single-end libraries were counted separately, with-p enabled for paired-end data. Strandedness was determined empirically, every library was counted three times, using unstranded (-s 0), stranded (-s 1) and reverse-stranded (-s 2) settings. If the two stranded settings differed by less than 10 % of the larger value, the library was classified as unstranded and the-s 0 counts were used. Otherwise, the library was classified as strand-specific and the counts from the better-performing orientation (-s 1 or-s 2) were retained. For a small number of datasets, the strandedness was overridden manually after visual inspection of the transcriptome coverage mapping.

#### Fold-change calculation

For the fold-change (FC) calculations the normalization was done independently for all samples belonging to a given fold-change group with DESeq2 (Love et al, 2014) on stably detected features with ≥ 10 counts in at least half of the included samples. Differential expression was calculated with DESeq2 (Love et al, 2014) using a predefined treatment versus control contrast and Benjamini– Hochberg adjustment (Benjamini & Hochberg, 1995), yielding log_2_ fold change, standard error of the log_2_ fold change, and an adjusted p-value for each gene.

#### BigWig coverage tracks

First strand specific coverage was calculated. For single-end libraries, reads were split by the reverse-complement flag (samtools view-F 16 /-f 16). For paired-end libraries, read pairs were assigned to the strand of the originating transcript by combining the appropriate flag sets (first-in-pair forward together with second-in-pair reverse, and vice versa). Depth was extracted with “samtools depth-a” keeping, zero-coverage positions. For each BioProject with at least two samples in the count matrix, counts were pre-filtered to features with at least 5 counts in at least 30 % of that project’s samples (minimum two samples) and size factors were estimated with DESeq2 (Love et al, 2014). BioProjects with less than two usable samples, and any project for which estimation failed, were assigned a size factor of 1.0. Each per-base depth file was scaled by the reciprocal of the sample specific BioProject-level size factor. Replicate tracks belonging to the same condition were then averaged positionwise and formatted as bedGraph, coordinate sorted and converted to BigWig with bedGraphToBigWig (Kent *et al*, 2010). For libraries classified as reverse-stranded, the forward and reverse depth files were swapped. Because size factors are project-based, tracks are only quantitatively comparable within a given study.

#### Microarray transcriptome data

NimbleGen tiling array data for *S. aureus* (GEO accession GSE70040, platform GPL20586) (Mäder *et al*, 2016) were downloaded from the Gene Expression Omnibus using the R package GEOquery. To transfer the array design from *S. aureus* NCTC8325 to the *S. aureus* HG001 reference genome, probe sequences were extracted from the platform design file (NDF) and aligned to the HG001 genome with BWA-MEM (Li, 2013). Only probes with a mapping quality of at least 30 were retained, which keeps probes with unambiguous genomic positions. Perfect-match (PM) intensities were extracted from the raw.pair files for all remapped probes. Intensities were log_2_-transformed and quantile-normalized across all arrays using limma. Probes were assigned to genes by overlapping their genomic coordinates with the custom HG001 annotation using GenomicRanges. Probes that overlapped no annotated gene were discarded. Expression values of all probes mapping to the same locus tag were then averaged to give gene-level expression estimates. Differential expression was calculated with limma. A linear model without an intercept was fitted to the gene-level data, with one coefficient per experimental group. Conditions grown in TSB were compared with exponential-phase growth in TSB. Conditions grown in RPMI were compared with exponential-phase growth in RPMI. The two exponential-phase references were also compared directly with each other, and all remaining conditions were compared with both references. P-values were adjusted for multiple testing with the Benjamini–Hochberg method (Benjamini & Hochberg, 1995). For each contrast, the log_2_ fold change, its moderated standard error, and the adjusted p-value were stored.

### Data clustering

Fold changes from RNA-seq and microarray contrasts were combined into a single matrix and standardized row-wise (gene-wise z-scores). Cells in which a gene was not measured were treated as missing Genes with zero variance, or measured in fewer than half of the contrasts, were excluded from the distance computation and appended after the clustered rows. Hierarchical clustering was performed on the standardized fold-change matrix using Euclidean distance and Ward’s minimum-variance criterion, as implemented by the ward.D2 method of hclust (Murtagh & Legendre, 2014), separately for genes (rows) and conditions (columns). Zero-variance genes were excluded from the distance computation and appended after the clustered rows.

### iModulon analysis

Feature based RNA-seq counts as used for the differential expression were used for the analysis. Duplicated loci in the custom HG001 annotation were resolved prior to the independent component analysis (ICA). Unnamed asRNAs were excluded, while named antitoxin asRNAs and annotated small ORFs were kept. Libraries with fewer than 4 × 10⁵ assigned reads, replicate samples correlating at *r* < 0.95 with their replicates, conditions with a single surviving sample and studies without a control were removed. Counts were log-TPM transformed and centered within each study (BioProject). The final input was 3,175 features (231 annotated sRNAs) × 829 samples. Robust ICA was performed with PyModulon and scikit-learn FastICA (Sastry *et al*, 2019; Poudel *et al*, 2020). FastICA was run repeatedly from random initializations and the pooled components clustered with DBSCAN, retaining clusters reproduced across runs. Dimensionality was selected by the OptICA criterion, sweeping 250, 260, 270 and 280 components with 25 runs each and choosing the dimensionality at which the number of components conserved against lower dimensionalities equals the number of non-single-gene components. The conservation threshold (|*r*| ≥ 0.87) was derived from a two-means split of the observed best-match distribution. The criterion was met exactly at 250 components (110 conserved vs 110 non-single-gene), which was used for a final decomposition of 100 runs, yielding 117 robust iModulons. Membership thresholds were set per component by the D’Agostino K² test as implemented in PyModulon. Cut-offs from 50 to 300 were scanned and assessed on membership density, module size, sRNA coverage, and the integrity of 16 named operons (*pyr, pur, ilv, leu, spl, arc, ure, cap, nar, lac, trp, his, fad, rib, sbn, isd*). A cut-off of 150 was adopted, giving 4,577 gene-module memberships and a median module size of 26 genes. Membership is reported together with a relative weight, defined as |*M*| divided by the largest absolute weight in that component, since raw weights are not comparable between components. An sRNA is additionally listed as a near-member of a module when its absolute weight reaches at least 80% of the respective membership threshold. Components were excluded if reproduced in fewer than 55% of ICA runs, or if they were simultaneously large and low-variance (more than 88 members, the 90th percentile of module sizes in this run, and under 0.4% explained variance). 109/117 modules were retained, accounting for 66.6% of the expression variance and placing 176 sRNAs in 98 modules. Modules were named after a regulator where their highest-weighted members include experimentally documented targets of that regulator (**Supplementary Table S4**). Modules without such support were given descriptive names based on their members, their genomic locus, or the mobile element they belong to. Naming was furthermore guided by the existing *S. aureus* iModulon analyses (Poudel *et al*, 2022, 2020). 76 modules contain biologically interpretable members and 36 correspond to regulons documented in the primary literature. The iModulon decomposition was related to the differential-expression contrasts for visualization. Each heatmap column was matched to the samples of the ICA input through their run accessions, and a change in module activity was computed as the difference between the mean activity of the treatment samples and the mean activity of the control samples of that group (Δ activity). Significance was assessed per module by a Welch two-sample t-test on the activities of the two arms and corrected across modules within each contrast by the Benjamini–Hochberg procedure. Contrasts that contributed no samples to the decomposition, e.g. microarray contrasts and filtered samples, were instead projected onto the modules. Their log₂ fold-change vector *x* was decomposed in the component basis by least squares, *M a ≈ x*, over the genes shared between the contrast and the decomposition, yielding inferred activity differences *a*. Because a fold change is itself a treatment-minus-control difference, these values are activity differences of the same kind as those computed from samples, though their magnitudes are not strictly comparable. A projected activity is retained only where at least half of the module’s members were measured in the contrast. Projected activities are flagged and not reported as significant activations.

### Genome-wide metabolic model

The metabolic context was derived from a genome-scale metabolic model of *S. aureus* NCTC 8325 (Bosi et al, 2016). We used the corrected and re-annotated release of that collection provided by Renz and Dräger (BioModels MODEL2007110001) (Renz & Dräger, 2021), specifically SA_NCTC_8325_uid57795_93061_5 (1,501 reactions, 1,413 metabolites, 824 gene products). Reactions without a mapped gene were discarded, leaving 669 HG001 genes carrying at least one reaction. The metabolic context is used for functional enrichment and network drawing.

### Correlation analysis

Each gene *i* is represented by a profile of log_2_ fold changes *x_i_* across *K* differential-expression contrasts, with the corresponding standard errors *s_i_*. Fold changes and standard errors are derived from DESeq2 (RNA-seq) or limma (microarray). Contrasts in which either RNA is flagged as undetected are excluded. Also contrasts flagged as interactome datasets are excluded from all correlation analyses. Furthermore, a contrast only enters a pairwise comparison if both fold changes are finite and both standard errors are valid. The number of usable contrasts *n* therefore varies between pairs and is reported with every coefficient. Standard errors that were present but non-positive were replaced by the 75th percentile of all positive standard errors. Pearson-, Spearman-and precision-weighted Pearson correlation (Spearman, 1904; Pearson, 1895) can be calculated for the remaining pairs. Pearson’s rho is computed on the *n* available contrasts as r = Σ_k_ (x_k_ − x̄)(y_k_ − ȳ) / √(Σ_k_ (x_k_ − x̄)² · Σ_k_ (y_k_ − ȳ)²), with x̄ and ȳ the unweighted means over those conditions. Spearman’s ρ is obtained by replacing each profile with the ranks of the retained contrasts, tied values receiving the average of their ranks. Because the precision of fold changes can vary significantly by orders of magnitude in the dataset, each contrast is weighted by the combined precision of the two measurements, so that unreliable contrasts do not contribute to the coefficient as strongly as well determined ones. For that reason, the default estimator is the precision-weighted Pearson correlation with correction for attenuation (Spearman, 1904). Weights are w_k_ = (s²_x,k_ + s²_y,k_ + δ)^−α^ with δ = 10⁻⁶ and α = 0.5 by default (α = 1 is strict inverse-variance weighting and α = 0 is unweighted), with standard errors floored at 0.05 and W = Σ_k_ w_k_. Weighted means, variances and covariance are calculated as x̄ = Σ_k_ w_k_ x_k_ / W, ȳ = Σ_k_ w_k_ y_k_ / W, v_x_ = Σ_k_ w_k_ (x_k_ − x̄)² / W, v_y_ = Σ_k_ w_k_ (y_k_ − ȳ)² / W and c = Σ_k_ w_k_ (x_k_ − x̄)(y_k_ − ȳ) / W, leading to the weighted coefficient r = c / √(v_x_ v_y_). Correction for attenuation: Because each fold change is an estimate with error, the observed variance of a profile is inflated while the covariance between two profiles is not, so an observed correlation is systematically shifted towards zero. The correction removes this bias so that pairs measured with different precision remain comparable. The mean error variance of each profile is calculated with the same weights, using the profile standard errors, ē_x_ = Σ_k_ w_k_ s²_x,k_ / W, ē_y_ = Σ_k_ w_k_ s²_y,k_ / W, and the observed variances are deflated to true-score variances v^T^_x_ = v_x_ − ē_x_ and v^T^_y_ = v_y_ − ē_y_, with reliabilities λ_x_ = v^T^_x_ / v_x_ and λ_y_ = v^T^_y_ / v_y_. If λ_x_ and λ_y_ both reach 0.2 and both deflated variances exceed 10⁻⁹, the coefficient is reported as r_corrected_ = c / √(v^T^_x_ v^T^_y_) = r / √(λ_x_ λ_y_), otherwise the uncorrected weighted r is reported and the result flagged. The effective sample size is n_eff_ = (Σ_k_ w_k_)² / Σ_k_ w_k_².

Two-sided p-values are obtained from t = r √((n − 2)/(1 − r²)) on n − 2 degrees of freedom, evaluated through the regularised incomplete beta function. For the unweighted estimators, n is the number of conditions used. For the weighted estimator, n is replaced by n_eff. The p-value is computed from the weighted coefficient before the attenuation correction, since the correction rescales a coefficient without adding information. Too few conditions, insufficient effective sample size (n_eff < 25), insufficient reliability, or an absence of usable measurements are labeled with a quality flag. p-values are corrected by the Benjamini–Hochberg procedure (Benjamini & Hochberg, 1995).

### Functional enrichment analysis

#### Annotation sources

Gene Ontology terms and UniProt keywords for *S. aureus* NCTC 8325 were retrieved from UniProt (The UniProt Consortium, 2025), and KEGG pathway assignments from KEGG (Kanehisa et al, 2025), and mapped onto the *S. aureus* HG001 annotation. Metabolite terms are defined from the metabolic network (Renz & Dräger, 2021; Bosi et al, 2016) as the set of genes producing or consuming a given compound. Compounds associated with fewer than two genes are skipped. Ubiquitous cofactors and currency metabolites are also excluded from the automatic scan. Gene universes: The background universes for all enrichment cases comprise all annotated genes, i.e. every gene carrying at least one term. Genes without annotation are excluded. Candidate set - target integration: Here the enrichment is done on the target set of a given sRNA candidate. Candidate sets are assembled per enrichment job as the union of the evidence sources selected for that job (the top N ranked target predictions, the top N MAPS-ranked genes, genes passing the fold-change criteria, and genes passing the correlation criteria with an optional restriction to negatively or positively correlated genes) with N = 200 and top ranked predictions used as default. Each job is tested against every term, and several jobs may be defined and compared side by side. Candidate set-reverse search: Here all sRNAs target sets are scanned against predefined terms. For each sRNA in the database the top *X* ranked predicted targets form the candidate set (CopraRNA predictions only), (*X*: default 20, maximum 3000). One test is performed per term and sRNA pair.

#### Statistical test

All term categories are tested identically, by a single shared implementation, so that p-values and fold enrichments are comparable across categories and can be pooled into one ranking. For a term containing *K* of the *N* annotated genes, with *k* of the *n* candidate genes belonging to the term, enrichment was assessed by a one-sided Fisher exact test. That is the hypergeometric probability of observing at least *k* term genes in a set of *n* drawn at random from the *N*. Fold enrichment is reported as *k* divided by its expectation value *Kn*/*N*. Terms containing fewer than three genes of the universe, or with fewer than two candidate genes, are not tested. Multiple testing and thresholds: P-values from all categories (Gene Ontology, keyword, pathway and metabolite) are corrected together by the Benjamini– Hochberg procedure (Benjamini & Hochberg, 1995), applied once over the whole set of tests. In target integration over all terms tested for one enrichment job, and in the reverse search over every term and sRNA pair evaluated in that search.

### Combined target prediction

#### Evidence sources

For each sRNA, candidate targets are assembled from up to five lines of evidence. 1. CopraRNA computational target prediction; 2. Experimental MAPS ranking; 3. Membership to a significantly enriched functional term; 4. Fold-change profile correlation with the sRNA; 5. Selected differential expression. By default, a gene enters within the top 200 predictions, within the top 200 MAPS target, if differentially expressed at |log₂FC| ≥ 1 and p_adj ≤ 0.05 in at least one selected contrast, as correlated at |ρ| ≥ 0.2 and p ≤ 0.05 across all non-interactome contrasts, and as functionally enriched when it belongs to a term with fold enrichment ≥ 2 and p ≤ 0.1 based on at least three background genes and two candidate genes. Significance thresholds apply to uncorrected p-values unless Benjamini–Hochberg correction is enabled, and all thresholds are user-adjustable. Every data source is ranked over the whole candidate gene set. Prediction (p-value sorted) and MAPS (fold change sorted) contribute their native ranks directly. The remaining three have no intrinsic rank and are ranked by an effect-size × significance statistic (Functional enrichment: fold enrichment × −log₁₀(p), Correlation: |ρ| × −log₁₀(p), Selected fold-change(s): |log₂FC| × −log₁₀(p_adj)). Tied values receive the average of their ranks, so that genes with identical evidence are not separated by an arbitrary order. A gene with no evidence from a source is assigned the worst available rank rather than being excluded, so that a missing source only penalizes a candidate. Two different scoring methods are available. Weighted rank product (default): Ranks are combined as a weighted geometric mean, the weighted form of the rank product (Breitling et al, 2004). Source weights are user-adjustable (arbitrary default: 2 for target prediction, 2 for MAPS and 1 for enrichment, correlation and differential expression). S = (Π_s_ r_s_^ws^)^(1/Σ_s_ w_s_), with r_s_ the gene’s rank in source s and w_s_ that source’s weight; lower values indicate stronger combined support. Because S depends on how many sources cover a gene, raw scores are not directly comparable between candidates of differing coverage. Each score is therefore compared only with scores generated from the same combination of sources. There are at most 2⁵ = 32 possibilities, and one null distribution is generated per combination and shared by all genes having it. Each of B = 2000 draws resamples, independently per source the observed ranks with replacement, preserving the empirical rank distribution including the tie blocks produced by the threshold cut-offs. The p-value is the fraction of null draws scoring at least as well as the observed value, with one pseudo-count added to numerator and denominator: p = (1 + m)/(1 + B), where m counts the draws with S_null_ ≤ S_obs_, and is corrected across all candidates of a run by the Benjamini–Hochberg procedure. Confidence score: As an alternative to the rank product, a custom additive score on a fixed 0–100 scale is provided. Each source gets a maximal point budget B_s_ = 100·w_s_/Σw proportional to its weight. Rank-based sources decay linearly with rank position, contributing B·max(0, 1 − rank/(N+1)) for a search depth of N. The remaining sources split their budget equally between effect size and significance, each term saturating so that no single extreme value dominates: enrichment contributes B·[0.5·min(fold,10)/10 + 0.5·min(−log₁₀p, 5)/5], differential expression B·[0.5·min(|log₂FC|,5)/5 + 0.5·min(−log₁₀p_adj, 5)/5], and correlation B·[0.5·√(min(|ρ|,0.5)/0.5) + 0.5·min(−log₁₀p, 5)/5]. Higher values indicate stronger support. Both scores are heuristic summaries intended for ranking candidates for experimental follow-up. They are not a calibrated quantity. Since only few experimentally verified sRNA/target interactions are available and validated true negative targets are rarely reported, no usable benchmark for fitting the weights is available.

#### Sequence logos

Per-position sequence conservation is displayed as sequence logos computed in the browser, following the approach of WebLogo (Crooks *et al*, 2004) but implemented independently. For each sRNA, the multiple sequence alignment of its homologues together with the corresponding upstream promoter regions, the accompanying phylogenetic tree and an annotation track are retrieved from the file store. Sequences entering the logo are selected interactively on the tree, and the logo is recomputed from the current selection, so that conservation can be examined within a clade as well as across the whole alignment. At each alignment column, base counts n_i_ are taken over the selected sequences, gaps and ambiguous positions being excluded, so that the effective number of sequences contributing to a column varies along the alignment. Column height is the expected relative entropy of the column composition against a uniform background (0.25 per base), estimated as the posterior mean under a Dirichlet prior, R = [−Σ_i_ (α_i_/A)·ln q_i_ − E(H)] / ln 2, with E(H) = ψ(A+1) − Σ_i_ (α_i_/A)·ψ(α_i_+1), where α_i_ = n_i_ + λ, A = Σ_i_ α_i_, q_i_ is the background frequency and ψ the digamma function. The pseudocount λ is 1 when the Bayesian correction is enabled and negligible otherwise. R is constrained to be non-negative, For the four-letter alphabet it ranges from 0 to 2 bits. Within each column, letters are scaled to the product of their observed frequency and the column height and stacked in order of increasing frequency from the baseline, so that the most frequent base appears on top.

#### Conserved structure prediction

The representative sRNA homolog sequences without promoter and trailing 3’ extensions were aligned with MAFFT (Katoh & Standley, 2013) (maxiterate 1000, localpair). The alignment was the input to create a consensus secondary structure with RNAalifold from the ViennaRNA package (Bernhart *et al*, 2008; Lorenz *et al*, 2011), using partition-function folding (-p), RIBOSUM covariance scoring (-r), the-d2 dangling-end model and suppression of isolated base pairs (--noLP), with results written in Stockholm format. The resulting alignment and consensus structure were tested for evidence of conserved RNA structure with R-scape (Rivas *et al*, 2017). R-scape returns the alignment as an R2R-annotated Stockholm file (Weinberg & Breaker, 2011). The file was parsed into a compact JSON representation holding the consensus sequence, the consensus structure in dot-bracket notation and one color per position. In the browser the consensus structure is drawn with Forna (Kerpedjiev *et al*, 2015), which renders the dot-bracket structure as a secondary-structure diagram with nucleotides as nodes and backbone and base-pairing as edges. The colors assigned to each position are provided as a custom color scheme, enabling the diagram to display both structural topology and evidential support. Furthermore, when a predicted target is selected, the sRNA nucleotides predicted to pair with it are marked on the structure.

#### sRNA synteny and phylogenetic distribution

To analyze and extract the synteny of sRNAs we used Syntney (https://github.com/CyanolabFreiburg/Syntney) based on the GLASSgo output fasta file. To construct a synteny network, for each sRNA, 4 upstream and 4 downstream protein coding genes, if available, were extracted from the respective genbank files. The genes were clustered based on the amino acid level with CD-HIT (Li et al, 2001) (-n 2,-aL 0.6,-d 50,-c 0.4). All sRNA homologs were combined in a single sRNA cluster. The final synteny network reflects all clusters and connections that exist in the input syntenies, with edges pointing in the direction of the central sRNA cluster. Each cluster gains a score based on the number of occurrences in the input syntenies. To account for the non-uniform organism coverage, a sum of branch (sob) normalization of the count score was applied. The sob-weighting was based on a 16S rDNA neighbor-joining tree of all input organisms. In order to score a network cluster, the lowest common ancestor node (LCA) of all organisms contributing to the respective cluster was identified. Next, the length of all edges from the LCA to tips, representing organisms involved in the investigated cluster, and the edge from the LCA to its parent node were summed up and divided by the total tree length, i.e. the sum of all edges. Edges leading to tips of organisms not included in the investigated clusters were not considered. Finally, the sob-weighted cluster scores of the network were normalized to sum up to one. Accordingly, the edges are scored by the sob weighted occurrences of the respective connection in the input syntenies. In order to create a sRNA/synteny network we next built a symmetric adjacency table storing the weighted number of co-appearances of clusters in the input syntenies. To account for the non-uniform organism coverage for different phylogenetic groups a weighting procedure based on 16S rDNA was used. To account for multiple copies of an sRNA in an organism also the sRNA sequences were used for the weighting. Weights were calculated for the 16S rDNAs and the sRNAs as in ClustalW (Thompson et al, 1994). The final weight for each sequence is the geometric mean of both individual weights. Each co-occurrence of two protein clusters in an input synteny is then counted as by the respective sRNA weight. The resulting matrix was used to identify synteny families by Markov clustering (Enright et al, 2002) using the R MCL package (addLoops = F, expansion = 2, inflation = 2). Before drawing the phylogenetic trees based on the sRNA sequences, sequence redundancy was reduced. Using a greedy procedure all sequences with a distance below a given threshold for the sRNA sequence and the 16S rDNA sequence to a reference sequence are removed if the synteny belongs to the same synteny family. This procedure is repeated until no remaining sequence fulfills the exclusion criteria. The threshold was dynamically adapted to yield a maximum of 150 remaining representative sequences. For the representative homologs sequences 150 nt upstream the genomic position of the first nucleotide of the sRNA and 50 nt downstream were extracted and aligned with MAFFT (Katoh & Standley, 2013) (maxiterate 1000, localpair). Based on the alignment a distance matrix was generated with the R package phangorn (model=F81). Subsequently, phangorn was used to calculate a neighbor joining tree, which served as the starting point for a maximum likelihood tree. The alignments were also used for the sequence logo described below.

#### System architecture

The resource is a three-tier application: a MongoDB document store, a Node.js REST service built on Express, and a browser front end of self-contained HTML pages. The service exposes read endpoints under /api, serves the static pages, compresses responses above 1 kB, and maintains a connection pool to a single MongoDB instance. Interactive analyses, e.g. correlation of fold-change profiles, functional enrichment, candidate scoring, hierarchical re-clustering of displayed subsets and figure export are computed in the browser from data the API delivers in bulk. The statistical routines are located in shared JavaScript modules so that pages performing the same computation cannot diverge in their results.

Content is divided between document collections and a binary store according to how it is queried. Record-level data occupies indexed collections, one document per sRNA, one per target prediction, and separate collections for gene annotations, strain annotations, synteny relationships, user-contributed wiki content and user accounts. Precomputed matrices and other whole-dataset snapshots are held in a reference collection keyed by data type, covering the combined fold-change matrix and its value arrays, expression patterns, iModulon decompositions and sample metadata, transcriptome metadata and genome feature tables. Large numeric arrays, fold changes, adjusted p-values, scaled values and standard errors, each with one entry per gene and contrast, are stored as ordered chunks of 400,000 values and reassembled by the API on request. Sequence alignments, structure predictions, coverage tracks and genome files are stored in a GridFS bucket and streamed on demand.

Indexes for the access patterns of the interface are created by the loading script once the collections are populated and are maintained incrementally afterwards. Single-field indexes on locus tags, gene names, GO identifiers and keywords for annotation lookup, a text index on strain annotation descriptions for free-text search. For target predictions single-field indexes on sRNA and target identifiers together with compound indexes on sRNA and rank and a covering index spanning sRNA, rank, target locus tag, gene name and significance, so that the ranked prediction lists underlying every analysis are served without document fetches.

#### Declaration of generative AI use

The software described in this work was partly written with the assistance of a large language model (Claude, Anthropic). This also covers the code of the aureoRNA web resource. The model was used for code drafting, refactoring and debugging. It was not used to generate experimental data or for analytical decisions. All code was read, executed and tested by the authors. Candidate iModulon module names and regulator assignments were drafted with model assistance from published literature and subsequently verified by the authors. AI was also used for language editing of the manuscript and to draft method sections based on the used code.

## Data Availability

The RsaD MAPS data have been deposited in GEO under accession number GSE355316. The Firmicutes accession list can be found on zenodo (<u>10.5281/zenodo.1320179</u>).

## Ethics declaration

Not applicable

## Author contributions

JG, SM, IC, PR and WRH designed the study. FH, SH, SL and JG developed the workflow. MPK and IC performed experimental analyses. FH, JG, SM, IC, MPK, DL, PR and WRH analyzed data. EA and KV are responsible for deploying and hosting the software and database on the web server. JG, SM, IC, PR and WRH drafted the manuscript. All authors read and approved the final manuscript.

## Acknowledgments

We appreciate financial support by the joint DFG-ANR program to PR and WRH (grants HE 2544/22-1, project no. 530000184, ANR-23-CE12-0041-01 (IntRNAReg)). MPK is supported by funds from IdEx Unistra (ANR-10-IDEX-0002) and by EUR IMCBio (ANR-17-EURE-0023). We thank Delphine Bronesky for the construction of the MS2-RsaD vector.

