## Supplemental Figures S1-22 & Table S2 for "aureoRNA enables the interactive, genome-wide analysis of *Staphylococcus aureus* small RNA targetomes and uncovers new regulatory functions"

[$]Co-sharing first authors

*Co-corresponding authors

### Supplementary Material

Supplementary Table S2: page 2

Supplementary Figures: page 4

Supplementary References: page 30

The additional Supplementary Dataset contains tables S1, S3 and S4

### Supplementary Table S2. Desoxyoligonucleotide primers used in this work.

| Name | Sequence 5'-3' | Purpose |
|---|---|---|
| oMK56 | TAATACGACTCACTATAgacataaaaatgtggattctagg | Φ11 sORF t7 fwd |
| oMK57 | ggatttttgagccatagcat | Φ11 sORF rev |
| oMK58 | gactcttaactgttatttacatttac | Φ11 sORF toe/probing |
| oMK59 | TAATACGACTCACTATAgacatcaagatgtggattctagg | Φ12 sORF t7 fwd |
| oMK60 | gcttttgatttgcccatttac | Φ12 sORF rev |
| oMK61 | gactcttaactattacttacacttac | Φ12 sORF toe/probing |
| oMK576 | TAATACGACTCACTATAGgaaagccatttcataaagaaac | t7+ fwd citZ |
| oMK577 | gatgagcaagctcttcttcg | citZ rev |
| oMK578 | gaacttattttagtctccg | citZ toeprint oligo 1 |
| oMK580 | TAATACGACTCACTATAGGtgaaattaatcacataacaaac | t7 fwd rsaE |
| common oligo lab Romby 623 | AAAAAAACGTCGTGTCTGAATAC | rsaE rev |
| oMK751* | TAATACGACTCACTATAgtctgctataatgtttttaacttcttc | rsaD fwd t7 |
| common oligo lab Romby 899 | AAAAACCAAGTGCACATGGTAA | rsaD rev |
| oMK752* | TAATACGACTCACTATAgtctaataaatatataaaggggatgg | HG001_00585 t7 fwd |
| oMK753* | cctatggcaatcaatactgcg | HG001_00585 rev |
| oMK754* | catttacacctgcagcaaag | HG001_00585 toe/probing |
| oMK755* | TAATACGACTCACTATAggtctgaaaataaaataaacatagaga | HG001_02571 t7 fwd |
| oMK756* | gccaccaatagcataacttaag | HG001_02571 rev |
| oMK757* | gtgcatactttaaaaatagcgtc | HG001_02571 toe/probing |

| | | |
|---|---|---|
| oMK758* | TAATACGACTCACTATAgaagatagaaacgaaaatttttcttaaaag | aroGH t7 fwd |
| oMK759* | gtgctagttcaccacgtttag | aroGH rev |
| oMK760* | gtctaaaatttgatgattcagtg | aroGH toe/probing |
| oMK761* | TAATACGACTCACTATAgaaaatgaatattatgtccgcttg | tdh t7 fwd |
| oMK762* | caaatggtccattttgtacagg | tdh rev |
| oMK763* | ctctgcacttaacaactaattc | tdh toe/probing |
| oMK763.5* | TAATACGACTCACTATAgaaaaaatgggttcttatcaatatg | argS T7 fwd |
| oMK764* | gtaccatcgcaatattagtagc | argS rev |
| oMK765* | ctaatcctgctttgttaatacttg | argS toe/probing |
| oMK766* | TAATACGACTCACTATAggttgctacattgggtg | arcC1 T7 fwd |
| oMK767* | gtggtccattaccatgacta | arcC1 rev |
| oMK768* | caagctcgagttgttcttg | arcC1 toe/probing |
| oMK30 | TAATACGACTCACTATAgcugaaauggaaguuacagacc | arcD t7 fwd |
| oMK31 | cgaacgctaatgaaatcattcc | arcD rev |
| oMK33 | ggatcctataactagtccaatt | arcD toe/probing |
| MS2-RsaD | CGGGATCCGTACACCATCAGGGTACGTTTTTCAGACAC CATCAGGGTCTGGTTTgtctgctataatgtttttaac | MAPS |
| RsaD rev | CGGAATTCgttaccgaaattgaagcattataa | MAPS |

### Supplementary Figures

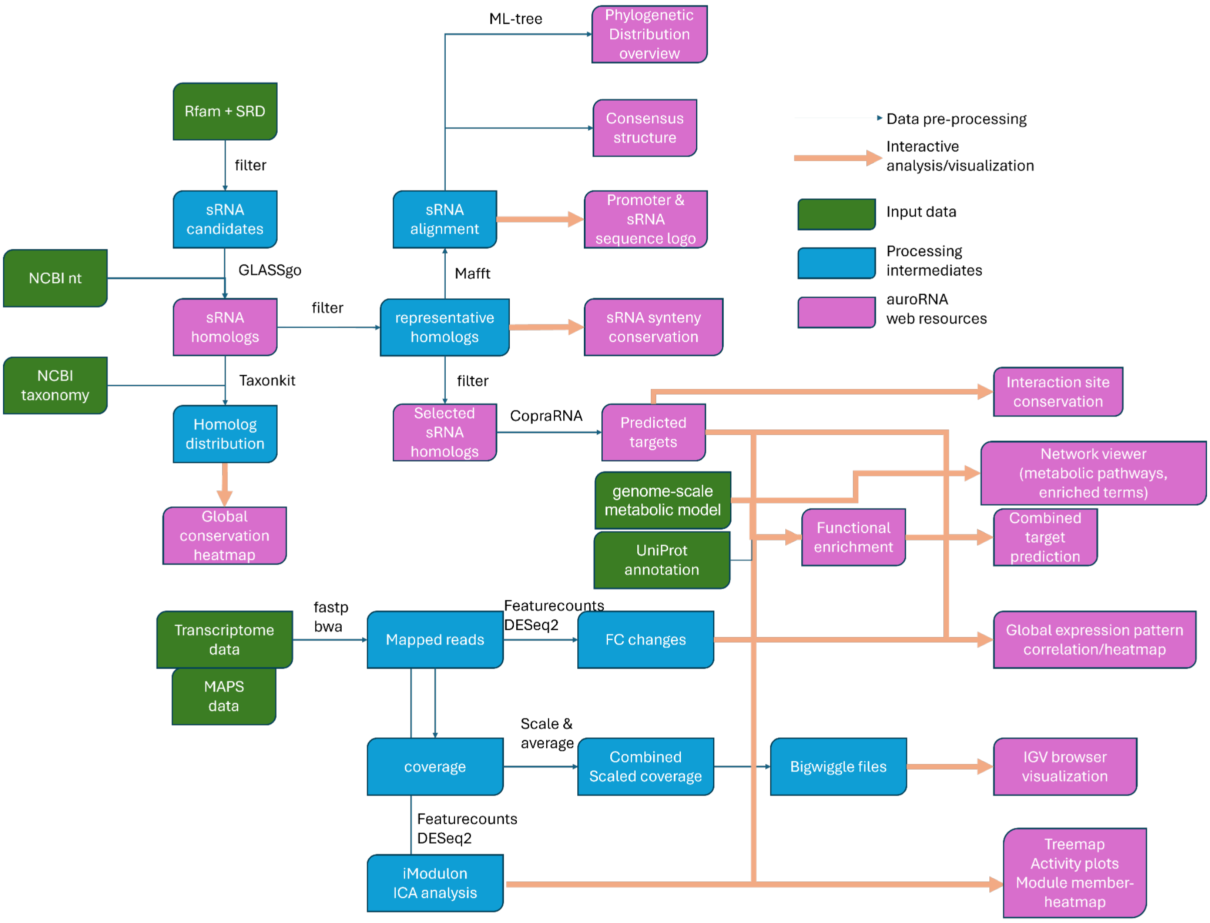


**Supplementary Figure S1.** Flow-chart of the biocomputational analyses.

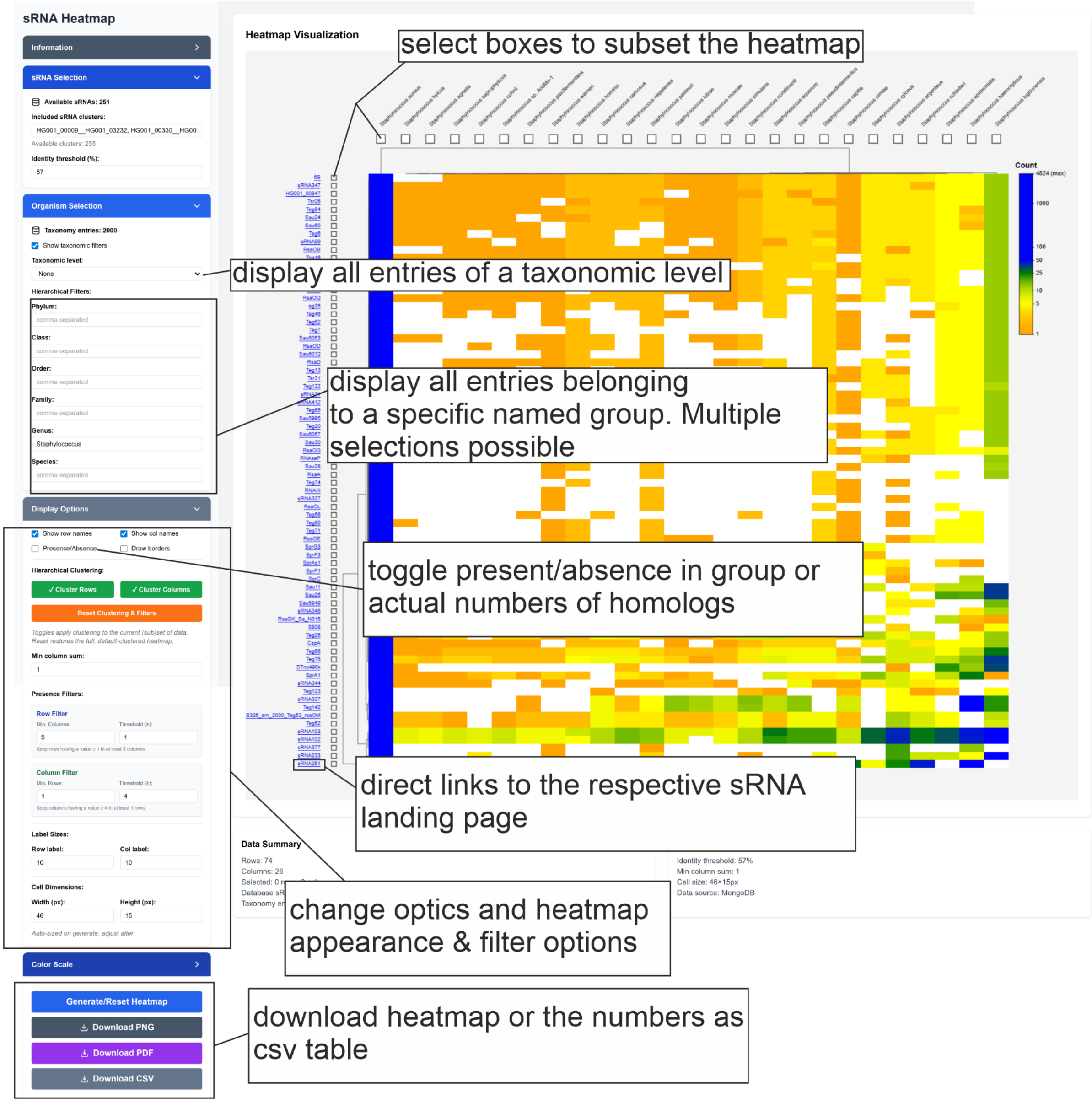


**Supplementary Figure S2**: Page for taxonomic sRNA analysis based on the GLASSgo results. This figure extends **Figure 1**. For detailed explanations, use the “guided tour” on the website.

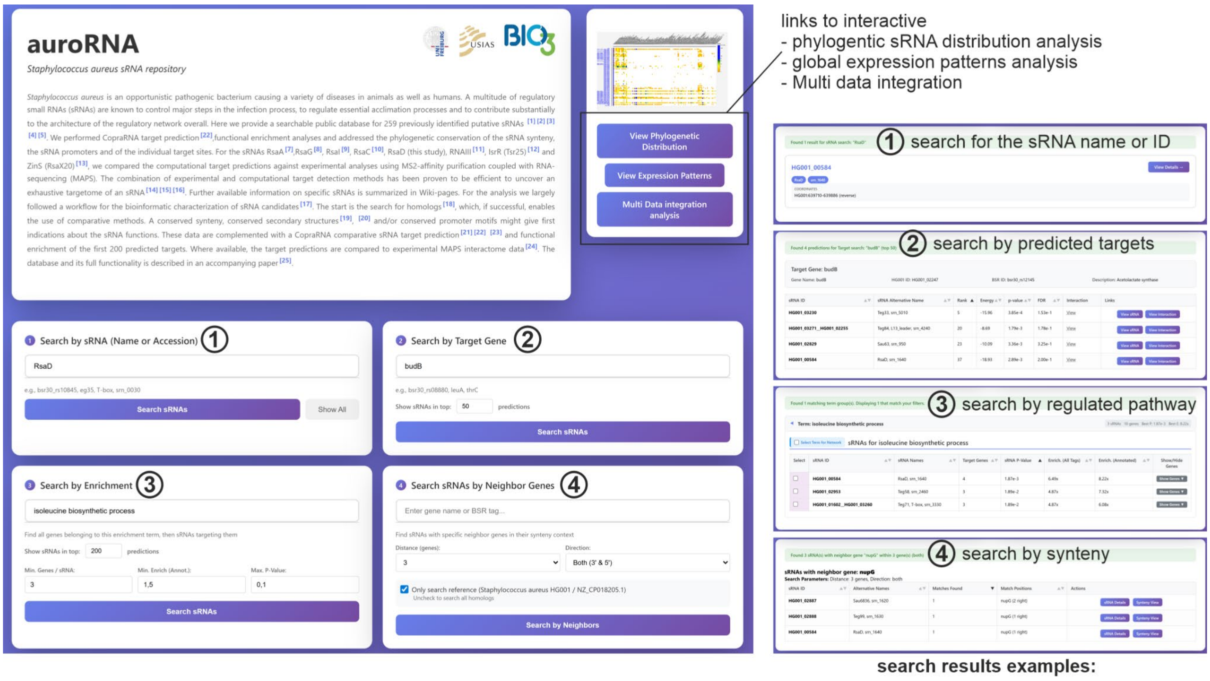


**Supplementary Figure S3.** Start page of aureoRNA with different search options for sRNAs of interest. For a full explanation, please use the “guided tour” on the website.

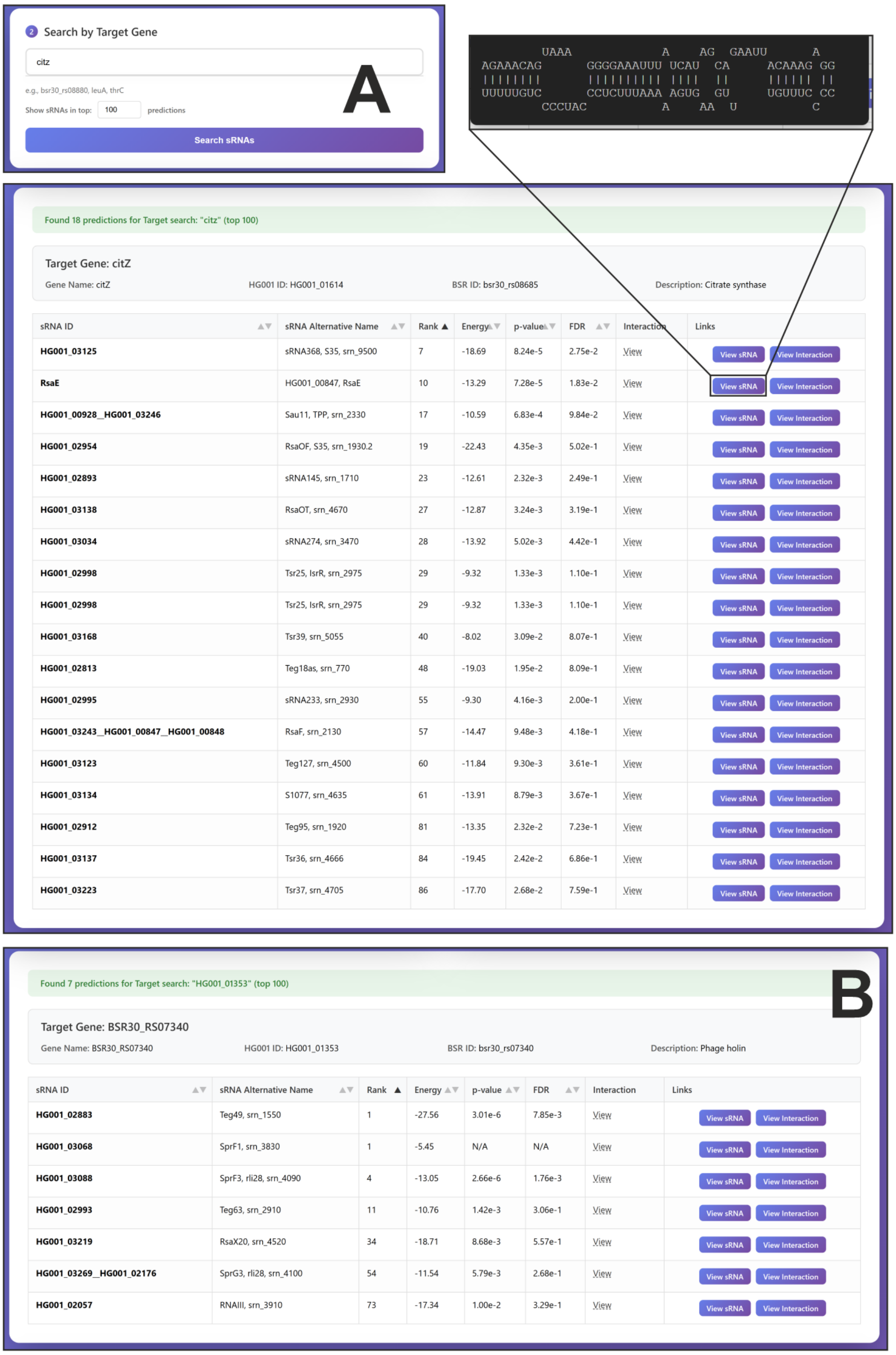


**Supplementary Figure S4.** This example illustrates a target-centric sRNA search. In this example, the search identifies sRNAs that include *citZ* **(A)** and *HG001_01353* **(B)** among their top 100 predicted targets. The *citZ*/RsaE and *HG001_01353*/RNAIII interactions were validated in this study. Within the results, the interaction region can be previewed via a tooltip, while a direct link navigates to the dedicated sRNA landing page (**Supplementary Figure S7**).

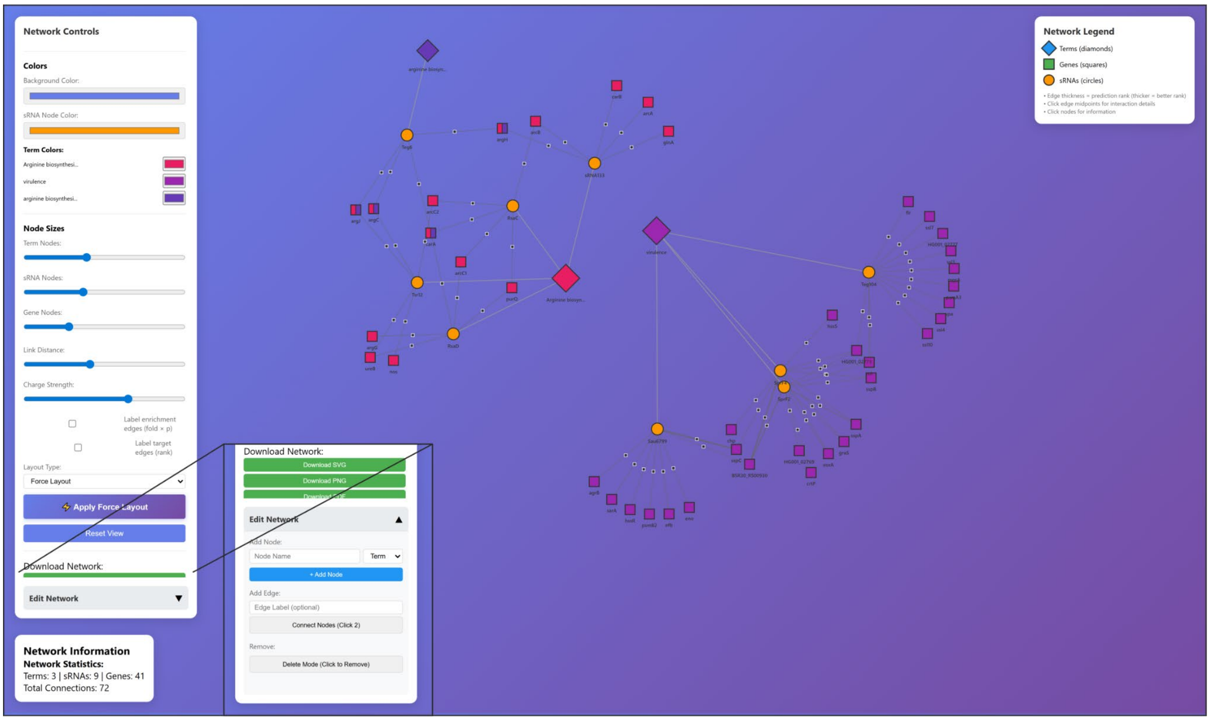


**Supplementary Figure S5.** aureoRNA network builder based on a Cytoscape plugin (Franz *et al.*, 2023). The tool can be accessed via the primary search page for selected enriched pathways or the “Multi data integration analysis” (**Supplementary Figure S13**) page for pathway networks. Networks can be modified in multiple ways (colors, symbols, adding and deleting nodes and edges, change names). The resulting network can be downloaded as png. svg or pdf file.

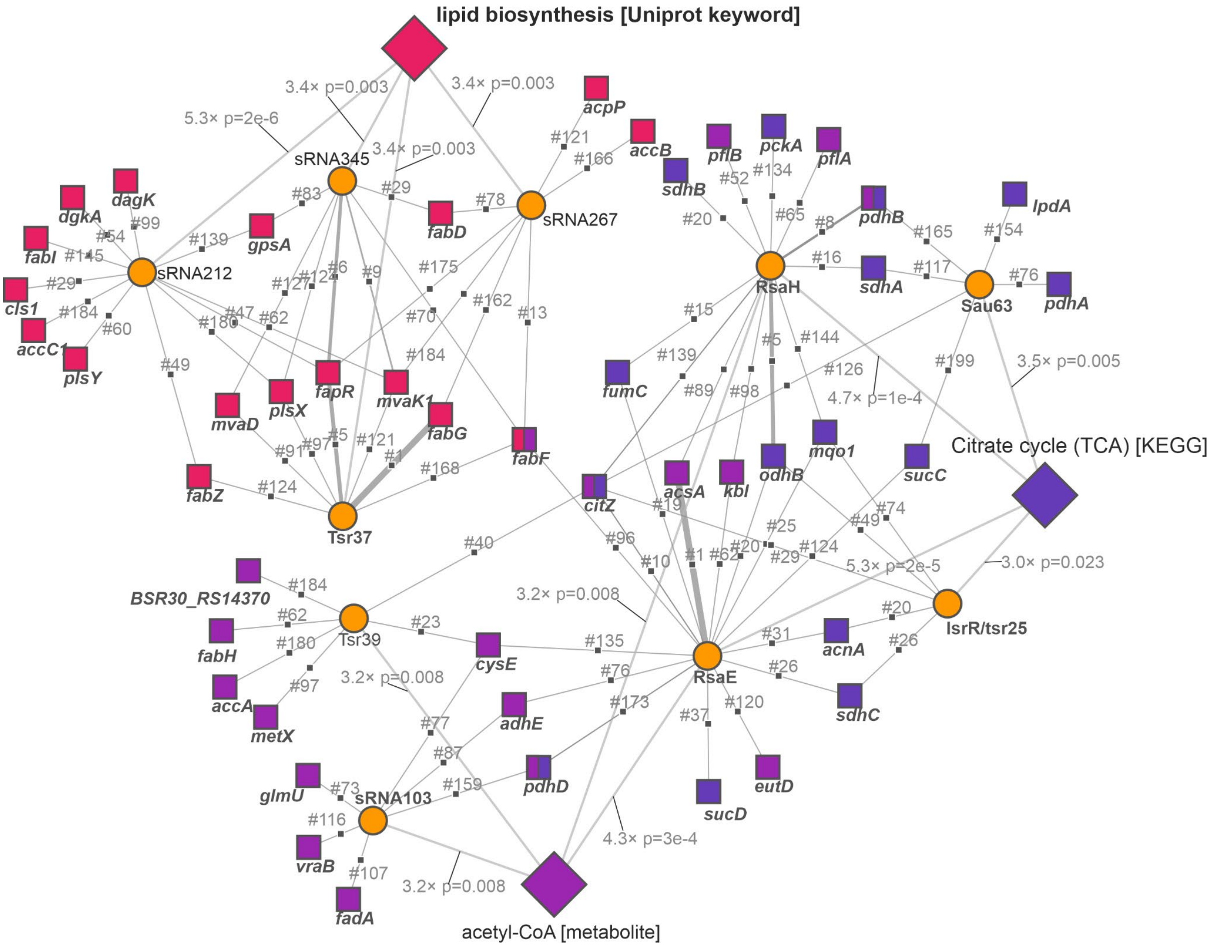


**Supplementary Figure S6.** Network generated with the aureoRNA network viewer tool visualizing examples for an explorative search based on enriched pathway search. Showing the top sRNAs with enriched predictions in lipid biosynthesis [UniProt keyword] (top 4), citrate cycle TCA [KEGG] (top 4) and usage of the acetyl-CoA metabolite (top 3). Edges from sRNA to predicted target show the prediction rank, edges from term to sRNA show the enrichment p-value.

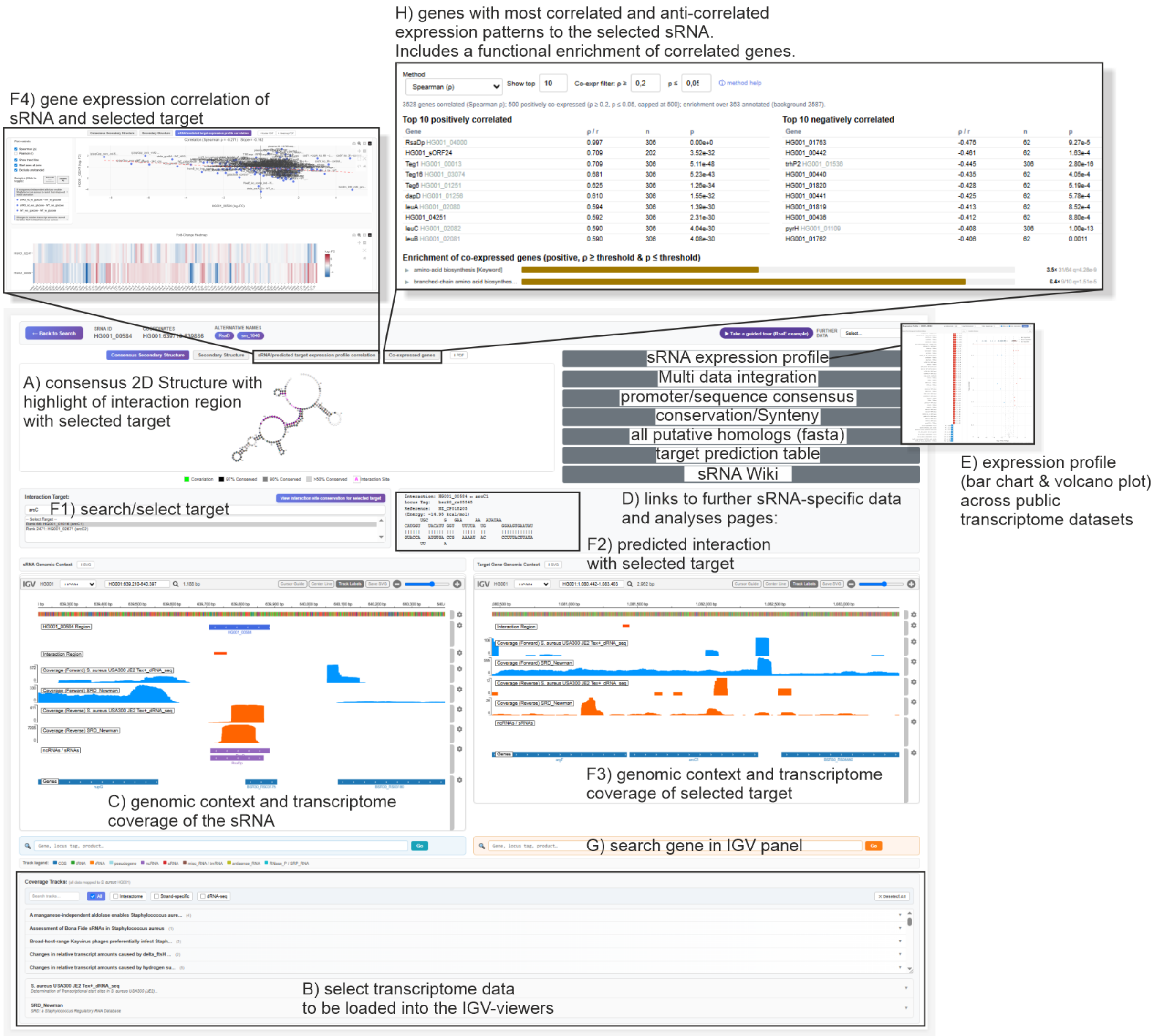


**Supplementary Figure S7**. Main sRNA landing page for RsaD. For a full explanation please use the “guided tour” on the website. **(A)** Window showing the consensus secondary structure prediction or single sequence secondary prediction. The bases involved in the interaction with the selected target (F1) are highlighted in purple (compare **Figure 9A**). **(B)** This panel allows searching and selecting different transcriptome datasets to display the coverage in the below IGV panels (C, F3). The study title and pubmed link as well as further information for the selected studies can be accessed via an expandable box below the selection panel. Strand specificity and source organism (e.g. NCTC8325/HG001 or USA300/JE2) for each dataset are given beside. By default a dRNAseq track focusing on actual primary transcription start sites (USA300/JE2)(Das *et al.*, 2016) and the transcriptome data from the SRD database study (Newman) (Sassi *et al.*, 2015) are selected. **(C)** IGV viewer (Robinson *et al.*, 2023) panel showing the genome context around the selected sRNA including the coverage of the selected transcriptome studies. **(D)** Links to further sRNA specific aureoRNA webpages. **(E)** A Popup window displaying the sRNA expression fold-change patterns as bar charts and volcano plots. **(F)**(F1) Search slot for target genes, by default the number one prediction is selected. For the selected target the predicted interaction is displayed (F2), as well as the target genome region and transcriptome coverage in the right IGV panel (F3). For both sRNA and target the predicted interaction region is indicated in the IGV panels as orange bars. (F4) By selecting the "sRNA/predicted target expression profile correlation" tab the fold-change expression correlation of sRNA and selected target is displayed as scatterplot and heatmap. **(G)** Search slot for any gene to be displayed in the IGV browser. **(H)** Display of the genes with most correlated and anti-correlated expression patterns to the selected sRNA. Spearman-, Pearson- or standard error weighted correlation can be selected.

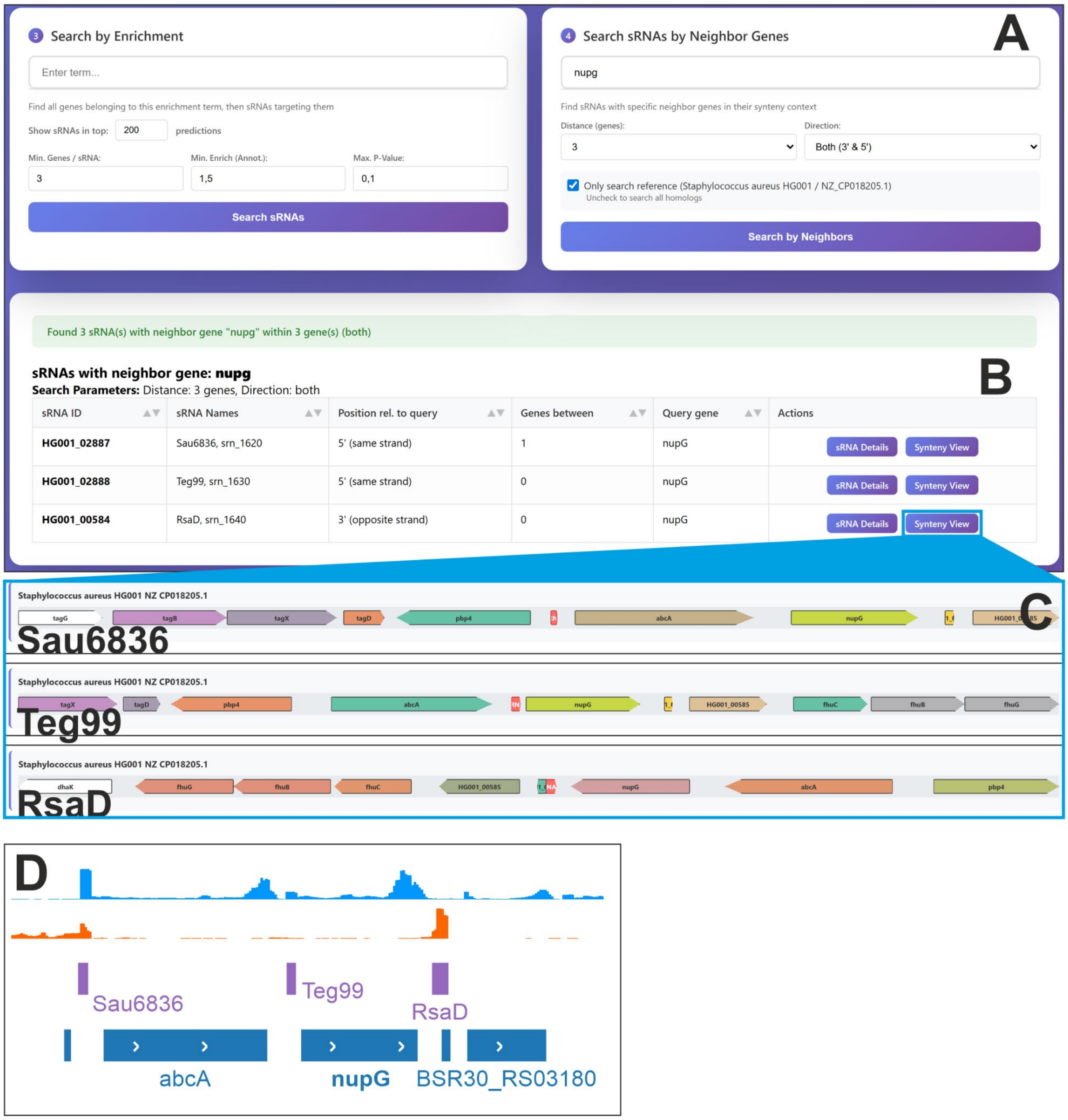


**Supplementary Figure S8.** Example for a genome neighborhood-centric search. **(A)** *nupG* was used as a query and all database sRNAs in 3 gene-distance 5' or 3' of the query gene are displayed. **(B)** The primary result is a table with sRNA ID and its orientation and distance relative to the query. In case of nupG two sRNAs, Sau6836 and Teg99, are located upstream on the same strand and one sRNA, RsaD, is located downstream on the opposite strand. **(C)** A click on the synteny view button directs to the sRNA specific synteny conservation page (**Supplementary Figure S9**). Here only a part of the view is shown. **(D)** On the sRNA landing page the synteny and the transcriptome coverage can be investigated in a IGV browser window.

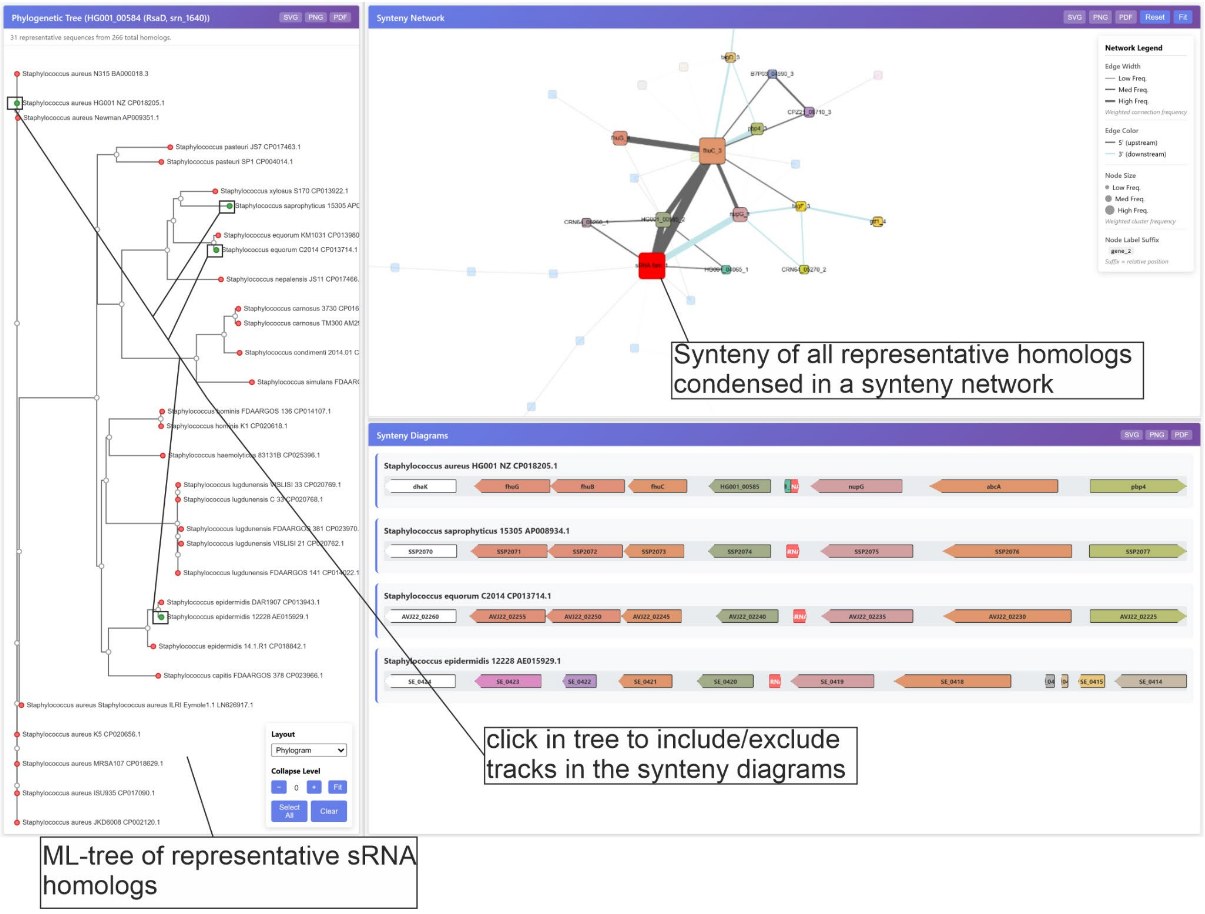


**Supplementary Figure S9.** Webpage to view the synteny and synteny conservation of sRNA homologs. For a full explanation please use the “guided tour” on the website.

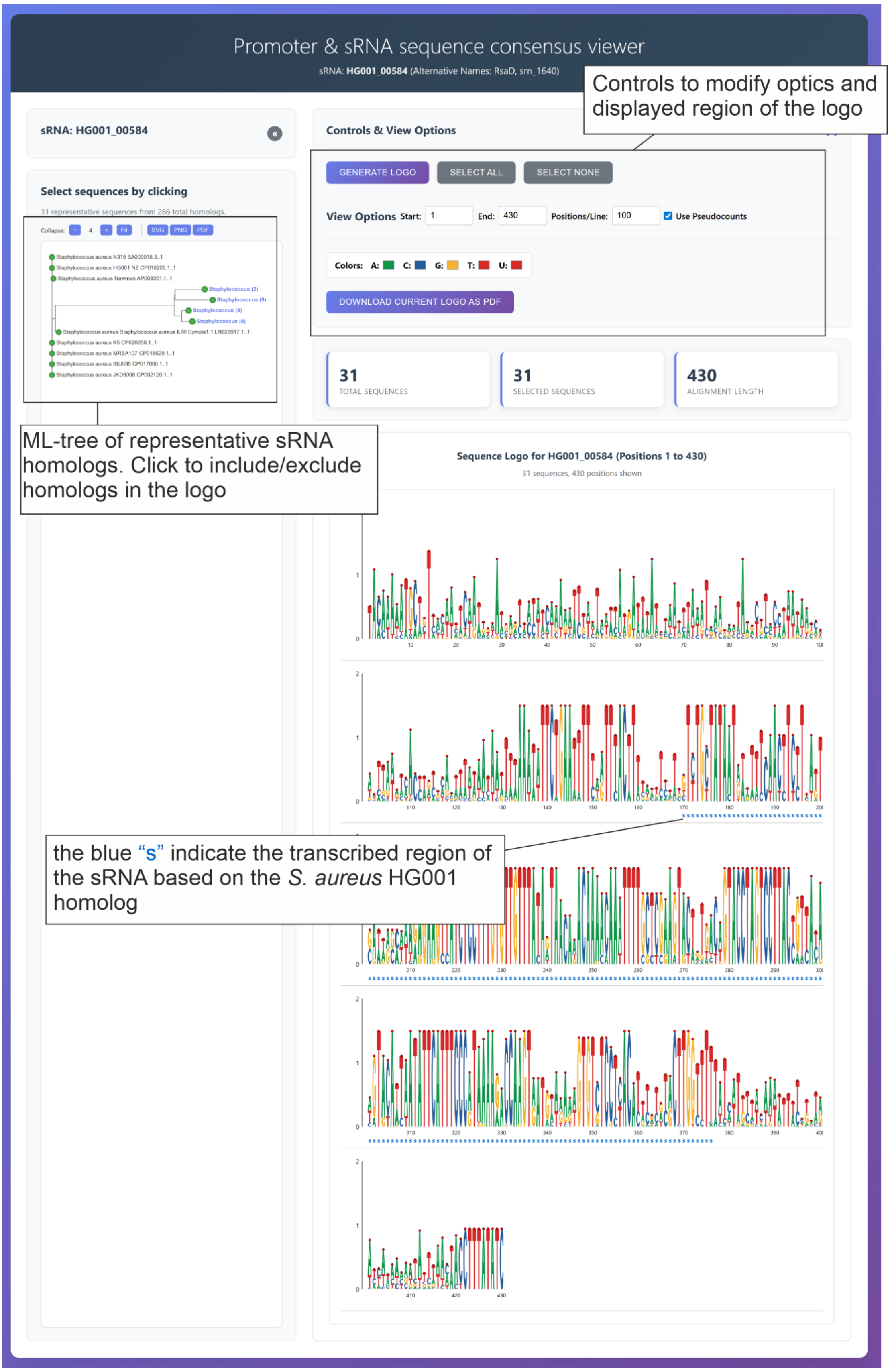

Promoter & sRNA sequence consensus viewer
sRNA: HG001_00584 (Alternative Names: RsaD, srn_1640)
Controls to modify optics and displayed region of the logo
sRNA: HG001_00584
Controls & View Options
GENERATE LOGO
SELECT ALL
SELECT NONE
View Options Start: 1 End: 430 Positions/Line: 100 Use Pseudocounts
Colors: A: C: G: T: U:
DOWNLOAD CURRENT LOGO AS PDF
Select sequences by clicking
31 representative sequences from 266 total homologs
31 TOTAL SEQUENCES
31 SELECTED SEQUENCES
430 ALIGNMENT LENGTH
ML-tree of representative sRNA homologs. Click to include/exclude homologs in the logo
Sequence Logo for HG001_00584 (Positions 1 to 430)
31 sequences, 430 positions shown
the blue "s" indicate the transcribed region of the sRNA based on the *S. aureus* HG001 homolog

**Supplementary Figure S10.** Webpage to view the sequence conservation of sRNA promoters and sRNA sequences. For a full explanation, use the “guided tour” on the website. Interactive sequence logo based on a pre-calculated multiple sequence alignment of the representative homologs including 200 nt upstream and 50 nt downstream of the transcribed sRNA sequences. At each position in the alignment, observed base counts are first adjusted with optional pseudocounts (prior = 1.0, effectively adding one observation of each base) to handle low-frequency bases, then modeled as a Dirichlet distribution with parameters $\alpha(b,i) = \text{count}(b,i) + \text{prior}$, from which the expected entropy $\bar{H}(i)$ is calculated using the digamma function: $\bar{H}(i) = -\Sigma[\alpha(b,i)/\Sigma\alpha \times \psi(1+\alpha(b,i))]/\Sigma\alpha + \psi(\Sigma\alpha+1)$, where $\psi$ is the digamma function accounting for uncertainty from finite sample sizes. The information content (height) at position i is then $R(i) = -\Sigma[p(b,i)\times\ln(p\_bg)] - \bar{H}(i)$ measured in bits (divided by ln(2)). Each letter's displayed height equals $p(b,i) \times R(i)$, creating a Bayesian-corrected visualization where conservation at positions with few sequences is appropriately down-weighted compared to well-sampled positions similar to webLogo (Crooks *et al.*, 2004).

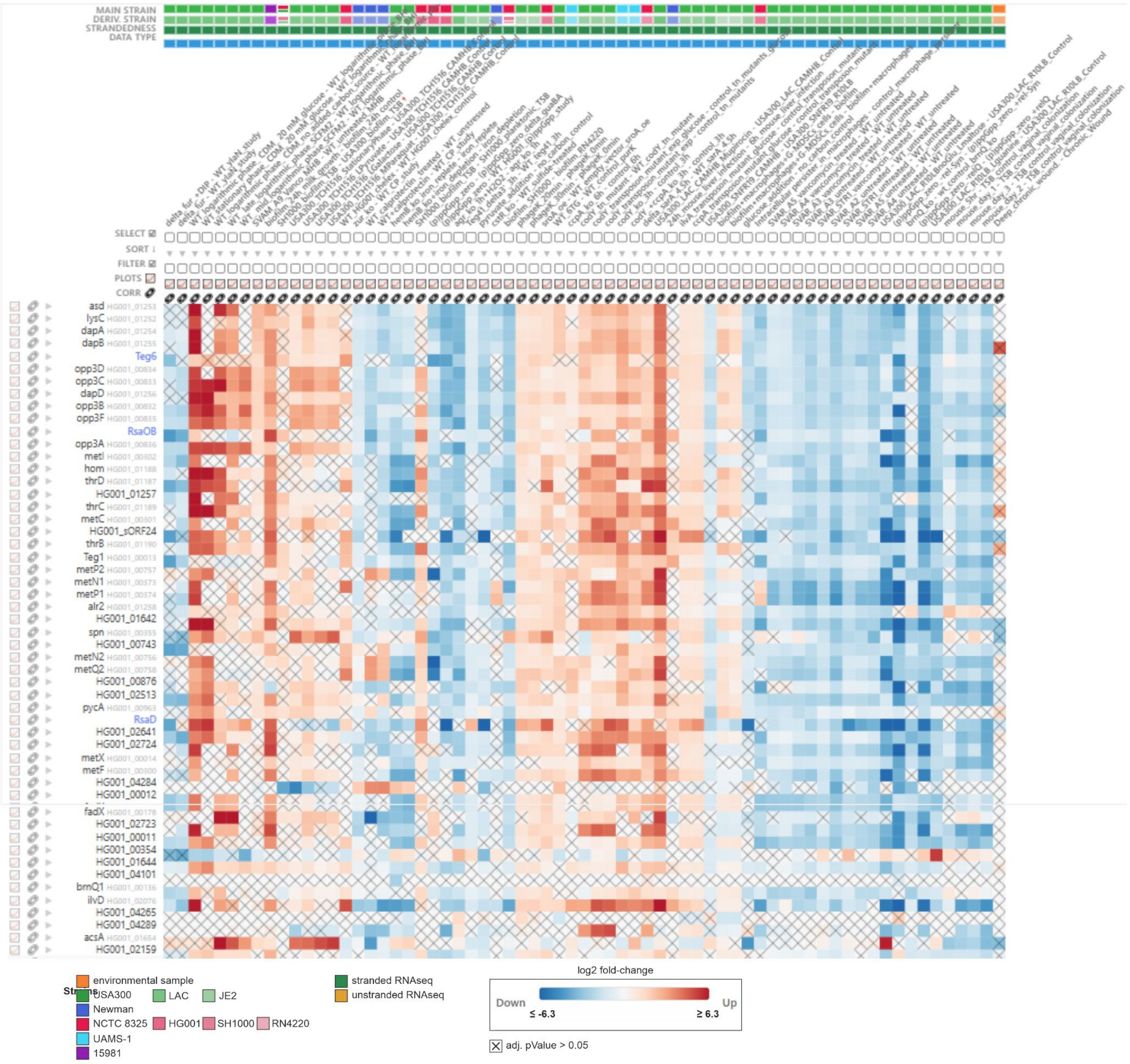


**Supplementary Figure S11.** The CodY iModulon module. The genes belonging to the module are shown, ordered by membership weight from high to low. Only fold-change columns in which the CodY iModulon activity changed by |Δ| ≥ 6 are displayed without a statistical significance threshold (RNAseq only). Cell colour gives the log2 fold change relative to the study-specific control (red, up; blue, down); crossed cells did not pass the significance threshold (i.e. adj. p-value > 0.05). The tracks above the heatmap indicate the main strain, the derivative strain (lighter shades of the main-strain color, e.g. Main strain NCTC 8325, derivative strain HG001). It's also indicated for each contrast column if the respective libraries were stranded (green box) or unstranded (yellow box).

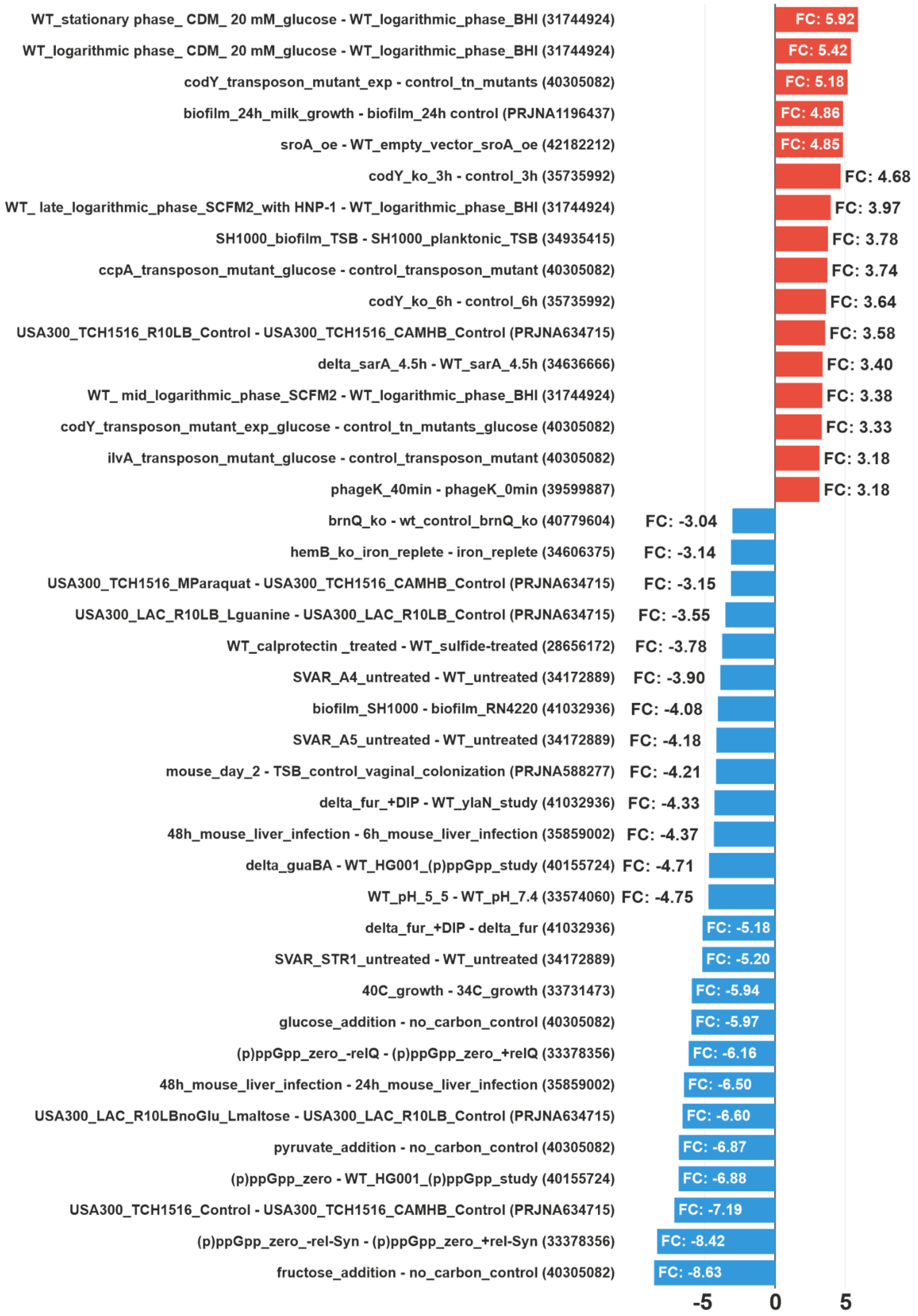


**Supplementary Figure S12.** Expression fold-change barchart for RsaD. All significant FCs with an absolute change ≥ 3 are shown. Available Pubmed IDs of the respective studies or the NCBI BioProject IDs are given in the brackets.

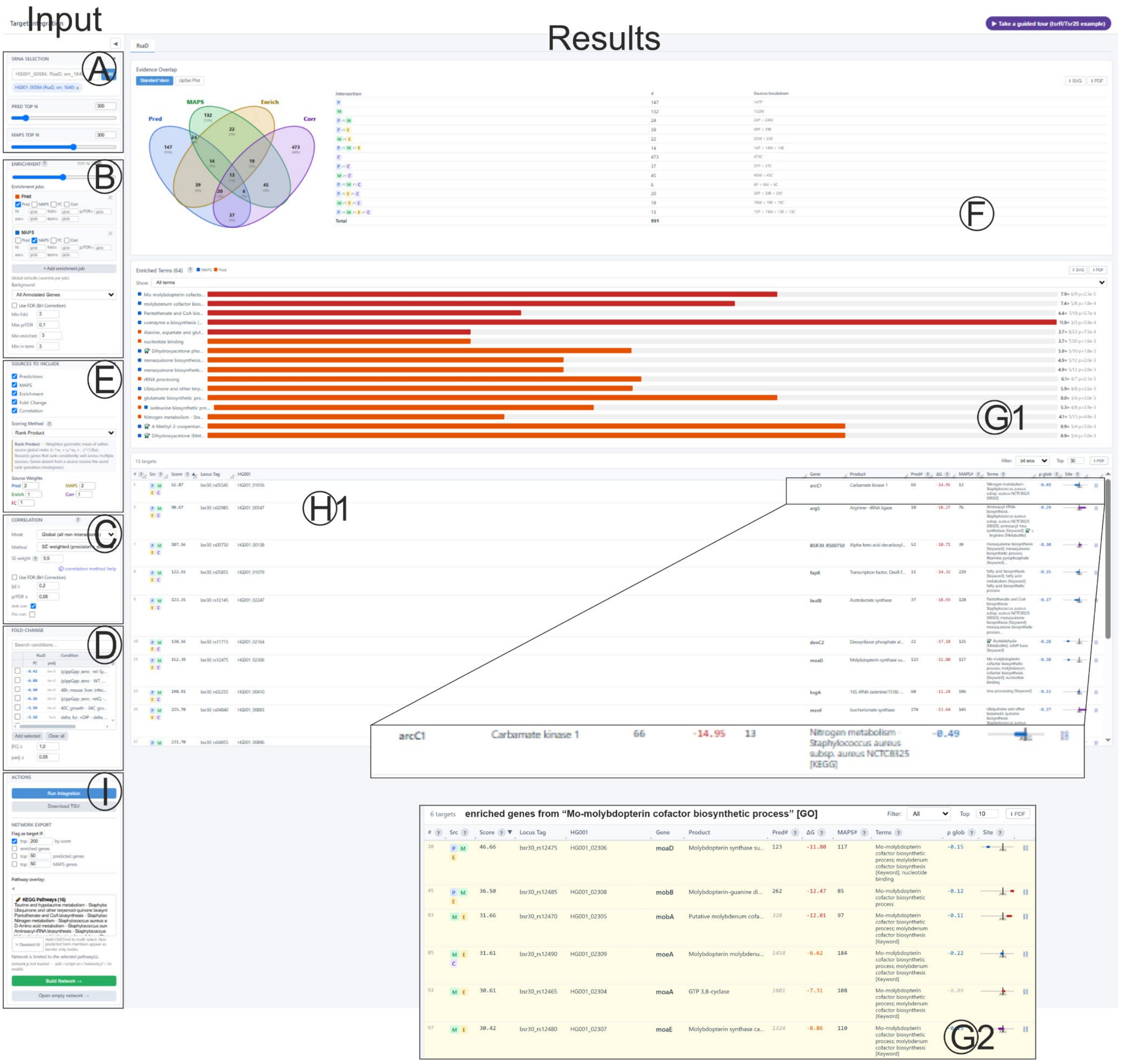


**Supplementary Figure S13.** aureoRNA “Multi data integration analysis” for RsaD. For a full explanation please use the “guided tour” on the website. The controls on the left side allow for sRNA, data and parameter selection. **(A)** Selection of the sRNA and rank thresholds for CopraRNA, MAPS (if available) and global threshold for the functional enrichment tool. **(B)** Functional enrichment tool. Multiple jobs for different datasets and dataset combinations can be added. Parameters (rank thresholds, p-Value thresholds, minimal genes in term and minimal genes in enrichment) can be adjusted independently for each job or via global thresholds. **(C)** Parameters for the gene expression correlation (Pearson, Spearman correlation or standard error weighted Pearson, global vs. for selected fold-changes only, absolute rho threshold for inclusion, p-Value threshold, only anti-correlated genes vs only correlated genes vs both). **(D)** Selection of gene expression fold-change data to be included into the analyses. The data are decreasingly ordered by the absolute fold-change of the selected sRNA. **(E)** Selection which of the up to five data sources should be combined for the final score. Four different score calculation methods are available (see methods). The weight of each data source can be manually adapted (default CoprarNA 2, MAPS 2, all other sources 1). The results of the analyses are shown in the right part of the view. **(F)** Data overlap shown as Venn-diagram or UpSet plot. **(G)**(G1) Results of the functional enrichment analysis. At the start of each bar one or more colored boxes indicate the data source the enrichment was based on. Bar length indicates the enrichment fold-change and the color the p-Value. G2) Clicking on a "term" bar displays only the genes belonging to the term in the below table. **(H)**(H1) Result table ranked by the combination score. Here, the weighted rank sum. The full list is available as Supplement (RsaD_full_target_list). The table can be downloaded as a tsv file and contains all relevant information (H2: prediction rank, MAPS rank, predicted interaction energy, enriched terms, correlation rho, graphical representation of interaction site relative to the start codon, link to interaction site conservation page, fold-changes of selected gene expression datasets). **(I)** The enriched genes and pathways can be visualized with the aureoRNA pathway tool (**Supplementary Figure S6**).

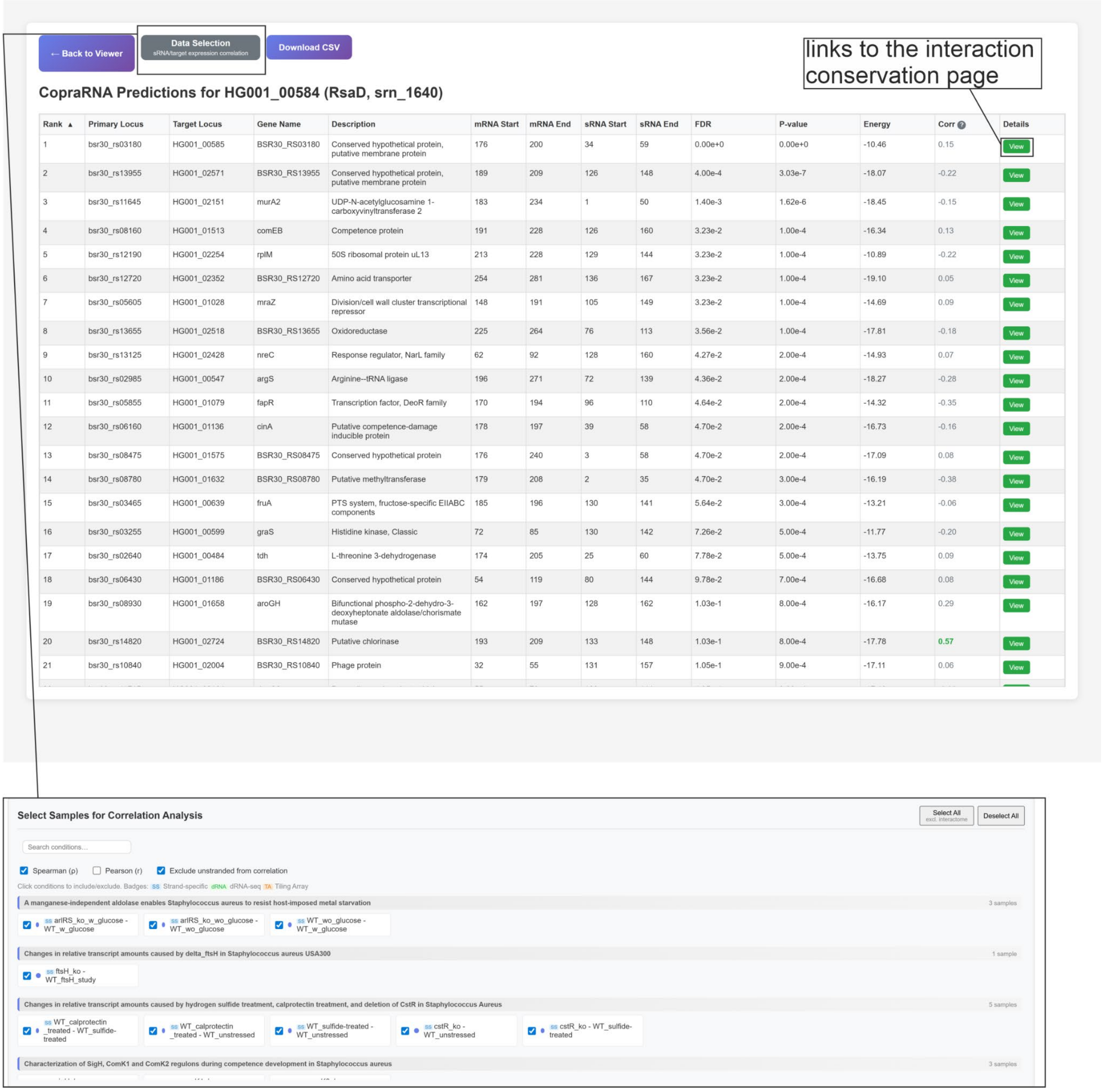


**Supplementary Figure S14.** CopraRNA target prediction table for RsaD. Primary CopraRNA results with associated global correlation (default: Spearman correlation) from gene expression data. For the top 300 predictions a link to a dedicated interaction conservation webpage is given. The data selection button can be used to specify the fold-changes to be included in the correlation score.

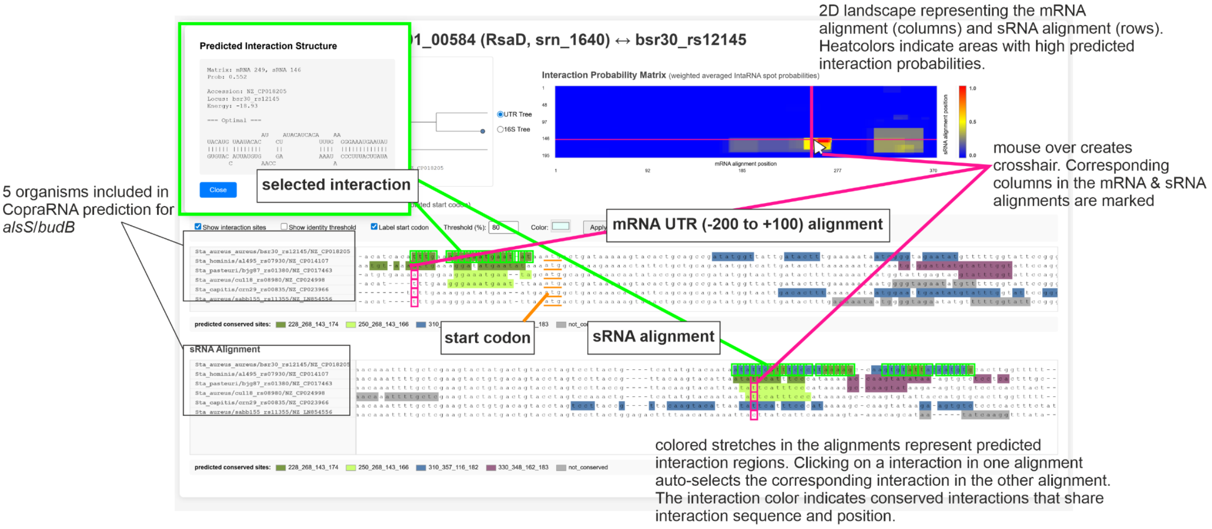


**Supplementary Figure S15.** Individual interaction conservation page for a specific sRNA/target pair. For a full explanation, please use the “guided tour” on the website.

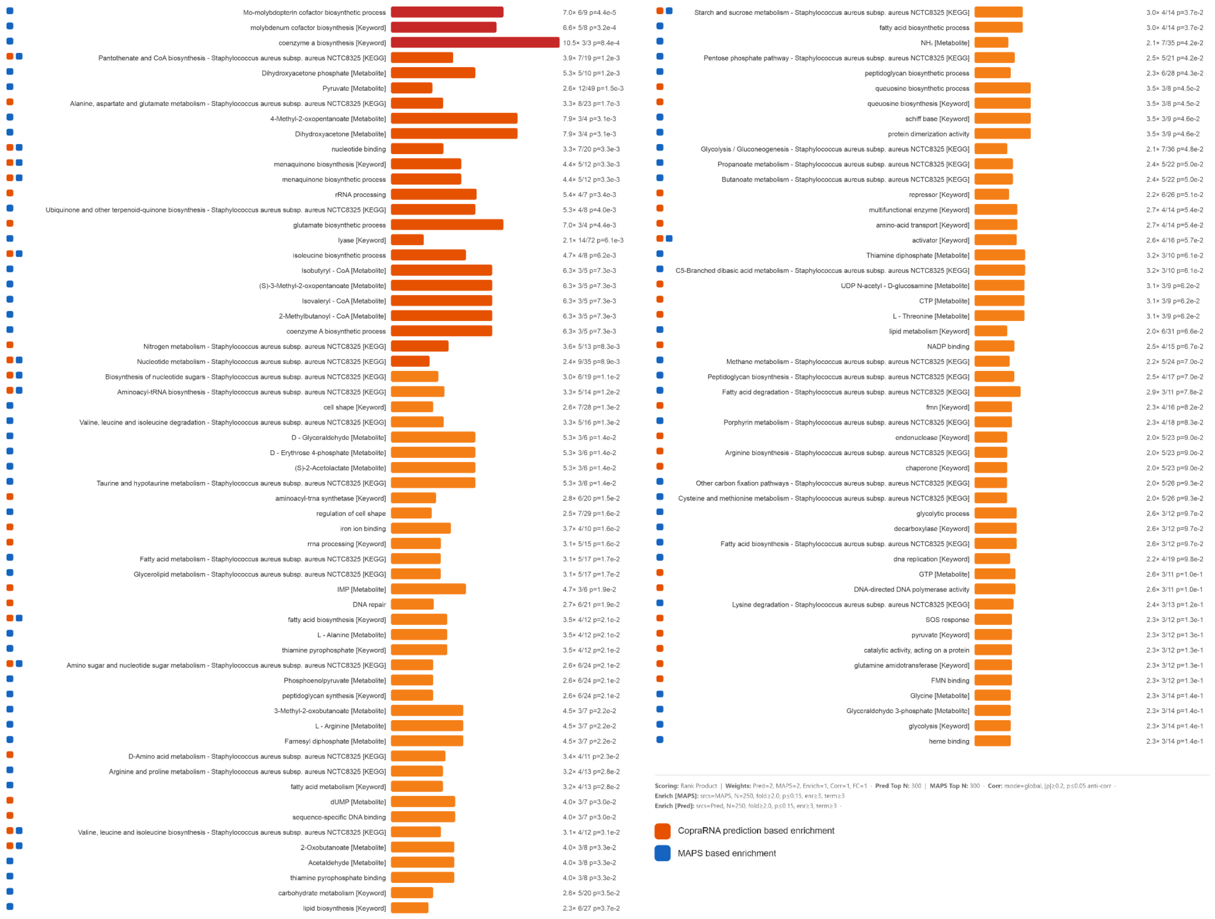

**Supplementary Figure S16.** Full RsaD functional enrichment from the aureoRNA website. The left square indicates the source data for the enriched term (■ CopraRNA prediction, ■ MAPS). The orange bar is relative to the enrichment fold change and at the right enrichment fold change, enriched genes/all genes in term and the enrichment p-Value are shown. This figure extends **Figure 8C**.

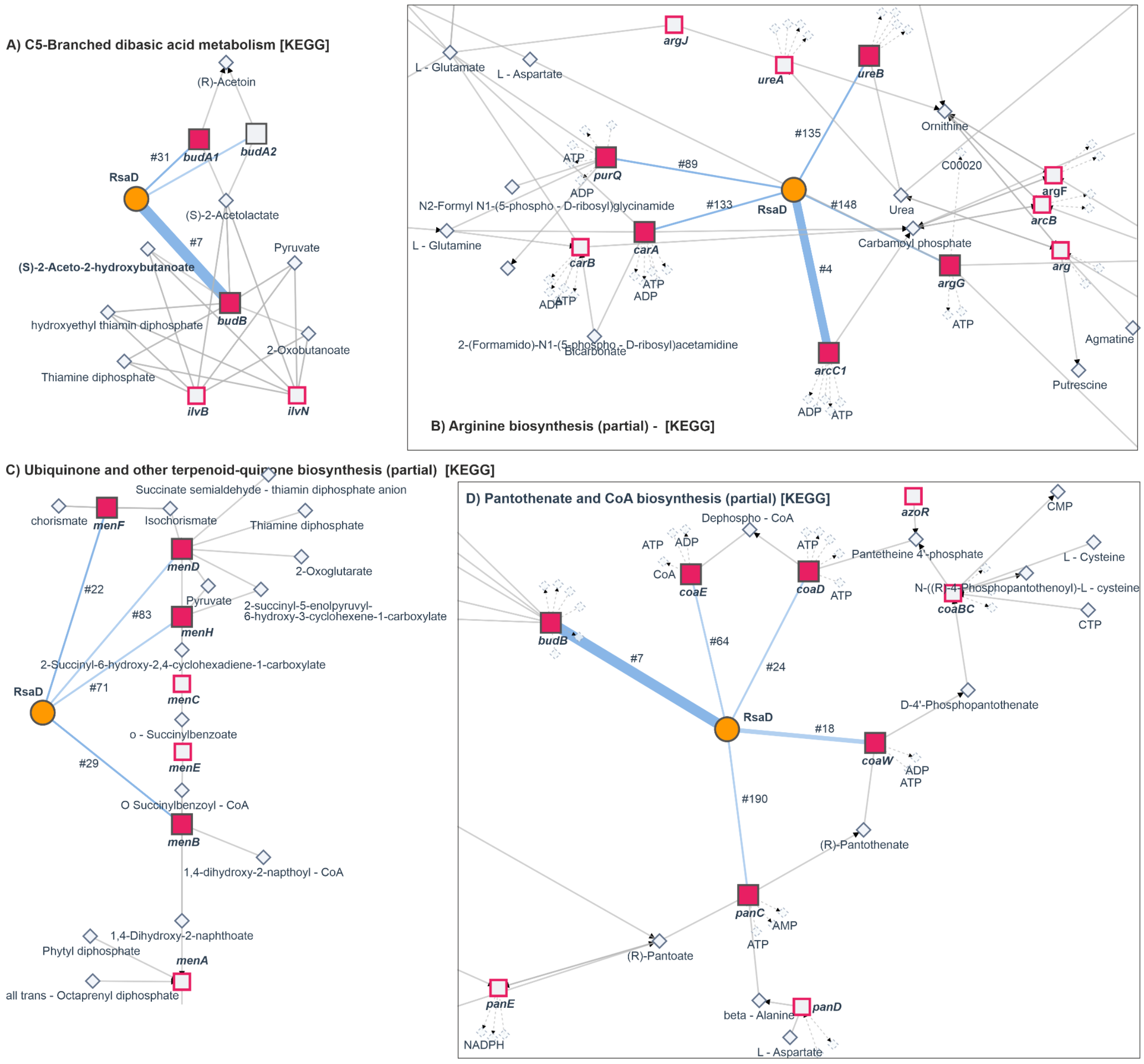


**Supplementary Figure S17.** Proposed targets after the aureoRNA data integration can be mapped to their enriched terms or metabolic pathways. Predicted targets within the pathways are indicated by filled red boxes and edges to the sRNA.

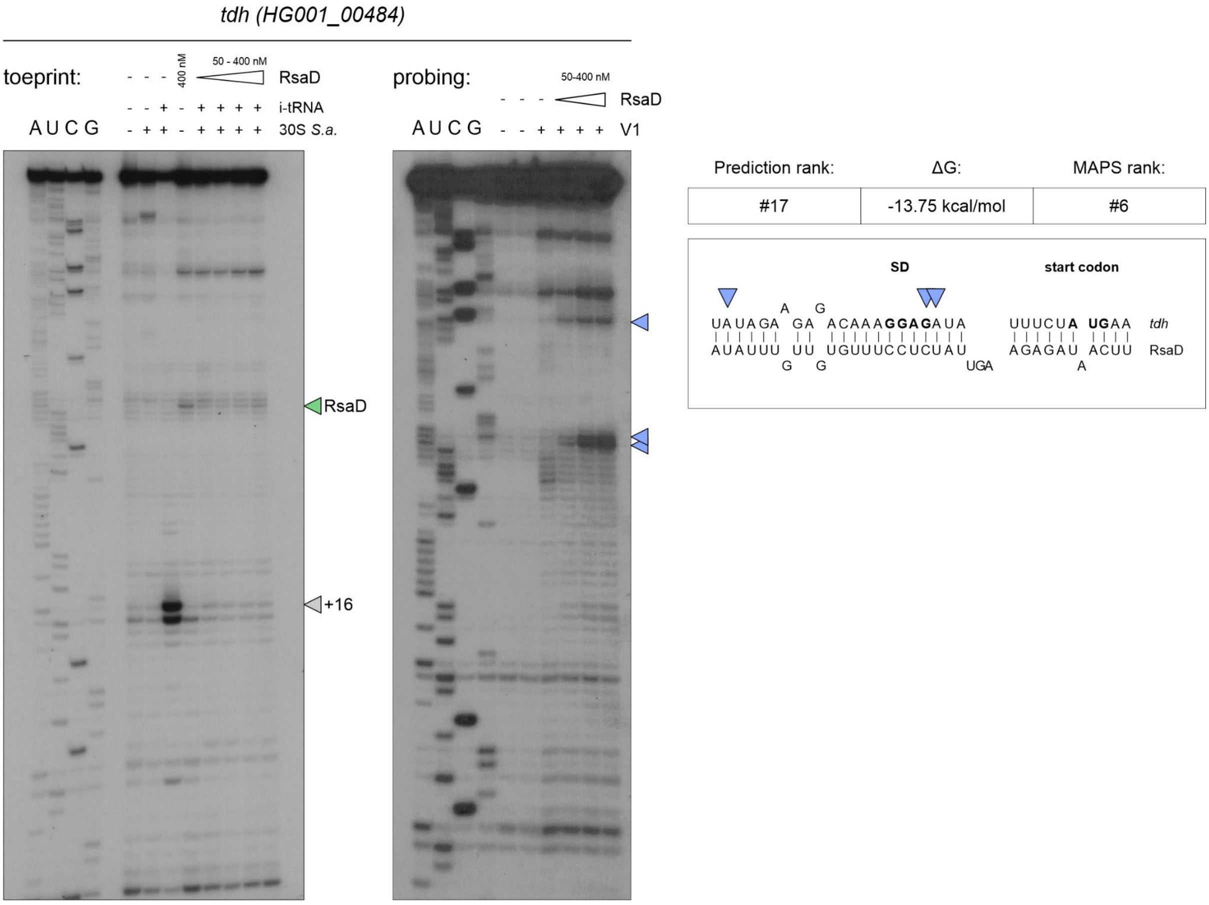


**Supplementary Figure S18. Toeprinting and RNase V1 probing of the *tdh* mRNA in the absence or presence of RsaD.** Full gel images corresponding to **Figure 10A** and **B**. **Left**: Toeprinting analysis of *tdh* (HG001_00484) mRNA with *S. aureus* 30S ribosomal subunits and initiator tRNA (i-tRNA). Experimental lanes, numbered from left to right after the sequencing ladders, contain mRNA alone (lane 1), mRNA with 30S subunits (lane 2), mRNA with 30S subunits and i-tRNA (lane 3), mRNA with RsaD alone (400 nM; lane 4), or mRNA with 30S subunits, i-tRNA and increasing RsaD concentrations (50–400 nM; lanes 5–8). Open arrowhead indicates toeprint at position +16 relative to the first nucleotide of the start codon, and green arrowhead marks an RsaD-associated reverse-transcription stop. **Right**: RNase V1 probing of *tdh* mRNA in the absence or presence of increasing RsaD concentrations (50–400 nM). The first two lanes contain controls without RNase V1, comprising mRNA alone or mRNA with RsaD, respectively. The remaining lanes contain RNase V1 without RsaD or with increasing RsaD concentrations. A, U, C and G denote sequencing ladders. Blue arrowheads mark RNase V1 cleavages resulting from RsaD binding; they are given on the autoradiography and on the *tdh* mRNA sequence paired with RsaD (shown in the insert).

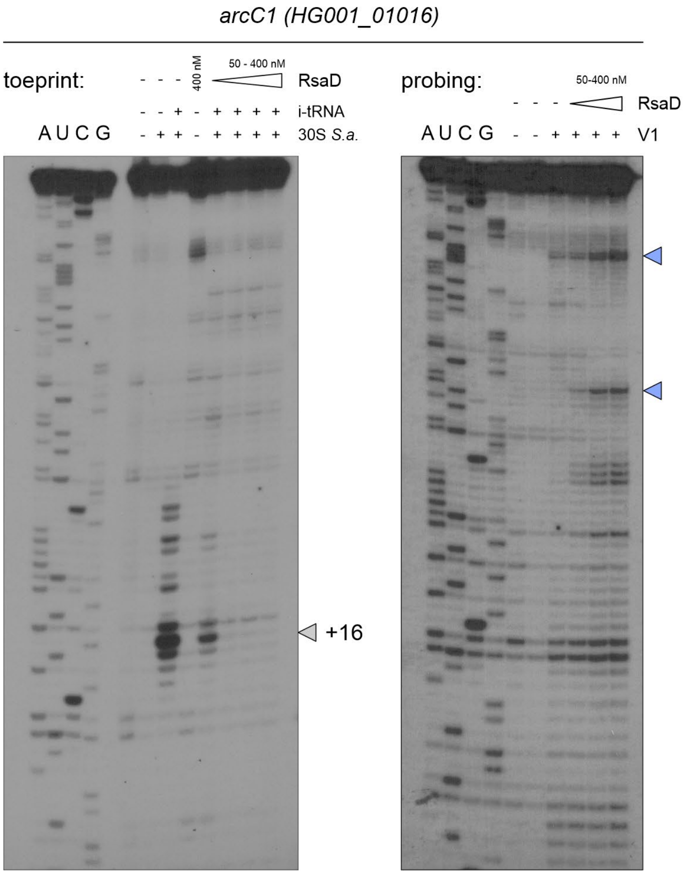


| Prediction rank: | ΔG: | MAPS rank: |
|---|---|---|
| #66 | -14.95 kcal/mol | #13 |

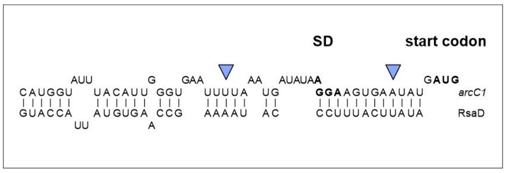


**Supplementary Figure S19. Toeprinting and RNase V1 probing of the *arcC1* mRNA in the absence or presence of RsaD.** Full gel images corresponding to Fig. 10G and H. **Left**: Toeprinting analysis of *arcC1* (HG001_01016) mRNA with *S. aureus* 30S ribosomal subunits and initiator tRNA (i-tRNA). Experimental lanes, numbered from left to right after the sequencing ladders, contain mRNA alone (lane 1), mRNA with 30S subunits (lane 2), mRNA with 30S subunits and i-tRNA (lane 3), mRNA with RsaD alone (400 nM; lane 4), or mRNA with 30S subunits, i-tRNA and increasing RsaD concentrations (50–400 nM; lanes 5–8). Open arrowheads indicate toeprint at position +16 relative to the first nucleotide of the start codon. **Right**: RNase V1 probing of *arcC1* mRNA in the absence or presence of increasing RsaD concentrations (50–400 nM). The first two lanes contain controls without RNase V1, comprising mRNA alone or mRNA with RsaD, respectively. The remaining lanes contain RNase V1 without RsaD or with increasing RsaD concentrations. A, U, C and G denote sequencing ladders. Blue arrowheads mark RNase V1 cleavages resulting from RsaD binding; they are given on the autoradiography and on the *arcC1* mRNA sequence paired with RsaD (shown in the insert).

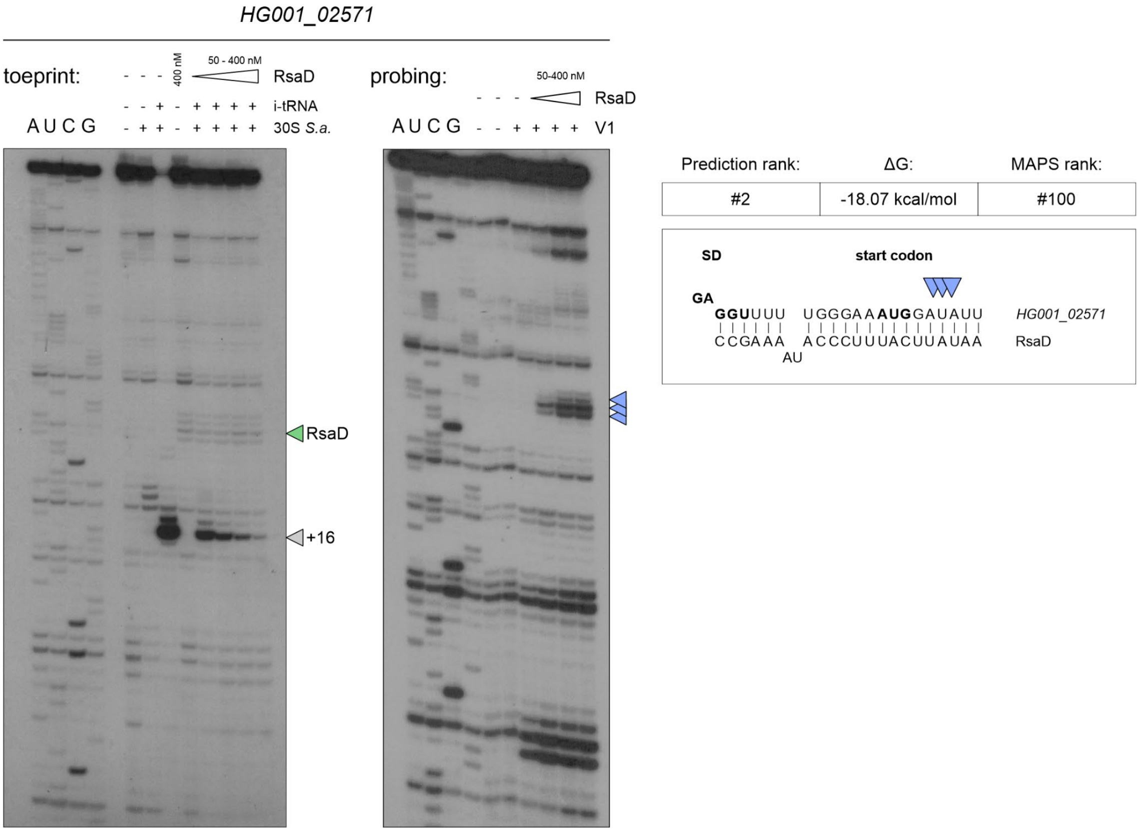


**Supplementary Figure S20. Toeprinting and RNase V1 probing of the HG001_02571 mRNA in the absence or presence of RsaD.** Full gel images corresponding to Fig. 10D and E. **Left**: Toeprinting analysis of HG001_02571 mRNA with ***S. aureus*** 30S ribosomal subunits and initiator tRNA (i-tRNA). Experimental lanes, numbered from left to right after the sequencing ladders, contain mRNA alone (lane 1), mRNA with 30S subunits (lane 2), mRNA with 30S subunits and i-tRNA (lane 3), mRNA with RsaD alone (400 nM; lane 4), or mRNA with 30S subunits, i-tRNA and increasing RsaD concentrations (50–400 nM; lanes 5–8). Open arrowheads indicate toeprint at position +16 relative to the first nucleotide of the start codon. The green arrowhead marks an RsaD-associated reverse-transcription stop. **Right**: RNase V1 probing of HG001_02571 mRNA in the absence or presence of increasing RsaD concentrations (50–400 nM). The first two lanes contain controls without RNase V1, comprising mRNA alone or mRNA with RsaD, respectively. The remaining lanes contain RNase V1 without RsaD or with increasing RsaD concentrations. A, U, C and G denote sequencing ladders. Blue arrowheads mark RNase V1 cleavages resulting from RsaD binding; they are given on the autoradiography and on the *arcC1* mRNA sequence paired with RsaD (shown in the insert).

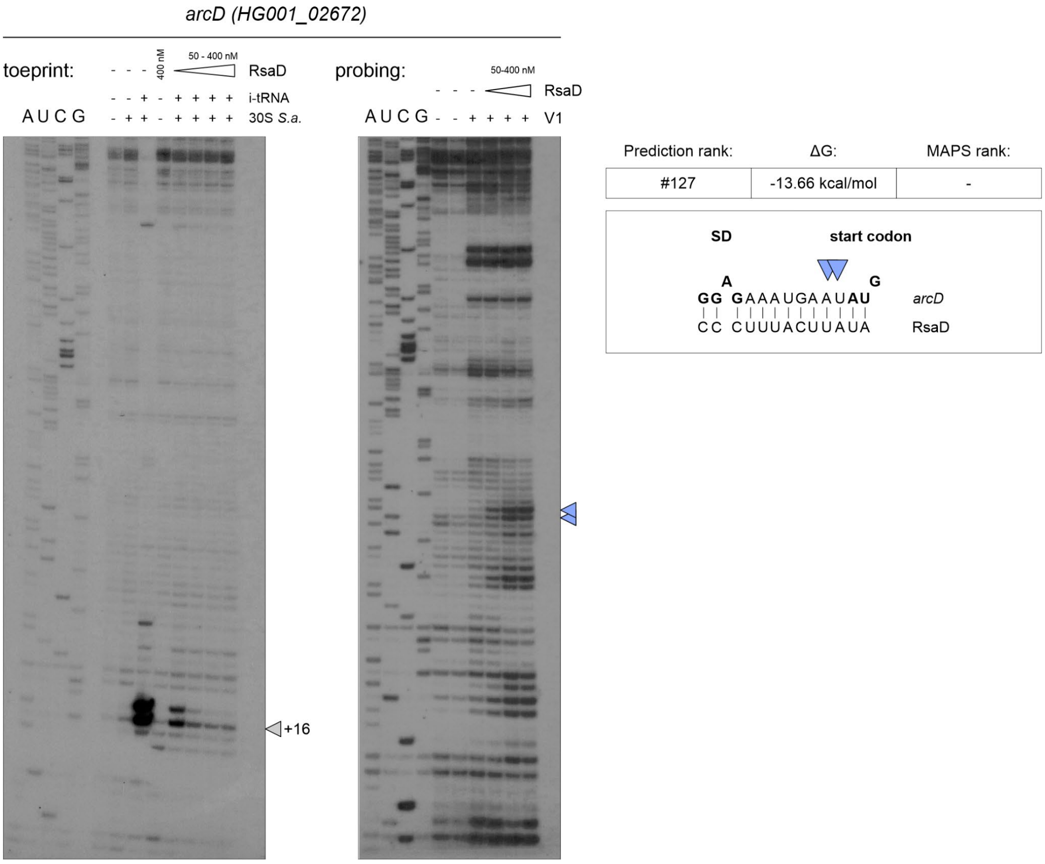


**Supplementary Figure S21. Toeprinting and RNase V1 probing of the *arcD* mRNA in the absence or presence of RsaD.** Full gel images corresponding to **Figure 10J** and **K**. **Left**: Toeprinting analysis of *arcD* (HG001_02672) mRNA with *S. aureus* 30S ribosomal subunits and initiator tRNA (i-tRNA). Experimental lanes, numbered from left to right after the sequencing ladders, contain mRNA alone (lane 1), mRNA with 30S subunits (lane 2), mRNA with 30S subunits and i-tRNA (lane 3), mRNA with RsaD alone (400 nM; lane 4), or mRNA with 30S subunits, i-tRNA and increasing RsaD concentrations (50–400 nM; lanes 5–8). Open arrowhead indicates toeprint at position +16 relative to the first nucleotide of the start codon. **Right**: RNase V1 probing of *arcD* mRNA in the absence or presence of increasing RsaD concentrations (50–400 nM). The first two experimental lanes contain controls without RNase V1, comprising mRNA alone or mRNA with RsaD, respectively. The remaining lanes contain RNase V1 without RsaD or with increasing RsaD concentrations. A, U, C and G denote sequencing ladders. Blue arrowheads mark RNase V1 cleavages resulting from RsaD binding; they are given on the autoradiography and on the *arcD* mRNA sequence paired with RsaD (shown in the insert).

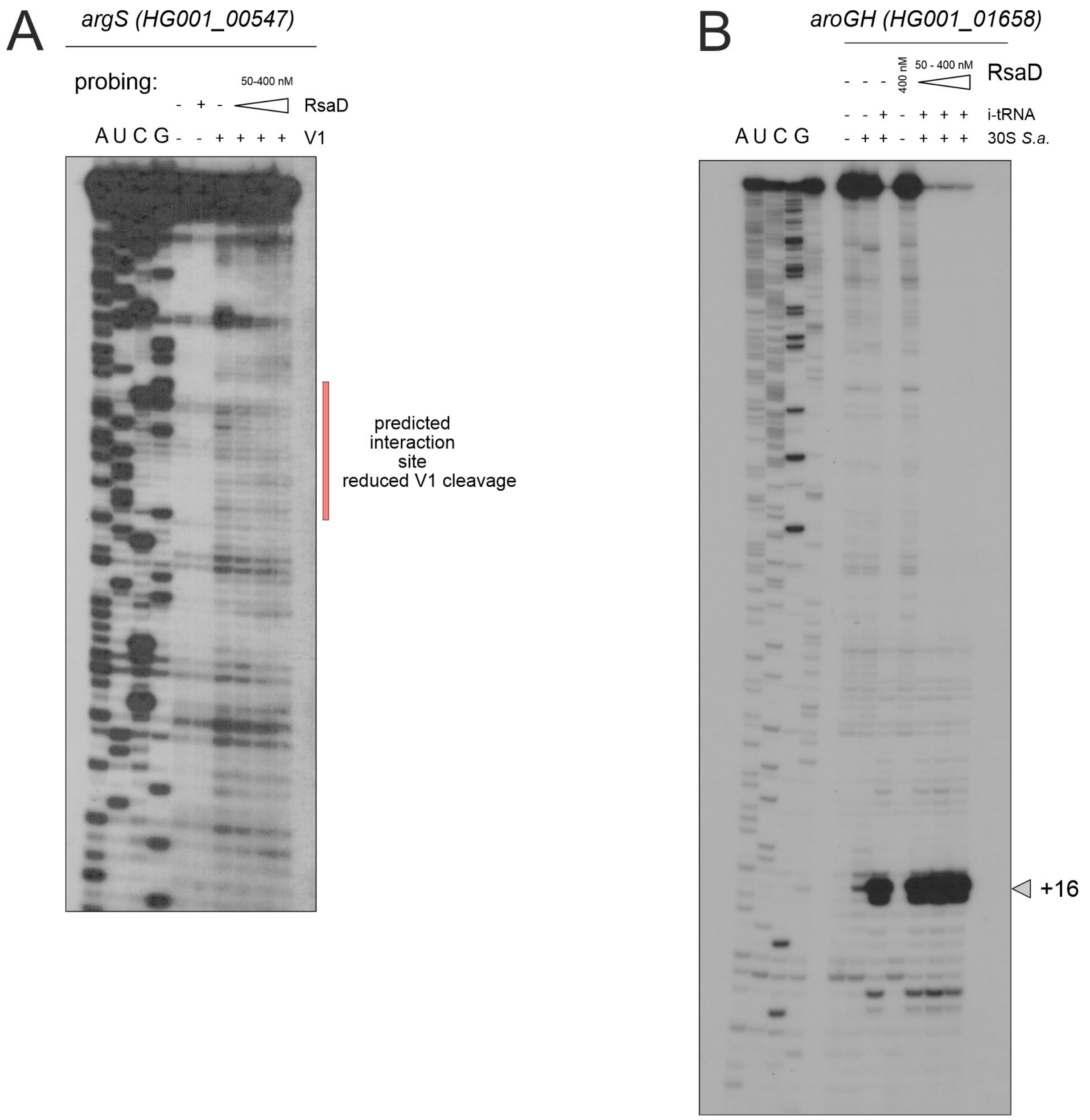


**Supplementary Figure S22. RNase V1 probing of *argS* mRNA and toeprinting analysis of *aroGH* mRNA in the absence or presence of RsaD.** (**A**) RNase V1 probing of *argS* (HG001_00547) mRNA in the absence or presence of increasing RsaD concentrations (50-400 nM). Experimental lanes, numbered from left to right after the sequencing ladders, contain mRNA alone (lane 1), mRNA with RsaD without RNase V1 (lane 2), mRNA with RNase V1 without RsaD (lane 3), or mRNA with RNase V1 and increasing RsaD concentrations (lanes 4-6). The red vertical bar indicates the predicted RsaD interaction site, within which no RsaD-dependent RNase V1 cleavage was detected. (**B**) Toeprinting analysis of *aroGH* (HG001_01658) mRNA with *S.*

*aureus* 30S ribosomal subunits and initiator tRNA (i-tRNA), in the absence or presence of RsaD. Experimental lanes, numbered as in A, contain mRNA alone (lane 1), mRNA with 30S subunits (lane 2), mRNA with 30S subunits and i-tRNA (lane 3), mRNA with RsaD alone (400 nM; lane 4), or mRNA with 30S subunits, i-tRNA and increasing RsaD concentrations (50-400 nM; lanes 5-7). The arrowhead indicates the toeprint at position +16 relative to the first nucleotide of the start codon. No RsaD-dependent reduction in the toeprint signal was observed under the tested conditions. Triangles indicate increasing RsaD concentrations. A, U, C and G denote sequencing ladders.
